# Temperature-dependent local conformations and conformational distributions of cyanine dimer labeled single-stranded – double-stranded DNA junctions by 2D fluorescence spectroscopy

**DOI:** 10.1101/2021.10.21.465365

**Authors:** Dylan Heussman, Justin Kittell, Peter H. von Hippel, Andrew H. Marcus

## Abstract

DNA replication, and the related processes of genome expression, require binding, assembly, and function of protein complexes at and near single-stranded (ss) – double-stranded (ds) DNA junctions. These central protein-DNA interactions are likely influenced by thermally induced conformational fluctuations of the DNA scaffold across an unknown distribution of functionally relevant states to provide regulatory proteins access to properly conformed DNA binding sites. Thus, characterizing the nature of conformational fluctuations and the associated structural disorder at ss-dsDNA junctions is likely critical for understanding the molecular mechanisms of these central biological processes. Here we describe spectroscopic studies of model ss-dsDNA fork constructs that contain dimers of ‘internally labeled’ cyanine (iCy3) chromophore probes that have been rigidly inserted within the sugar-phosphate backbones of the DNA strands. Our combined analyses of absorbance, circular dichroism (CD) and two-dimensional fluorescence spectroscopy (2DFS) permit us to characterize the local conformational parameters and conformational distributions. We find that the DNA sugar-phosphate backbones undergo abrupt successive changes in their local conformations – initially from a right-handed and ordered DNA state to a disordered splayed-open structure and then to a disordered left-handed conformation – as the dimer probes are moved across the ss-dsDNA junction. Our results suggest that the sugar-phosphate backbones at and near ss-dsDNA junctions adopt specific position-dependent local conformations and exhibit varying extents of conformational disorder that deviate widely from the Watson-Crick structure. We suggest that some of these conformations are likely to function as secondary-structure motifs for interaction with protein complexes that bind to and assemble at these sites.

## 1. Introduction

The Watson-Crick (W-C) B-form double-helix (1) is the most stable of the myriad structures that DNA can (and must) adopt in order to function both as a template for gene expression and as a vehicle for transmitting heredity. Under physiological conditions, double-stranded (ds) DNA exists primarily as a narrow, Boltzmann-weighted distribution of base-sequence-dependent conformations, for which the W-C structure represents an approximate free energy minimum. The molecular interactions that stabilize dsDNA includes cooperative stacking of adjacent nucleotide (nt) bases, internal strain that stacking induces in the sugar-phosphate backbones, W-C hydrogen bonding between opposite complementary strands, intra- and inter-chain repulsion between adjacent backbone phosphates, counterion condensation and orientation of polar water molecules within the nearest solvation layers at exposed DNA surfaces. All these interactions are subject to thermally-induced fluctuations (i.e., DNA ‘breathing’), which may lead to local segments adopting transiently unstable conformations over time scales spanning tens-of microseconds to several seconds (2). For example, on sub-second time scales and depending on temperature relative to the overall melting temperature of the DNA duplex, the interior of local AT-rich regions of dsDNA may become exposed to the surrounding aqueous environment by spontaneously disrupting the W-C structure and forming open ‘bubble-like’ conformations. On the time scale of multiple seconds, segments of dsDNA may undergo higher order sequence-dependent distortions of the local conformation, such as the formation of a local ‘kink,’ ‘bend,’ or ‘twist.’

The spontaneous formation of an unstable local dsDNA conformation is very likely a key initial step in the assembly mechanisms of complexes of gene regulatory proteins that recognize and bind to specific nt base sequences. In contrast, protein-DNA assembly mechanisms that occur largely independently of specific nt base sequences must utilize the ability of the protein or protein complex to recognize certain secondary-structure motifs that can be adopted by the sugar-phosphate backbones of the DNA (3). For example, the assembly of DNA replication complexes involves the preferential binding of proteins to single-stranded (ss) – dsDNA forks and junctions (2). In principle, various types of DNA breathing near ss-dsDNA junctions may facilitate the interconversion between various unstable conformations, of which one or more may be expected to resemble that of the DNA framework within a stable protein-DNA complex. Such an unstable conformational species may serve as a ‘transition state’ for the interaction of DNA binding sites with replication proteins.

In this work, we present spectroscopic studies of the distributions of structural parameters that characterize the local conformations of the sugar-phosphate backbones at and near ss-dsDNA fork junctions. These experiments use DNA constructs in which the carbocyanine dye Cy3 has been covalently attached as a dimer pair within the sugar-phosphate backbones at specific positions relative to the ss-ds DNA junction (see Fig. 1). The Cy3 chromophore is often used as a fluorescent marker for DNA sequencing and other biotechnological applications due to its relatively high absorption cross-section and favorable fluorescence quantum yield (4). The Cy3 chromophore consists of a conjugated trimethine bridge that cojoins two indole-like substituents (see Fig. 1A). The lowest energy *π* → *π*\* electronic transition between ground state *g* and excited state *e* occurs when the molecule is in its *all-trans* ground state configuration.

**Figure 1.**
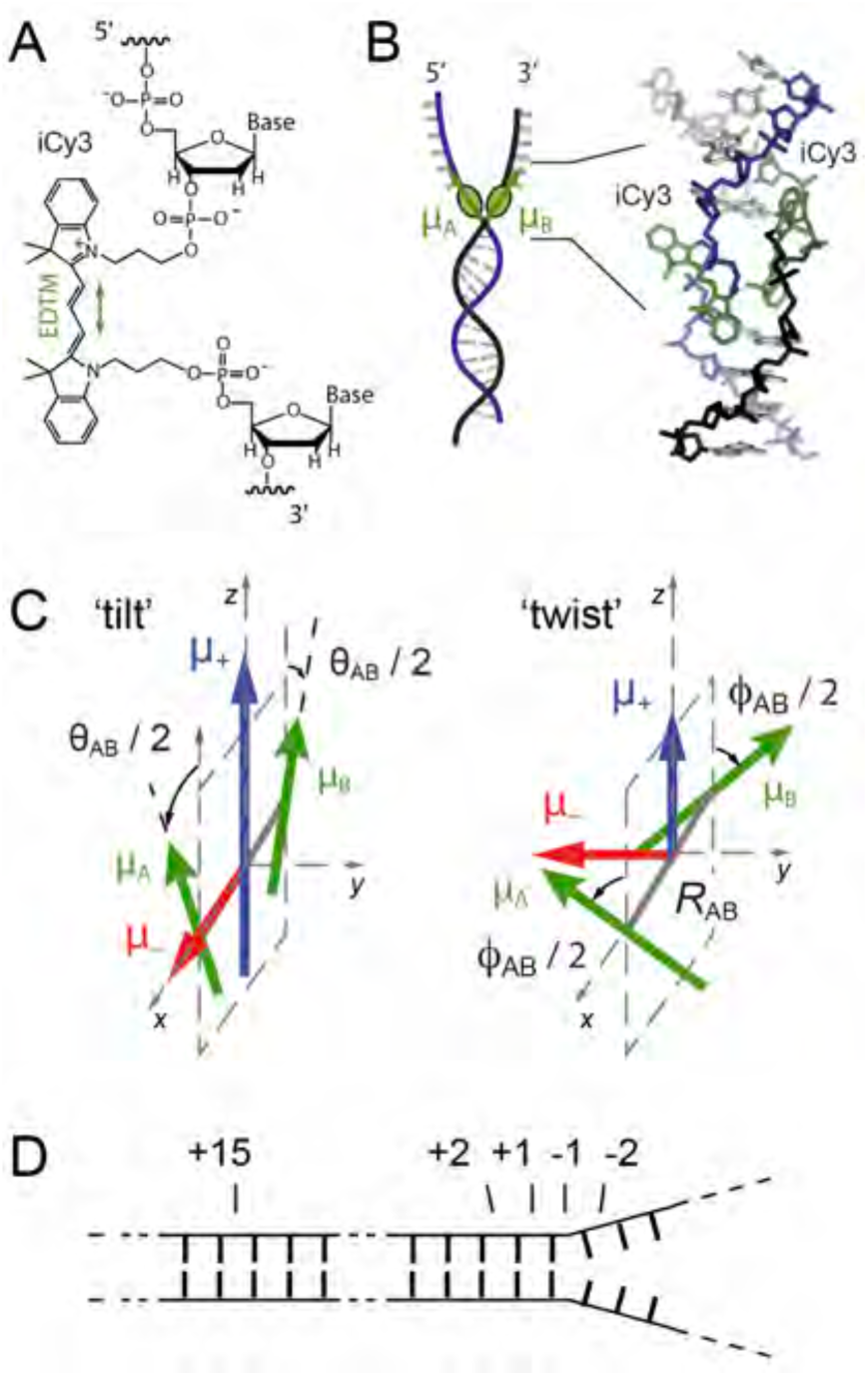
Labeling chemistry and nomenclature of the internal iCy3 dimer probes positioned within the sugar-phosphate backbones of model ss-dsDNA fork constructs. (***A***) The Lewis structure of the iCy3 chromophore is shown with its 3’ and 5’ linkages to the sugar-phosphate backbone of a local segment of ssDNA. The double-headed green arrow indicates the orientation of the electric dipole transition moment (EDTM). (***B***) An iCy3 dimer labeled DNA fork construct contains the dimer probe near the ss – ds DNA fork junction. The conformation of the Cy3 dimer probe reflects the local secondary structure of the sugar-phosphate backbones at the probe insertion site position. The sugar-phosphate backbones of the conjugate DNA strands are shown in black and blue, the bases in gray, and the iCy3 chromophores in green. (***C***) The structural parameters that define the local conformation of the iCy3 dimer probe are the inter-chromophore separation vector *R*_*AB*_, the tilt angle *θ*_*AB*_, and the twist angle *ϕ*_*AB*_. The electrostatic coupling between the iCy3 chromophores gives rise to the anti-symmetric (−) and symmetric (+) excitons, which are indicated by the red and blue arrows, respectively, and whose magnitudes and transition energies depend on the structural parameters. (***D***) The insertion site position of the iCy3 dimer probe is indicated relative to the pseudo-fork junction using positive integers in the direction towards the double-stranded region, and negative integers in the direction towards the single-stranded region.

The linear absorbance spectrum of the free Cy3 chromophore in solution, as well as when it is attached covalently to a nucleic acid, exhibits a pronounced vibronic progression, which can be simulated using a Holstein-Frenkel Hamiltonian with electronic transition energy *ε*_*eg*_ = ∼18,250 cm^-1^, vibrational frequency *ω*_0_ = ∼1100 cm^-1^ and Huang-Rhys electronic-vibrational coupling parameter *λ*^2^ = ∼0.55 (5). The electric dipole transition moment (EDTM) has magnitude *μ*_*eg*_ = ∼12.8 D and orientation that lies parallel to the Cy3 trimethine bridge (see Fig. 1A). Cy3 can be chemically attached ‘internally’ to the DNA with the ‘iCy3’ acting as a molecular bridge between bases as an extension of the sugar-phosphate backbone (6). When two complementary single strands of DNA with opposed iCy3 labeling positions are annealed, an (iCy3)_2_ dimer probe is formed within the DNA duplex. If the sequence of nt bases at or near one side of the (iCy3)_2_ dimer is non-complementary, the labeling location occurs at a ss-dsDNA fork junction, as shown schematically in Fig. 1*B*. iCy3 monomer-labeled ss-dsDNA constructs are similarly prepared with a thymine (T) base at the position opposite to the probe within the complementary strand.

Because of the relatively small separation between iCy3 chromophores within the (iCy3)_2_ dimer-labeled ss-dsDNA fork constructs, the monomer EDTMs (labeled as sites *A* and *B* in Fig. 1*C*) can couple through a resonant electrostatic interaction *J*. This coupling gives rise to symmetric (+) and anti-symmetric (−) excitons with orthogonally polarized dipole moments 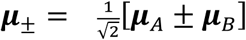 and with relative magnitudes that depend on the local conformation of the (iCy3)_2_ dimer probe. The symmetric and anti-symmetric excitons each consist of a manifold of delocalized electronic-vibrationally coupled states, which are superpositions of electronic-vibrational product states of the *A* and *B* monomer sites (7, 8). The absorbance and circular dichroism (CD) spectra of (iCy3)_2_ dimer labeled ss-dsDNA constructs are well-described using the Holstein-Frenkel model, and can be used to determine local conformational parameters (5, 7). The structural parameters that characterize the dimer conformation are the ‘tilt’ angle *θ*_*AB*_, the ‘twist’ angle *ϕ*_*AB*_, and the separation *R*_*AB*_. In our previous spectroscopic studies of (iCy3)_2_ dimer labeled ss-dsDNA constructs, we calculated the resonant electrostatic coupling by treating the EDTMs as point dipoles (7). We later repeated these calculations using an ‘extended-dipole’ model (9, 10) that more accurately accounted for the extension of the transition charge density across the length of the molecule, and which yielded nearly identical results to those provided by the point-dipole approximation (5). We note that more accurate models of electrostatic coupling for Cy3, which are based on *ab initio* calculations of atomic transition charges, have recently become available and provide future opportunities to test the validity of point-dipole and extended dipole models (11).

Here we focus on the distributions of structural parameters obtained from theoretical analyses of absorbance, CD, and two-dimensional fluorescence spectroscopy (2DFS) of (iCy3)_2_ dimer labeled ss-ds DNA fork constructs. While absorbance and CD can be used to determine the mean structural parameters of the (iCy3)_2_ dimer probes, 2DFS provides additional information about the distributions of these parameters. The underlying optical principles of 2DFS resemble those of 2D NMR (12, 13) and 2DIR, with the latter providing structural and dynamic information about local vibrational modes in proteins (14, 15), nucleic acids (16, 17) and biomolecular hydration shells (18). 2DIR has been used to distinguish sequence-dependent inter-base H-bonds in duplex DNA (16, 17) and the rearrangements of water molecules at or near the exposed surfaces of DNA strands (18). While these relatively fast processes likely contribute to nucleic acid stability and dynamics, they do not directly probe the DNA breathing fluctuations involved in protein recognition events (19, 20). In contrast, the signals detected by 2DFS on (iCy3)_2_ dimer probe labeled ss-dsDNA constructs do directly monitor DNA backbone conformations and conformational disorder, which likely play a central role in protein recognition and binding events.

In the following experiments, we studied several different ss-dsDNA fork constructs in which we varied the (iCy3)_2_ dimer labeling position, as shown in Fig. 1*D*, and for some of these constructs we varied the temperature. In contrast to the (iCy3)_2_ dimer ss-dsDNA constructs, the linear spectra of the iCy3 monomer ss-dsDNA constructs are relatively insensitive to probe labeling position and temperature, as previously reported (5, 7). These findings suggest that for the (iCy3)_2_ dimer ss-dsDNA constructs, the sensitivity of the homogeneous lineshapes to labeling position and temperature are due largely to variations of the coupling interaction *J*, which is sometimes referred to as ‘off-diagonal disorder’ in the reference frame of the monomer sites (15, 21).

In our prior studies, we established that a combination of absorbance and CD spectra contain sufficient information to determine mean values of the structural parameters 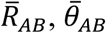 and 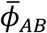, in addition to an estimate of the inhomogeneous line broadening parameter *σ*_*I*_ (5, 7). Inhomogeneous line broadening is a direct measure of structural heterogeneity due to individual molecules of the sample exhibiting uniquely different homogeneous lineshapes, which depends on the local conformation of the (iCy3)_2_ dimer probe. Our prior estimates of *σ*_*I*_ were obtained from a deconvolution of absorbance and CD spectra and were based on the value of the homogeneous line width Γ_*H*_ = ∼186 cm^-1^. We determined the latter value from room temperature 2DFS experiments on iCy3 monomer and (iCy3)_2_ dimer labeled DNA constructs in which the probe labeling position was deep within the double-stranded region, and for which the laser bandwidth was Δ*λ*_*L*_ = ∼16 nm (7). In the current work, we perform a more accurate analysis of 2DFS data in which the laser bandwidth was Δ*λ*_*L*_ = ∼33 nm. This increase in laser bandwidth permits us to determine simultaneously the homogeneous and inhomogeneous line shape parameters as a function of probe labeling position and temperature. In addition, we here extend our line shape analysis to model the distributions of the structural parameters *R*_*AB*_, *θ*_*AB*_, and *ϕ*_*AB*_, because variation in these parameters ‘builds in’ the structural heterogeneity measured by our 2DFS experiments. In the analyses that follow, we have assumed that the distributions of structural parameters can be successfully modeled as Gaussians, which can be characterized using the standard deviations *σ*_*R*_, *σ*_*θ*_ and *σ*_*ϕ*_.

Among the significant findings of this work is that the (iCy3)_2_ dimer is a reliable probe of the local conformation of the sugar-phosphate backbones at and near ss-dsDNA fork junctions, which depends sensitively on the labeling site position and temperature. We studied the temperature-dependence of the local conformation of (iCy3)_2_ dimer labeled ss-dsDNA constructs, both at positions deep within the duplex region (+15) and near the ss-dsDNA fork junction (−1). We find that local conformations and conformational disorder of the sugar-phosphate backbones at the +15 position are minimized at room temperature (23°C) and change rapidly as the temperature is either raised or lowered away from room temperature under physiological salt conditions (100 mM NaCl, 6 mM MgCl_2_), permitting the development of local conformations that deviate significantly from the W-C duplex DNA structure, such as bubbles, bends and kinks. In contrast, local conformations, and conformational disorder of the ss-dsDNA junction at the −1 position do not vary significantly with increasing temperature, suggesting that the distribution of thermally accessible states is relatively narrow. Moreover, the mean local conformation and conformational disorder vary systematically with (iCy3)_2_ dimer labeling position (from +2 to −2, refer to Fig. 1*C* for probe labeling nomenclature). This transition is characterized by an increase in conformational disorder and a loss of cylindrical symmetry characterized by the mean tilt angle 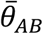, followed by a change in the local symmetry of the DNA backbones from right-handed to left-handed, as reflected by the mean twist angle 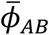. Perhaps contrary to expectations, regions of the ss-dsDNA junction extending into the single strands appear to be relatively well-ordered.

## 2. Experimental Methods

### A. Sample preparation

The sequences and nomenclature of the iCy3 monomer and (iCy3)_2_ dimer labeled ss-ds DNA constructs used in our studies are shown in Table 1. Oligonucleotide samples were purchased from Integrated DNA Technologies (IDT, Coralville, IA) and used as received. Solutions were prepared containing ∼1 μM oligonucleotide in 10 mM TRIS buffer with 100 mM NaCl and 6 mM MgCl_2_. Complementary strands were combined in equimolar concentrations. The samples were heated to 95°C for 4 minutes and left to cool slowly on a heat block overnight prior to data collection. The annealed iCy3 monomer and (iCy3)_2_ dimer labeled ss-ds DNA fork constructs contained both ds and ss DNA regions, with the probe labeling positions indicated by the nomenclature described in Fig. 1*D*. The iCy3 monomer labeled constructs contained a thymine base (T) in the complementary strand at the position directly opposite to the probe chromophore.

**Table 1.**
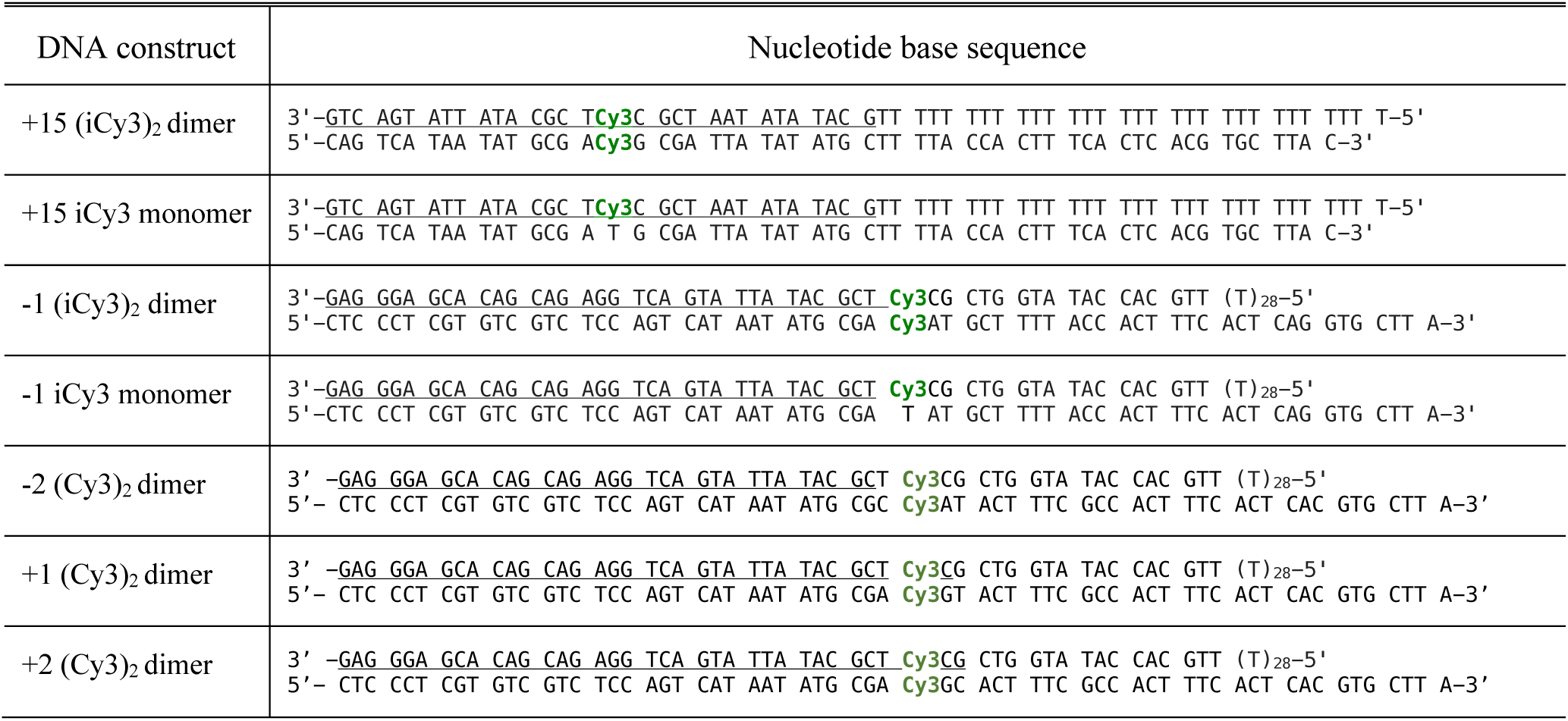
Base sequences and nomenclature for the iCy3 monomer and (iCy3)_2_ dimer ss-ds DNA fork constructs used in these studies. The horizontal lines indicate regions of complementary base pairing.

### B. Absorbance and circular dichroism (CD) measurements

We performed linear absorbance measurements using a Cary 3E UV-Vis spectrophotometer, and CD measurements with a JASCo model J-720 CD spectrophotometer. Series of temperature-dependent measurements were performed over the range 1 – 75°C. For all absorbance and CD measurements, the samples were housed in a 1 cm quartz cuvette. CD measurements over the temperature range 1 – 25°C were performed using a JASCo J-1500 CD spectrophotometer equipped with a Koolance EXOS liquid cooling system, which can operate reliably at near-freezing temperatures.

### C. Two-dimensional fluorescence spectroscopy (2DFS)

Phase-modulation 2DFS experiments were performed on the iCy3 labeled ss-ds DNA fork constructs listed in Table I using methods and procedures described previously (5, 7, 22-25). The train of four collinear laser pulses used to excite the sample was centered on wavelength *λ*_*L*_ = ∼532 nm (∼18,800 cm^-1^), with bandwidth Δ*λ*_*L*_ = ∼33 nm (∼1,100 cm^-1^). The pulses were generated using a custom-built non-collinear optical parametric amplifier (NOPA) that was pumped using a 140 kHz Ti:Sa regenerative amplifier (Coherent, RegA). The NOPA output was divided into two paths using a 50/50 beam-splitter, and each beam was directed to a separate Mach-Zehnder interferometer (MZI). Acousto-optic Bragg cells, placed within the beam paths of each MZI, were used to apply a relative temporal phase sweep to the pulses exiting the MZI. Thus, the relative phase of pulses 1 and 2, and that of pulses 3 and 4, were swept continuously at the frequencies Ω_21_ = 5 kHz and Ω_43_ = 8 kHz, respectively. The relative paths of the pulses were varied using computer-controlled translation stages to step the time delay *t*_21_ between the first pair of pulses, and the delay *t*_21_ between the second pair of pulses. For all our measurements, the time delay *t*_43_ between the second and third pulse was set to zero. For each combination of time delays, the four pulses were used to excite resonant electronic transitions of the iCy3 probes, and the ensuing fluorescence was detected and demodulated simultaneously at the sum frequency Ω_43_ + Ω_21_ = 13 kHz and the difference frequency Ω_43_ − Ω_21_ = 3 kHz, which correspond, respectively, to the fourth-order non-rephasing (NRP) and rephasing (RP) signals (23, 25, 26).

The optical pulses were compressed using a quadruple-pass fused-silica prism pair to compensate for dispersive media in the optical paths preceding the sample. Pulse widths were characterized by placing a beta-barium borate (BBO) frequency doubling crystal at the sample position where a phase-modulated train of pulse pairs was incident. The frequency-doubled signal output was detected using a lock-in amplifier, which was referenced to the waveform signal used to modulate the relative phase of the pulses (22, 25). The pulse compressor was adjusted so that the full-width-at-half-maximum (FWHM) of the pulse-pulse autocorrelation, for each of the pulse pairs, was Δ*τ*_*L*_ = ∼28 fs. We measured the laser bandwidth Δ*λ*_*L*_ = ∼33 nm (∼1,100 cm^-1^) centered at *λ*_*L*_ = ∼532 nm using an Ocean Optics spectrometer. The measured time-bandwidth product was thus 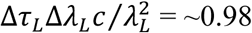.

The laser pulse spectrum with the above spectral properties was reproducibly maintained and continuously monitored during each 2DFS measurement described in this work. Samples were housed in a 1 mm quartz cuvette that was mounted to a small aluminum heating block, which was itself placed in thermal contact with a copper block equipped with internally circulating cooling water. The temperature of the sample was maintained to within ∼±0.1 °C using two thermoelectric chips, which were mounted directly to the aluminum block. Fluorescence was detected at a 45° angle of incidence relative to the front face of the sample cuvette using a 5 mm collection lens and a 615 nm long-pass filter (Chroma, HQ615LP), which served to minimize scattered excitation light. A light stream of nitrogen was continuously flowed across the front face of the cuvette to prevent condensation of vapor for measurements performed at reduced temperatures.

## 3. Theoretical Modeling

### A. Simulation of absorbance and CD spectra

We simulated the absorbance and CD spectra of the iCy3 monomer and (iCy3)_2_ dimer ss-ds DNA fork constructs (see Table I) by applying the Holstein–Frenkel (H–F) model (8, 27), as described in our previous studies (5, 7). The H-F model treats each iCy3 monomer as a two-electronic-level molecule with ground state |*g*⟩ and excited state |*e*⟩, which are coupled to a single harmonic vibrational mode of frequency *ω*_0_. The electronic-vibrational coupling is characterized by the Huang-Rhys parameter, 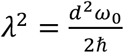, where *d* is the displacement of the minimum of the electronically excited vibrational potential energy surface relative to that of the ground state surface. The Huang-Rhys parameter corresponds physically to the number of vibrational quanta absorbed by the system upon electronic excitation. Each monomer (*M*, labeled *A* and *B*) is chemically identical with electronic transition energy, *ε*_*eg*_, and electric dipole transition moment (EDTM), 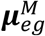. The Hamiltonian operator representing the monomer is given by

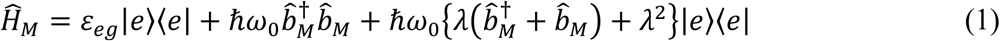

where 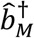 and 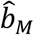 are, respectively, the operators for creating and annihilating a vibrational excitation in the electronic potential energy surfaces. These operators obey the commutation relation 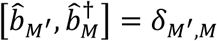, where *δ*_*M*′,*M*_, is the Kronecker delta function and the monomer labels *M* ′, *M* ∈ {*A, B*}.

The monomer absorbance spectrum is a weighted sum of homogeneous lineshapes associated with the individual vibronic transitions from the initially unexcited ground state, |*g* ⟩ |*n*_*g*_ = 0 ⟩:

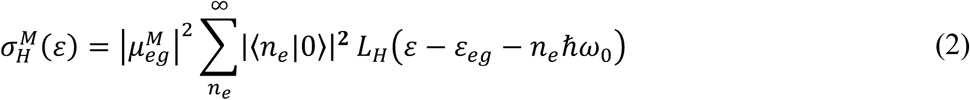

In Eq. (2) the index *n*_*e(g)*_ is the vibrational quantum number of the electronically excited (unexcited) potential energy surface, the homogeneous line shape is given by the Lorentzian, 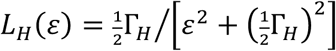, with FWHM line width equal to Γ_*H*_, and the individual peak intensities depend on the Franck-Condon factors,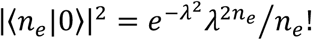.

To account for the influence of varying local environments on the transition energy (i.e., spectral inhomogeneity), we describe the total absorbance spectrum as a Voigt convolution integral (28)

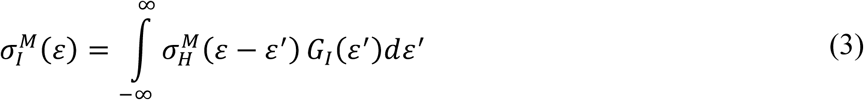

In Eq. (3) the Gaussian distribution, 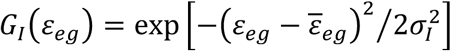, represents the probability that a given monomer has its transition energy relative to an average value 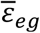, and *σ*_*I*_ is the standard deviation of the inhomogeneously broadened spectrum.

The Hamiltonian of the *AB* dimer is written

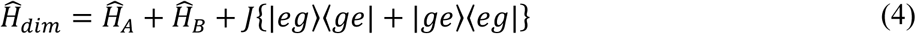

where the final term couples the singly excited electronic transitions of the monomers through an electrostatic interaction *J*. In our nomenclature used here |*eg*⟩ represents the product state in which monomer *A* is electronically excited and monomer *B* is unexcited. The electronic coupling parameter *J* depends on the dimer conformation and can be modeled in terms of the Coulomb interaction between the individual monomer transition charge densities 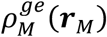

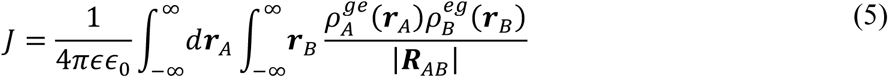

where ***R***_*AB*_= ***r***_*B*_ − ***r***_*A*_. In the current work we approximate the electrostatic coupling parameter using the ‘extended’ transition dipole-dipole model, which accounts for the finite size of the iCy3 chromophore by including a one-dimensional displacement vector ***l*** that lies parallel to the monomer EDTM (9, 10).

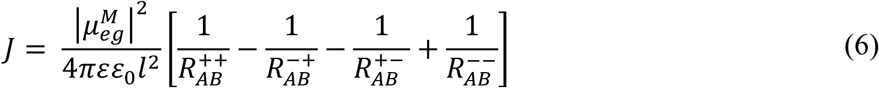

Equation (6) assumes that the transition charge density for each monomer is composed of two point charges of equal magnitude (*q*) and opposite sign separated by the distance *l*, such that 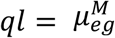 (5, 10). The distances between point charges are given by 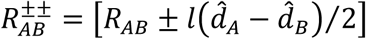 and 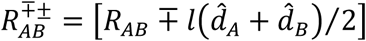, where 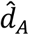 and 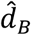 are unit vectors that lie parallel to the monomer EDTMs. For the calculations that follow, we used the values 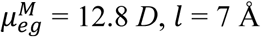 and *q* = 0.38*e* (where *e* is the electronic charge unit), as in our previous studies (5). In the extended dipole model, the value of *J* depends on the inter-chromophore separation *R*_*AB*_, the twist angle *ϕ*_*AB*_, and the tilt angle *θ*_*AB*_ (see Fig. 1*C*), which collectively specify the (iCy3)_2_ dimer conformation.

For a given value of the coupling parameter *J*, we determined the eigen-energies and the eigen-states of the dimer Hamiltonian given by Eq. (4). The singly excited states are symmetric (+) and anti-symmetric (−) superpositions of electronic-vibrational product states 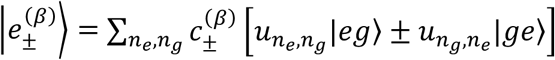, where the coefficients 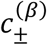 depend on the vibrational coordinates of the monomers, 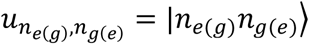 is the vibrational product state, and *β* = (0, 1, …) is a state index that advances in order of increasing energy (8).

The dimer absorbance spectrum is the sum of symmetric (+) and anti-symmetric (−) exciton features

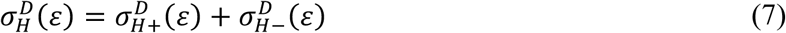

where 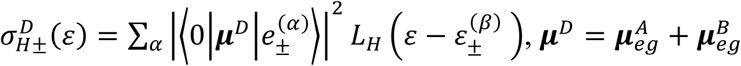 is the collective EDTM and 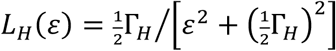 is a Lorentzian function that represents the homogeneous lineshape of the transition with eigen-energy 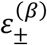 and FWHM line width Γ_*H*_. Similarly, the dimer CD spectrum is the sum of symmetric and anti-symmetric rotational strengths

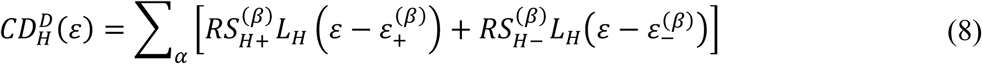

where 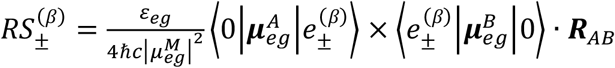. In the above expressions, we have defined the ground vibrational-electronic state of the *AB* dimer |0⟩ = |*v*_*A*_ = 0, *v*_*B*_ = 0⟩|*gg*⟩.

The (iCy3)_2_ dimer conformation may vary from molecule to molecule due to local DNA ‘breathing’ fluctuations, so that the homogeneous dimer absorbance and CD line shapes are convolved with an inhomogeneous distribution function 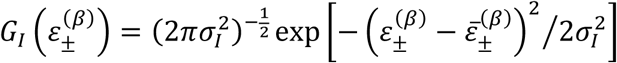, which is centered at the average transition energy 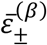 and has standard deviation *σ*_*I*_. The final expressions for the absorbance and CD spectra are given, respectively, by the Voigt profiles:

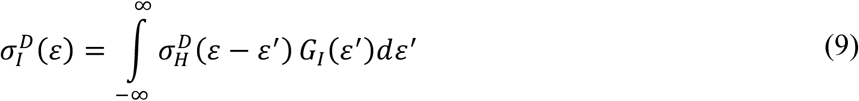

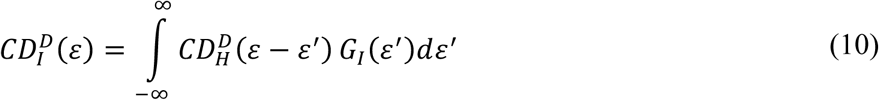

### B. Simulation of two-dimensional fluorescence spectra (2DFS)

We applied the H-F model to simulate our 2D fluorescence spectra according to methods developed previously (23, 26). The 2DFS signals are written in terms of the rephasing (RP) and non-rephasing (NRP) fourth-order response functions

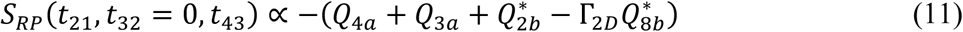

and

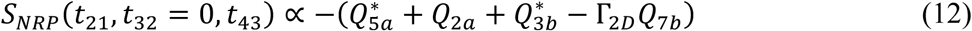

In Eqs. (11) and (12), the first two terms on the right-hand sides of the proportionalities represent, respectively, the ground state bleach (GSB) and stimulated emission (SE) contributions. The final two terms represent excited state absorption (ESA) contributions for the singly-and doubly-excited-state manifolds, respectively. The parameter Γ_2*D*_ represents the fluorescence quantum yield of the doubly-excited-state-manifold relative to that of the singly excited state manifold. The possible values of Γ_2*D*_ may range from 0 to 2, which has the effect of modifying the sign and magnitude of the ESA contributions relative to those of the GSB and SE. In our analyses of the 2D spectral lineshapes of the iCy3 monomer and (iCy3)_2_ dimer ss-dsDNA constructs discussed below, we treated Γ_2*D*_ as one of three parameters (the others being the homogeneous and inhomogeneous line width parameters, Γ_*H*_ and *σ*_*I*_, respectively) that were optimized to our experimental data. For all our calculations, we found that the optimized value for Γ_2*D*_ was ∼ 0.3 (see Fig. S1 of the SI).

In writing the response functions, we designate |*v*⟩ = |*v*_*A*_*v*_*B*_⟩|*gg*⟩ as the dimer state with both monomers electronically unexcited and with vibrational quantum number *v* = *v*_*A*_ *+ v*_*B*_, such that, for example, state |0⟩ is the electronic ground state with zero vibrational occupancy. The states |*e*⟩ and |*e*′⟩ represent any two of the symmetric and anti-symmetric excitons 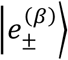 within the singly-excited-state manifold, and the state |*f*⟩ represents any one state within the doubly-excited-state manifold. When the effects of inhomogeneous broadening are included, the individual terms of the RP response functions can be written (29, 30):

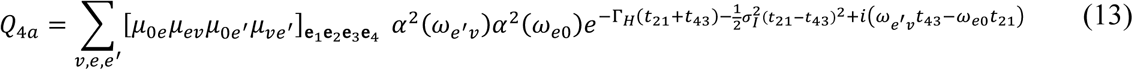

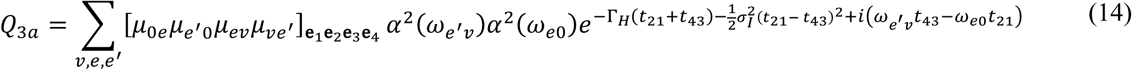

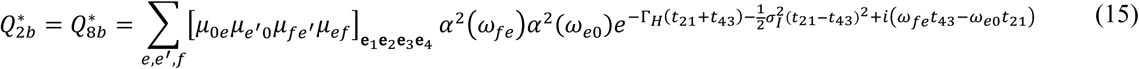

and for NRP:

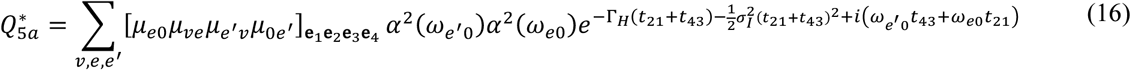

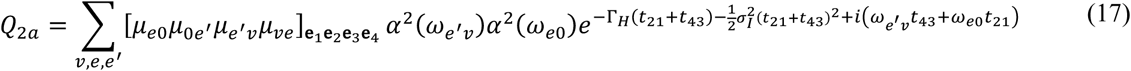

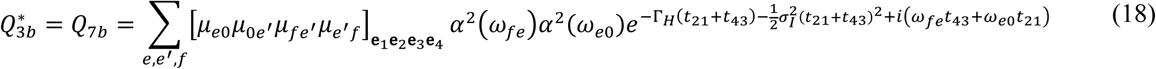

In Eqs (13) – (18), the factor 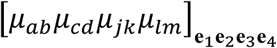 denotes the orientationally averaged four-point product, ⟨(***μ***_*ab*_ ⋅ **e**_1_)(***μ***_*cd*_ ⋅ **e**_2_)(***μ***_*jk*_ ⋅ **e**_3_)(***μ***_*lm*_ ⋅ **e**_4_)⟩, which accounts for the projections of the (iCy3)_2_ dimer transition dipole moments onto the (parallel) plane polarizations of the four laser pulses and includes an average over the isotropic distribution of dimer orientations in solution (31). The factor *α*^2^(*ω*_*ab*_)*α*^2^(*ω*_*cd*_) is the product of the intensities of the laser at the transition frequencies *ω*_*ab*_ and *ω*_*cd*_.

An important feature of the RP and NRP response functions is their distinct dependences on the homogeneous and inhomogeneous line width parameters, Γ_*H*_ and *σ*_*I*_, respectively (30, 32) (see Fig. 2) The RP response functions [Eqs. (13) – (15)] contain the lineshape factor 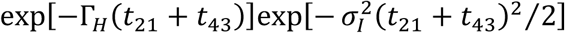, which decays exponentially at the homogeneous dephasing rate along the diagonal axis (*t*_21_ + *t*_43_), and as a Gaussian envelope with inhomogeneous decay constant 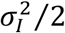 along the anti-diagonal axis (*t*_21_ *– t*_43_) (see Fig. 2*A*, top panel). In contrast, the NRP response functions [Eqs. (16) – (18)] contain the lineshape factor 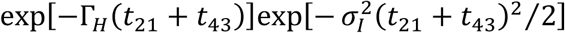, which decays along the diagonal axis with rate constants that depend on both the homogeneous and inhomogeneous parameters (see Fig. 2*A*, bottom panel). Examples of the RP and NRP response functions are shown in Fig. 2*B*. The RP and NRP 2D fluorescence spectra, which are functions of the optical frequencies 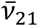 and 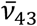, are obtained by performing two-dimensional Fourier transforms of the response functions with respect to the delay variables *t*_21_ and *t*_43_ given by Eqs. (11) and (12), respectively, (Fig. 2*C*).

**Figure 2.**
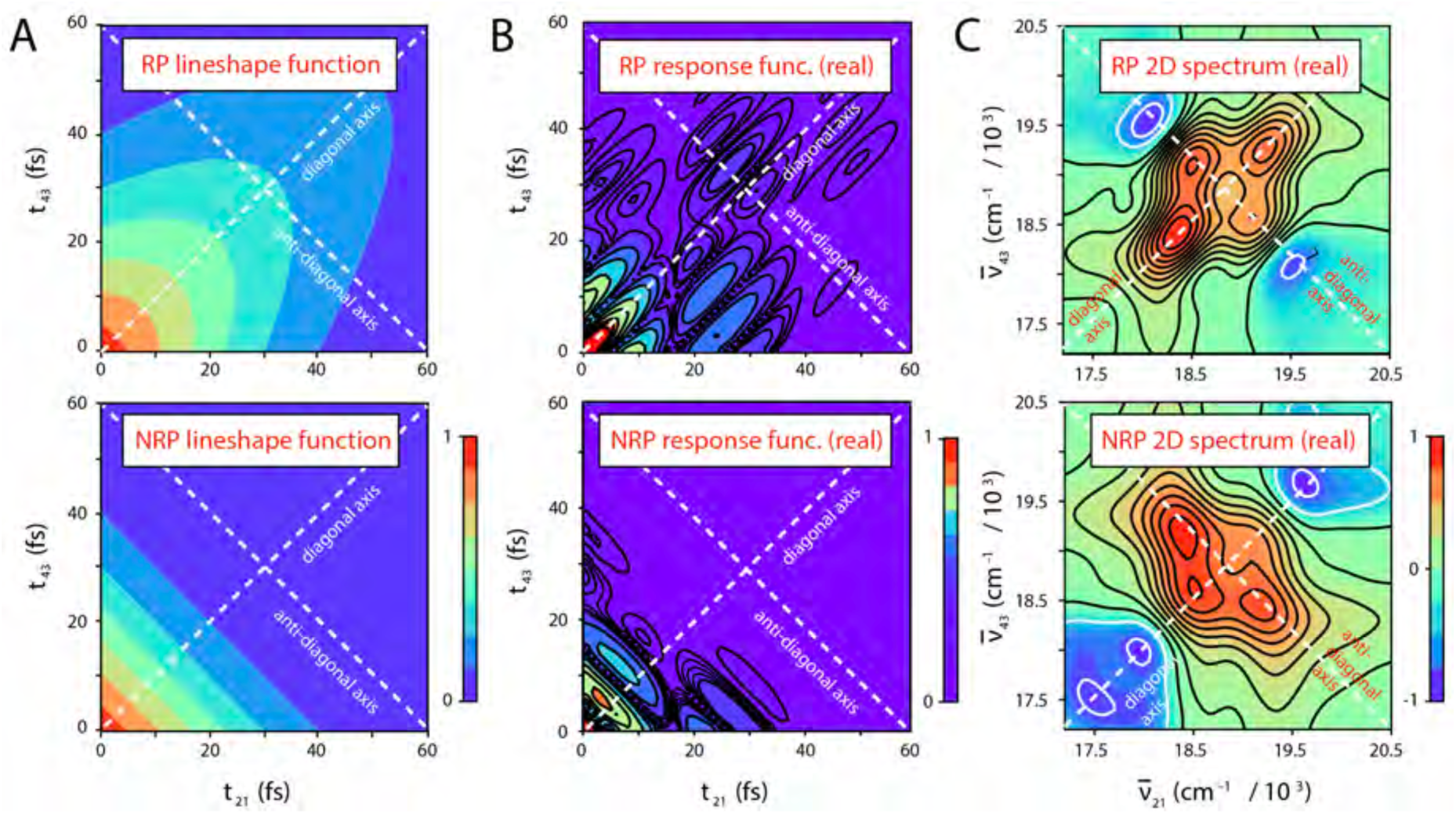
Example calculations of the 2DFS rephasing (RP) and non-rephasing (NRP) response functions and 2D spectra for the −1 iCy3 monomer ss-dsDNA construct. All functions are displayed as two-dimensional contour plots with diagonal and anti-diagonal axes indicated as white dashed lines. (***A***) The RP lineshape function (top panel), 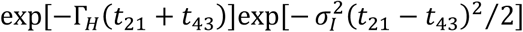, contains two independent factors, one representing an exponential (homogeneous) decay along the diagonal axis (*t*_21_ + *t*_43_), and the other a Gaussian (inhomogeneous) decay along the anti-diagonal axis (*t*_21_ − *t*_43_). The NRP lineshape function (bottom panel), 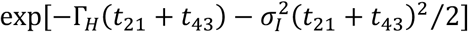, depends on both the homogeneous and inhomogeneous line width parameters, which contribute to the decays along both diagonal and anti-diagonal axes. (***B***) The absolute value of the real parts of the RP response function *S*_*RP*_(*t*_21_, *t*_32_ = 0, *t*_43_) [Eq. (11), top panel] and the NRP response function *S*_*NRP*_(*t*_21_, *t*_32_ = 0, *t*_43_) [Eq. (12), bottom panel] contain the lineshape functions shown in panel *A* and the transition frequency phase factors given by Eqs. (13) – (18). (***C***) The RP 2D spectrum 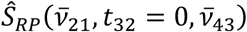 (top panel) and the NRP spectrum 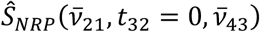 are calculated by performing a two-dimensional Fourier transform of the response functions shown in panel *B* with respect to the time variables *t*_21_ and *t*_43_.

In the limiting case for which spectral inhomogeneity greatly exceeds the homogeneous line width (*σ*_*I*_ ≫ Γ_*H*_), individual features of the RP 2D spectrum exhibit a Lorentzian lineshape in the direction of the anti-diagonal axis 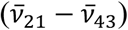 and an inhomogeneously broadened lineshape in the direction of the diagonal axis 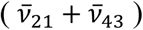 (30). In this limit, the homogeneous and inhomogeneous line width parameters can be determined directly from the RP 2D spectrum by fitting the anti-diagonal and diagonal cross-sections of the 2D spectral lineshape to model Lorentzian and Gaussian functions, respectively. However, in the more general case of moderate inhomogeneity (*σ*_*I*_ ≃ Γ_*H*_), the homogeneous and inhomogeneous broadening mechanisms each contribute to the RP 2D lineshape in both the diagonal and anti-diagonal directions. In our analyses of the 2D spectral lineshapes of the iCy3 monomer and (iCy3)_2_ dimer ss-dsDNA constructs presented below, we determined the homogeneous and inhomogeneous lineshape parameters by simultaneously fitting experimental RP and NRP 2DFS spectra to the numerical Fourier transforms of the model response functions given by Eq. (11) and (12). This approach provided an accurate description of the 2D spectra without imposing any assumed restrictions on the degree of inhomogeneity present.

### C. Numerical optimization procedures

In previous work (5, 7), we characterized the iCy3 monomer absorbance spectrum using four independent parameters: the mean electronic transition energy 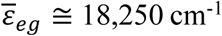, the Huang-Rhys vibronic coupling parameter *λ*^2^ ≅ 0.57, the vibrational frequency *ω*_0_ ≅ 1,100 cm^-1^, and the spectral inhomogeneity parameter *σ*_*I*_ ≅ 300 cm^-1^. We determined these values by performing a numerical optimization procedure in which we directly compared the simulated spectra [Eq. (3)] to experimental data. In our previous studies, we assumed constant values for the homogeneous FWHM line width Γ_*H*_ = 186 cm^-1^ and the monomer EDTM 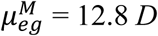, which we determined in separate experiments (7). We used these monomer parameters as inputs to our analyses of the absorbance and CD spectra of the (iCy3)_2_ dimer ss-dsDNA constructs [Eqs. (9) and (10), respectively]. In our calculations of the dimer spectra, we included six vibrational levels in the monomer electronic-vibrational manifold of states to ensure numerical convergence (7). The dimer absorbance and CD spectra were thus used to obtain optimized values of the mean structural parameters 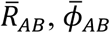, and 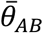, which determined the mean electrostatic coupling 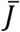 [Eq. (6)].

We next used the set of optimized structural coordinates as inputs to our model analyses of the RP and NRP 2DFS data, which are described by Eqs. (11) and (12), respectively. The transition frequencies, *ω*_*ab*_ and *ω*_*cd*_, appearing in the RP and NRP response functions [Eqs. (13) – (18)], in addition to the laser spectral overlap factors *α*^2^(*ω*_*ab*_)*α*^2^(*ω*_*cd*_) and the transition-dipole orientation factors 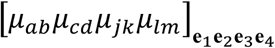, are all constants determined by the values of the mean structural coordinates. For our simulations of the 2DFS data, it was necessary to carry out the sums of Eqs. (13) – (18) over the space of transition pathways between the dimer ground electronic-vibrational state manifold (labeled |*v*⟩, with dimension 6 × 6 = 36), the singly-excited electronic-vibrational state manifold (labeled |*e*⟩ and |*e* ′⟩, with dimension 36 + 36 = 72) and the doubly-excited electronic-vibrational state manifold (|*f*⟩, with dimension 36). Nevertheless, the actual number of terms needed to simulate the response functions accurately is a small fraction of the total number of possible pathways, due to the resonance conditions imposed by the laser pulse spectrum [reflected by the factors *α*^2^(*ω*_*ab*_)*α*^2^(*ω*_*cd*_)]. In practice, for each response function the summation over transition pathways was calculated and stored as a two-dimensional interferogram that was multiplied by the lineshape function 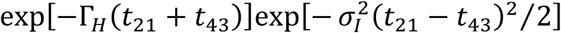 in the case of the RP response functions [Eqs. (13) – (15)], and the lineshape function 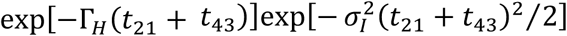 in the case of the NRP response functions [Eqs. (16) – (18)]. We thus performed optimization analyses of our 2DFS data to obtain accurate values for the homogeneous and inhomogeneous line width parameters, Γ_*H*_ and *σ*_*I*_, respectively.

For our optimization calculations, we implemented an automated multi-variable regression analysis to efficiently explore the parameter space of the spectroscopic models. We have applied similar procedures in past studies (5, 7, 23, 26, 33-35), in which a random search algorithm was used to select an initial set of input parameters that were refined iteratively using commercial software (KNITRO) (36). For each set of input trial parameters, we calculated a linear least squares error function *χ*^2^, which was minimized to obtain the optimized solution. Thus, for our optimizations of the absorbance and CD spectra of the (iCy3)_2_ dimer ss-dsDNA constructs, we minimized the function

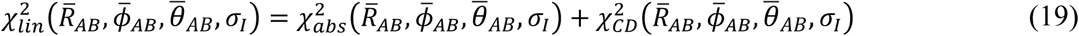

and for our optimizations of 2DFS data, we minimized the function

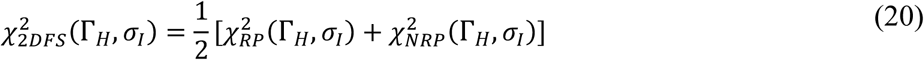

We performed error-bar analyses of the optimized parameters, which we determined by a 1% deviation of the *χ*^2^ function from its optimized value.

### D. Modeling conformational heterogeneity of (iCy3)_2_ dimer labeled ss-dsDNA constructs

As discussed in previous sections, the information provided by the linear absorbance and CD spectra of the (iCy3)_2_ dimer permits us to determine the mean values of the conformational coordinates 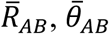, and 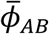. By expanding the analysis to include 2DFS data, we determined additional information about the inhomogeneously broadened distribution of homogeneous lineshapes, which is due primarily to the variation of local (iCy3)_2_ dimer conformations within the ensemble of ss-dsDNA molecules. To develop our interpretation of the inhomogeneous lineshape in terms of structural disorder, we assumed that the conformational coordinates, *R*_*AB*_, *θ*_*AB*_ and *ϕ*_*AB*_, can be treated as independent variables, and that their distributions can be described as a product of Gaussians

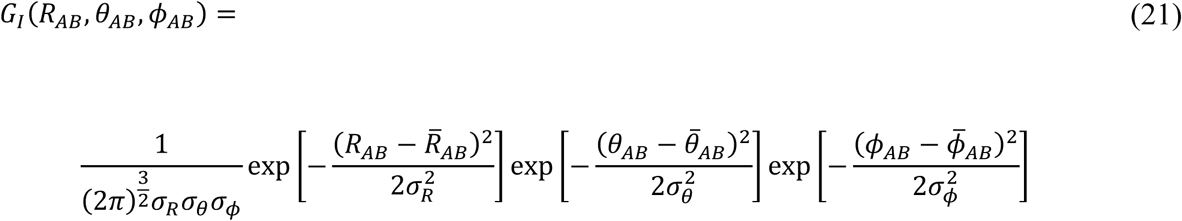

We emphasize that by assuming that the structural coordinates are independent variables – i.e., that possible covariance terms are negligible – Eq. (21) may only be used to determine estimates of the standard deviations.

To model our inhomogeneously broadened 2DFS data, we calculated a library of ‘homogeneous’ RP and NRP 2D fluorescence spectra spanning a range of equally spaced values for the conformational coordinates, and for which the homogeneous and inhomogeneous line width parameters were set equal – i.e., Γ_*H*_ = *σ*_*I*_ = 100 cm^-1^. In Fig. 3*A*, we show examples of simulated homogeneous RP and NRP 2D fluorescence spectra of the +2 ss-dsDNA construct corresponding to three different values of the mean twist angle 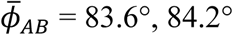 and 84.8°, and for mean twist angle 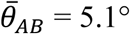 and mean separation 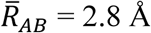. From the library of homogeneous 2D spectral lineshapes, we simulated inhomogeneously broadened RP and NRP 2D spectra by numerically sampling the library according to the Gaussian distribution given by Eq. (21). We thus followed a procedure similar to that described in previous sections to iteratively calculate the linear least squares error function 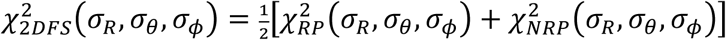 between simulated and experimental spectra, which we minimized to obtain optimized values for the standard deviations of the conformational parameters, *σ*_*R*_, *σ*_*θ*_ and *σ*_*ϕ*_. Optimized values were obtained according to the definitions discussed below, which depended on the functional dependence of the error function on the standard deviation parameter.

**Figure 3.**
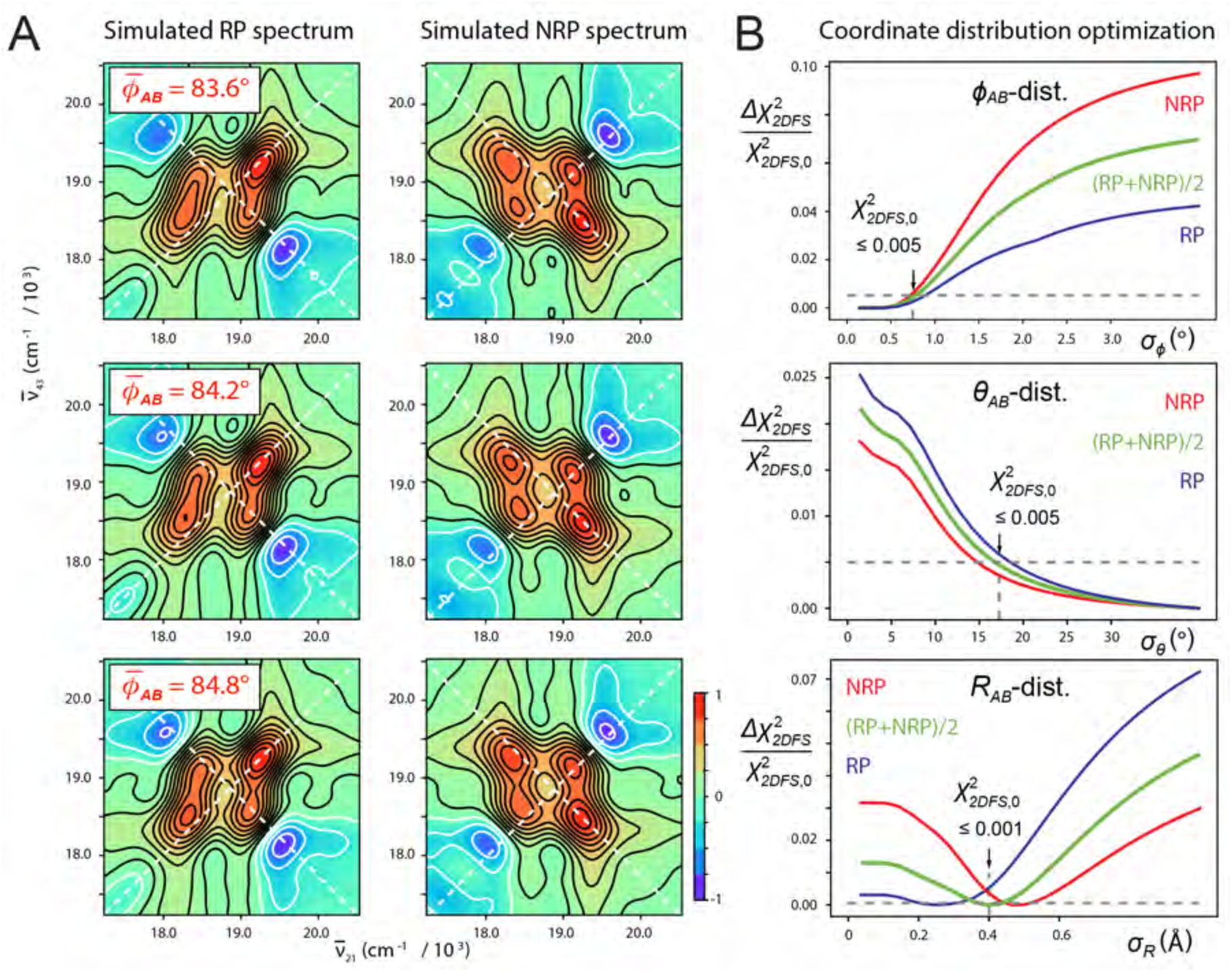
(***A***) Simulated RP and NRP ‘homogeneous’ 2D fluorescence spectra (real part) of the +2 (iCy3)_2_ dimer ss-dsDNA construct for various values of the mean twist angle: 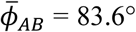 (top row), 84.2° (middle) and 84.8° (bottom), mean tilt angle 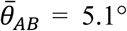, mean separation 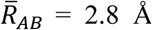, and homogeneous and inhomogeneous linewidth parameters Γ_*H*_ = *σ*_*I*_ = 100 cm^-1^. (***B***) Cross-sections of the relative deviation of the linear least squares error function, 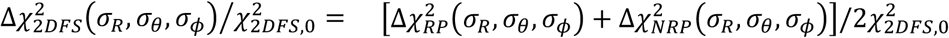, are shown as functions of the standard deviations *σ*_*ϕ*_ (top), *σ*_*θ*_ (middle) and *σ*_*R*_ (bottom). Inhomogeneously broadened spectra of the +2 (iCy3)_2_ dimer ss-dsDNA construct were simulated by numerically sampling the library of ‘homogeneous’ 2D fluorescence spectra according to the Gaussian distribution of structural coordinates given by Eq. (21). Error function cross-sections are shown plotted relative to their ‘optimized’ values, 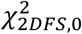 (indicated by vertical arrows), which are defined as the 0.5% threshold for cases in which the function approached its minimum asymptotically (as do the *σ*_*ϕ*_ and *σ*_*θ*_ cross-sections), and the 0.1% threshold for cases in which the function exhibited a distinct minimum (as shown for the *σ*_*R*_ cross-section).

In Fig. 3*B*, we show example cross-sections of the relative deviation of the linear least squares error function 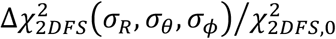 and their RP and NRP contributions, 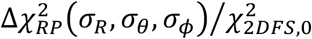 and 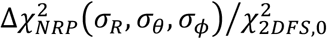, respectively, with 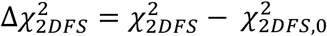. These functions are plotted relative to their ‘optimized’ values, 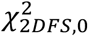, which we defined as the 0.5% threshold (i.e., 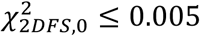) in cases for which the minimum is approached asymptotically. In cases for which the function exhibited a distinct minimum, we defined the optimized value such that 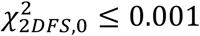. For the error function plotted along the *σ*_*ϕ*_-axis (Fig. 3*B*, top), we see that 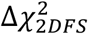 approaches an asymptotic minimum for small values of the standard deviation and increases abruptly for *σ*_*ϕ*_ ≥ 0.7°. This indicates that the distribution of the local twist angle parameter for the +2 (iCy3)_2_ ss-dsDNA construct is relatively narrow, and that the optimized value for *σ*_*ϕ*_ is an upper bound. In contrast, the cross-section plotted along the *σ*_*θ*_-axis (Fig. 3*B*, middle) decreases monotonically with increasing standard deviation, indicating that the optimized value *σ*_*θ*_ = 17° is a lower bound. The average cross-section plotted along the *R*_*AB*_-distribution axis (Fig. 3*B*, bottom) exhibits a distinct minimum with optimized value *σ*_*R*_ = 0.4 Å. In all the panels shown in Fig. 3*B*, threshold values are indicated by horizontal dashed lines.

### E. Determination of the relative fluorescence quantum yield parameter Γ_2*D*_

The value used for the parameter Γ_2*D*_, which characterizes the relative fluorescence quantum yield of the doubly versus singly excited state populations [described by Eqs. (11) and (12)], is important for fitting 2DFS data to theoretical models (23, 26, 37, 38). To determine the value of Γ_2*D*_, we performed an optimization analysis of our temperature-dependent 2DFS data for the (iCy3)_2_ dimer labeled +15 (duplex) ss-dsDNA construct based on the H-F Hamiltonian model. In these calculations, we minimized the function 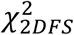 described by Eq. (20) while varying the parameter Γ_2*D*_. The procedure was carried out for data sets taken at five different temperatures (5, 15, 23, 35 and 45 °C). The remaining input parameters for the H-F model were obtained from the optimized fits to the linear absorbance and CD spectroscopic measurements taken at the same temperatures, as discussed further below. We show the results of these calculations in Fig. S1 of the SI. For each temperature, we observe a progression of the parameter Γ_2*D*_ that favors lower values except for 25 °C, for which the analysis is relatively insensitive to the value of Γ_2*D*_. We therefore adopted the value Γ_2*D*_ = 0.3 for all the (iCy3)_2_ dimer calculations presented in the remainder of this work.

## 4. Results and Discussion

### A. Local conformations and spectral inhomogeneity of the iCy3 monomer and (iCy3)_2_ dimer ss-dsDNA constructs labeled at the +15 ‘duplex’ and −1 ‘fork’ positions

In previous studies, we examined the temperature-dependent absorbance and CD spectra of iCy3 monomer and (iCy3)_2_ dimer labeled ss-dsDNA constructs, in which the chromophore probes were positioned either at the +15 position (deep within the duplex region) or at the −1 position relative to the ss-dsDNA fork junction (see Fig. 1*C* and Table 1) (5, 7). We found that the structural parameters and coupling strengths that characterized the absorbance and CD spectra of the (iCy3)_2_ dimer ss-dsDNA constructs varied with probe labeling position and temperature. In Fig. 4, we compare our results for the room temperature (25 °C) CD, absorbance and 2DFS measurements of the iCy3 monomer and (iCy3)_2_ dimer labeled +15 ‘duplex’ and −1 ‘fork’ ss-dsDNA constructs.

**Figure 4.**
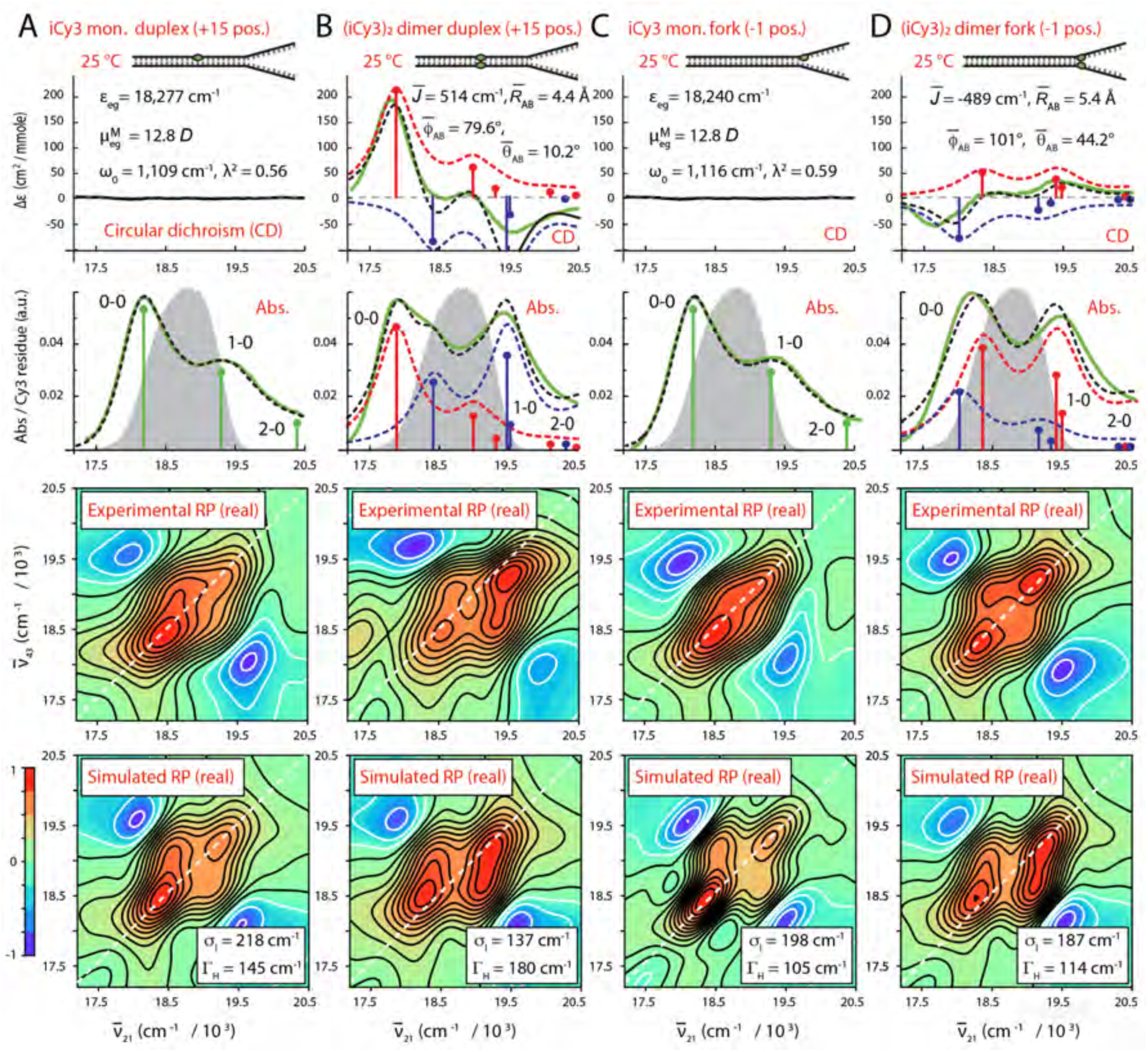
Experimental and simulated spectroscopic measurements of iCy3 monomer and (iCy3)_2_ dimer labeled +15 ‘duplex’ and −1 ‘fork’ ss-dsDNA constructs performed at room temperature (25 °C). (***A***) iCy3 monomer +15 ss-dsDNA construct; (***B***) (iCy3)_2_ dimer +15 ss-dsDNA construct; (***C***) iCy3 monomer −1 ss-dsDNA construct; and (***D***) (iCy3)_2_ dimer −1 ss-dsDNA construct. The experimental CD (top row) and absorbance spectra (second row) are shown (in green) overlaid with vibronic spectral features (black dashed curves) obtained from the optimized fits to the H-F model. For the monomer constructs (***A*** and ***C***), the vibronic features are shown in green, and for the dimer constructs (***B*** and ***D***) the symmetric (+) and anti-symmetric (–) excitons are shown in blue and red, respectively. Values of optimized parameters are shown in the insets of the corresponding panels. The laser spectrum, with center frequency 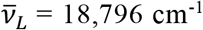 (*λ*_*L*_ = 532 nm) and FWHM bandwidth 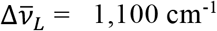 (Δ*λ*_*L*_= ∼33 nm), is shown (in gray) overlaid with the absorbance spectra and spans a region containing both the 0-0 and 1-0 vibronic sub-bands, as shown. Experimental RP spectra (third row) are compared to the optimized simulated RP spectra (fourth row). Simulated spectra are based on the structural parameters obtained from our optimization analyses of the CD and absorbance spectra, the fluorescence quantum yield parameter Γ_2*D*_= 0.3, and the homogeneous and inhomogeneous line width parameters (Γ_*H*_ and *σ*_*I*_, respectively), which are listed for the (iCy3)_2_ dimer labeled constructs in Table 2 and Table 3, and for the iCy3 monomer labeled constructs in Table S1 and Table S2 of the SI.

We first consider the CD, absorbance and 2D fluorescence spectra of the iCy3 monomer labeled +15 ‘duplex’ and −1 ‘fork’ ss-dsDNA constructs (see Fig. 4*A* and *C*, respectively), which are well-described using the monomer Hamiltonian [Eqs. (1) – (3)], as expected. Values obtained from model fits to the absorbance and CD spectra for the mean electronic transition energy *ε*_*eg*_, electric dipole transition moment 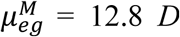, vibrational frequency *ω*_0_, and Huang-Rhys parameter *λ*^2^ are shown in the insets. The laser spectrum used for the 2DFS experiments is shown (in gray) overlaid with the absorbance spectra (second row) and spans the spectral region containing the 0-0 and 0-1 vibronic transitions of the monomer (shown as green line segments). The signatures of these transitions are present in the experimental 2DFS data (RP spectra, third row). We simulated the 2DFS data (fourth row) using the monomer Hamiltonian and the optimized parameters obtained from the CD and absorbance spectra, as described in Sect. 3B. We note the excellent agreement between simulated and experimental 2D spectra. Our analyses of the 2DFS data thus provide optimized values for the homogeneous and inhomogeneous line width parameters of the iCy3 monomer labeled ss-dsDNA constructs: Γ_*H*_ = 145 cm^-1^ and *σ*_*I*_ = 218 cm^-1^ for the +15 ss-dsDNA construct, and Γ_*H*_ = 105 cm^-1^ and *σ*_*I*_ = 198 cm^-1^ for the −1 ss-dsDNA construct. Optimized values for the iCy3 monomer Hamiltonian parameters for the +15 and −1 ss-dsDNA constructs, and their associated error bars, are listed in Table S1 and Table S2 of the SI, respectively.

**Table 2.** Mean structural parameters and 2DFS line widths determined for the (Cy3)_2_ dimer labeled +15 ‘duplex’ ss-dsDNA construct. The parameters determined from model analyses of linear absorbance and CD spectra are the mean resonant coupling 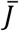, the mean twist angle 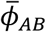, the mean tilt angle 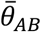, and the mean interchromophore separation 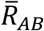. The parameters determined from the 2DFS data are the inhomogeneous and homogeneous line widths *σ*_*I*_ and Γ_*H*_. Error bars were calculated based on a 1% deviation of the *χ*^2^ function from its minimum (optimized) value.

| Optimized Parameters for (iCy3)<sub>2</sub> Dimer +15 ‘Duplex’ ss-dsDNA Constructs |  |  |  |  |  |  |
| --- | --- | --- | --- | --- | --- | --- |
| From Absorbance and CD Optimization |  |  |  |  | From 2DFS Optimization |  |
| $T$ (°C) | $\bar{J}$ (cm<sup>-1</sup>) | $\bar{\phi}_{AB}$ (°) | $\bar{\theta}_{AB}$ (°) | $\bar{R}_{AB}$ (Å) | $\sigma_I$ (cm<sup>-1</sup>) | $\Gamma_H$ (cm<sup>-1</sup>) |
| 1 | 493 | 78.3+0.3/-0.1 | 44.2+1.4/-0.4 | 3.49+0.9/-0.6 | 121.5 +35.4/-32.4 | 105.1+22.2/-17.4 |
| 5 | 488 | 79.3+0.2/-0.1 | 39.3+0.7/-0.4 | 2.7+1.1/-0.4 | 212.7 +36.8/-31.0 | 109.5 +21.2/-17.9 |
| 15 | 532 | 80.7 +0.2/-0.1 | 18.1 +2.9/-2.1 | 3.7 ± 0.1 | 141.8 +34.3/-33.4 | 149.4 +19.7/-18.7 |
| 23 | 514 | 79.6 ± 0.2 | 10.2 +6.4/-27 | 4.4 ± 0.1 | 136.7 +45.8/-40.1 | 180.4 +25.0/-20.3 |
| 35 | 496 | 76.0 ± 0.3 | 3.0 +13/-19 | 5.5 ± 0.1 | 253.2 +27.6/-25.3 | 91.8 +14.0/-14.4 |
| 45 | 483 | 71.9 ± 0.4 | 7.7 +8.5/-24 | 6.4 ± 0.1 | 268.4 +31.8/-26.4 | 78.5 +17.0/-12.0 |
| 55 | 467 | 70.4 ± 0.5 | 7.0 +8.4/-22 | 6.8 ± 0.1 | 253.2 +36.2/-33.9 | 60.8 +18.7/-13.6 |
| 65 | 354 | 73.7 ± 0.6 | 5.4 +11/-22 | 7.1 ± 0.1 | 217.8 +31.4/-25.2 | 56.3 +16.7/-14.1 |

**Table 3.**
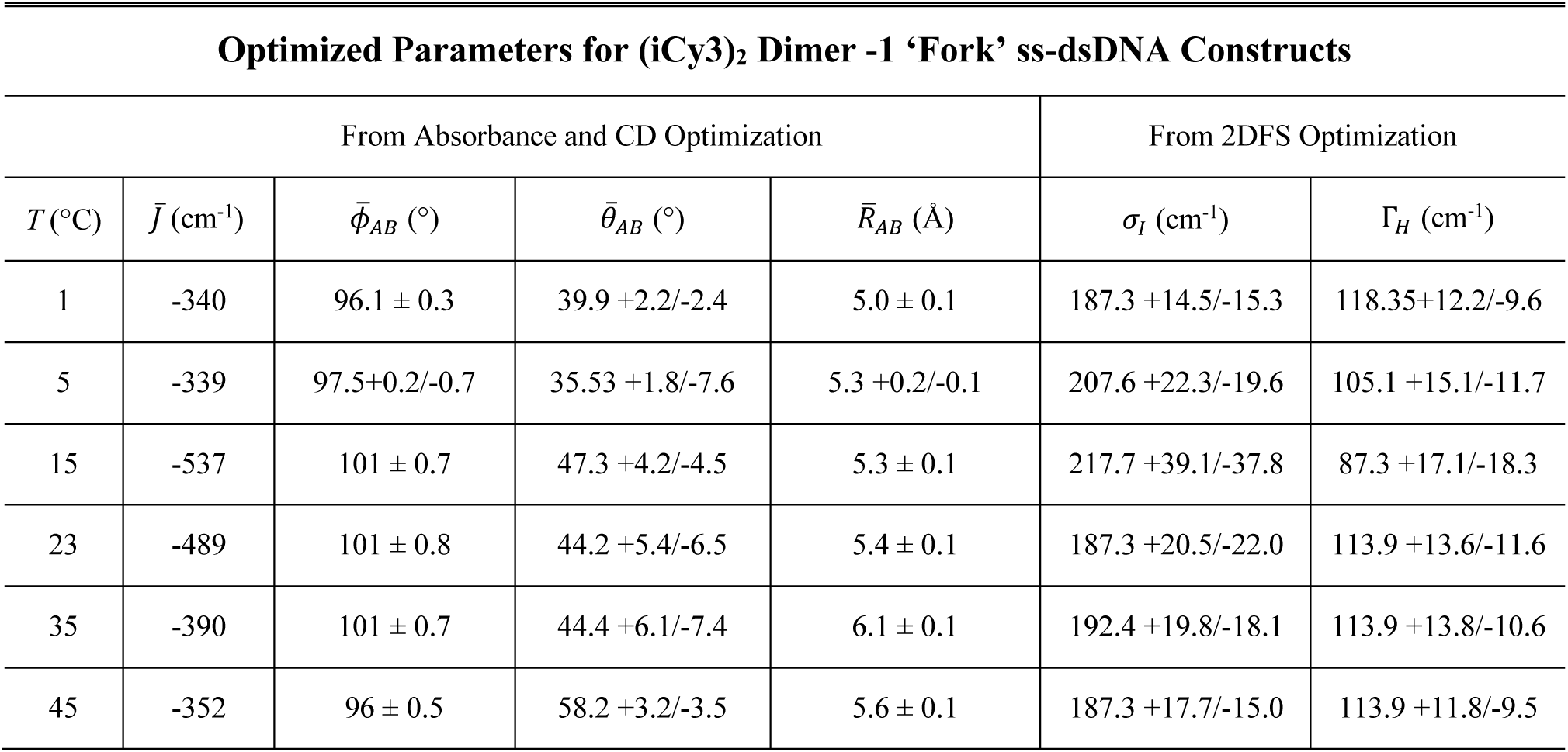
Mean structural parameters and 2DFS line widths determined for the (Cy3)_2_ dimer labeled −1 ‘fork’ ss-dsDNA construct. The parameters determined from model analyses of spectroscopic data are the same as defined in Table 2.

For the (iCy3)_2_ dimer labeled +15 ‘duplex’ and −1 ‘fork’ ss-dsDNA constructs, the CD, absorbance and 2DFS data are well described using the H-F dimer Hamiltonian [Eqs. (4) – (10)] (see Fig. 4*B* and 4*D*, respectively). Mean values of the structural parameters and electronic coupling are shown in the insets. The simulated symmetric (+) and anti-symmetric (–) vibronic manifolds are shown overlaid with the experimental spectra as blue and red dashed curves, respectively. The symmetries of the CD spectra, and the corresponding signs of the electrostatic couplings 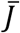, determine the handedness of the (iCy3)_2_ dimer conformation. The optimization of the H-F model to the +15 ‘duplex’ ss-dsDNA construct provides values for the mean separation, twist and tilt angles 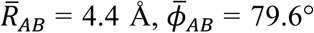 and 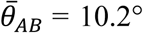, respectively, which indicates that the sugar-phosphate backbones adopt a right-handed, cylindrically symmetric local conformation at the +15 position consistent with the Watson-Crick B-form crystallographic structure of duplex DNA. In contrast, the optimized values obtained for the −1 ‘fork’ ss-dsDNA construct are 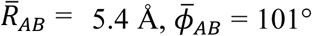 and 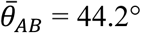, which indicates that the sugar-phosphate backbones at the −1 position adopt, on average, a left-handed, splayed open conformation. The experimental 2DFS data (third row) exhibit diagonal and off-diagonal peaks, which are due to optically resonant transitions involving the symmetric and anti-symmetric vibronic manifolds. The 2DFS data were simulated (fourth row) using the same optimized structural parameters obtained from the CD and absorbance spectra according to the procedure discussed in Sect. 3B. We note that the simulated and experimental 2DFS data are in excellent agreement. As we discuss further below, from our analyses of the 2DFS lineshapes we obtained optimized values of the homogeneous and inhomogeneous line width parameters for the (iCy3)_2_ dimer labeled ss-dsDNA constructs: Γ_*H*_ = 180 cm^-1^ and *σ*_*I*_ = 137 cm^-1^ for the +15 ss-dsDNA construct, and Γ_*H*_ = 114 cm^-1^ and *σ*_*I*_ = 187 cm^-1^ for the −1 ss-dsDNA construct.

The spectral inhomogeneity that we determined from our room temperature 2DFS lineshape analyses was generally larger for the iCy3 monomer labeled constructs than for the (iCy3)_2_ dimer labeled constructs. This finding is especially evident for the +15 ss-dsDNA constructs and suggests that the local environment of the sugar-phosphate backbones of the iCy3 monomer probes is more disordered than that of the (iCy3)_2_ dimer probes. We note that the (iCy3)_2_ dimer ss-dsDNA constructs have the two monomer probes positioned directly opposite to one another in a symmetric manner, while the monomer labeled ss-dsDNA constructs have a single thymine (T) base positioned on the conjugate strand directly opposite to the iCy3 probe. Monomer substitution may thus introduce a ‘defect site,’ which is more disruptive to the local conformation and dynamics of the sugar-phosphate backbones than dimer substitution at the same probe labeling positions at room temperature. This finding is consistent with recent studies of the sensitivity of cyanine monomer substituted DNA constructs to the local environment (39), which can influence fluorescence intensity, local mobility and photostability (6).

### B. Temperature-dependent local conformations and spectral line width parameters of iCy3 monomer and (iCy3)_2_ dimer ss-dsDNA constructs labeled at the +15 ‘duplex’ and −1 ‘fork’ positions

We carried out temperature-dependent absorbance, CD and 2DFS measurements of both the iCy3 monomer and the (iCy3)_2_ dimer labeled +15 and −1 ss-dsDNA constructs. In previous work, we reported the results of our temperature-dependent studies of the absorbance and CD spectra of these constructs to determine optimized structural and spectroscopic parameters using the monomer and H-F Hamiltonian models (5, 7). Although these studies determined the mean values of the monomer and dimer Hamiltonian parameters, including the mean structural parameters 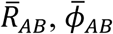 and 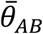, they could not provide accurate assessments for the homogeneous and inhomogeneous line widths as we report in the current study. Comparisons between our temperature-dependent experimental 2DFS data and optimized simulations for the monomer and dimer labeled +15 ‘duplex’ ss-dsDNA constructs are shown in Figs. S2 – S5 and Figs. S6 – S9 of the SI, respectively. Similar comparisons for the monomer and dimer labeled −1 ‘fork’ ss-dsDNA constructs are shown in Figs. S10 – S13 and Figs. S14 – S17 of the SI, respectively. In Table 2 and Table 3, we list for these same constructs the mean structural parameters obtained from the absorbance and CD spectra, and the associated line width parameters obtained from our analyses of the 2DFS data over the full range of temperatures studied. The optimized monomer Hamiltonian parameters for the +15 and −1 ss-dsDNA constructs are listed in Table S1 and Table S2 of the SI.

In Fig. 5, we illustrate the temperature-dependence of the absorbance (panel *A*), CD (panel *B*), experimental 2DFS data (panel *C*) and simulated 2DFS data (panel *D*) for the (iCy3)_2_ dimer labeled +15 ss-dsDNA construct. The data presented in Fig. 5 and the corresponding parameters listed in Table 2 show that for this construct the mean coupling strength is maximized at 15°C 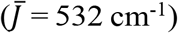, and decreases systematically with increasing and decreasing temperature. At the lowest temperatures (1 – 25°C), the absorbance and CD spectra exhibit, respectively, the intensity borrowing and bi-signate lineshapes that are characteristic of the vibronically coupled iCy3 dimer system (7, 8). The H-F analysis of the absorbance and CD data indicate that the sugar-phosphate backbones of the duplex adopt a progressively increasing mean tilt angle with decreasing temperature, while the mean twist angle does not change significantly (Table 2). Moreover, the 2DFS data at low temperatures show well-separated and relatively narrow peaks and cross-peaks, which indicate the presence of the delocalized symmetric (+) and anti-symmetric (−) excitons.

**Figure 5.**
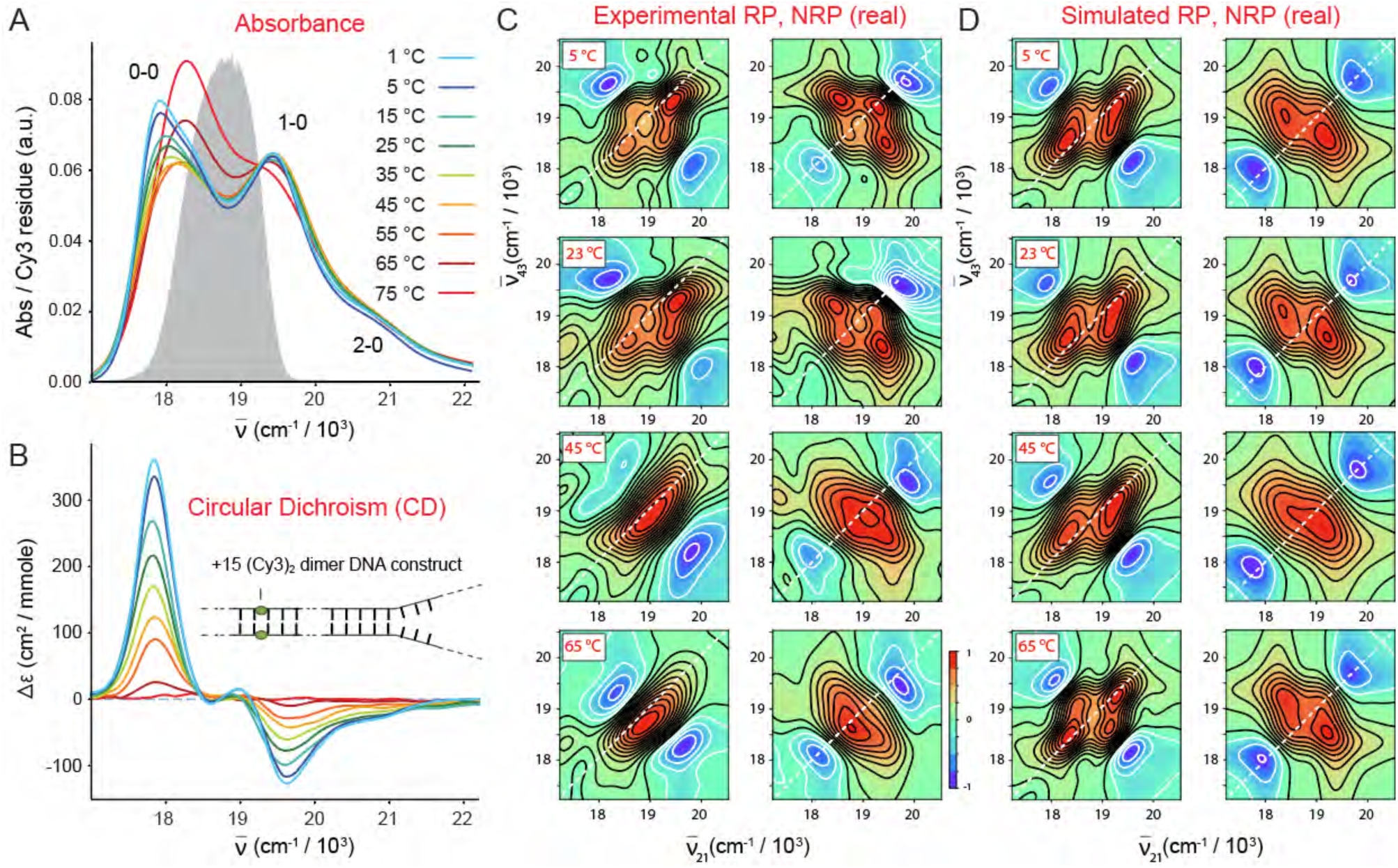
Temperature-dependent spectroscopic measurements of the (iCy3)_2_ dimer +15 ss-dsDNA construct. (***A***) absorbance, (***B***) CD, (***C***) experimental 2DFS and (***D***) simulated 2DFS. The gray curve in panel (***A***) is the same laser spectrum as shown in Fig. 4. In panels (***C***) and (***D***), both RP (left columns) and NRP (right columns) 2D spectra are shown. Additional comparison between the experimental 2DFS data and the optimized simulated spectra for the (iCy3)_2_ dimer +15 ss-dsDNA construct are presented in Figs. S6 – S9 of the SI.

The simulated 2DFS data shown in Fig. 5*D*, which assumes the optimized structural parameters obtained from the H-F analysis of the absorbance and CD, are in excellent agreement with the experimental data and provide an accurate determination of the homogeneous and inhomogeneous lineshape parameters. As the temperature is increased, the absorbance and CD spectral lineshapes change systematically to reflect the decrease in the mean coupling strength. At the highest temperature for which the single strands of the duplex have fully separated (75°C) the absorbance and CD spectra resemble those of the iCy3 monomer substituted ss-dsDNA constructs. The spectral features of the 2DFS data, which are well-defined at low temperatures, become progressively less pronounced as the temperature is increased. We note an abrupt change in the experimental 2DFS lineshapes at temperatures above 23°C in which the intensities of the off-diagonal features decrease and the diagonal features merge into a single diffuse feature. The simulations of the 2DFS data capture this behavior with increasing temperature, which is due to increasing the optical dephasing rate while simultaneously decreasing the electronic coupling strength.

As mentioned previously, we simulated our 2DFS data for the iCy3 monomer and (iCy3)_2_ dimer +15 ‘duplex’ and −1 ‘fork’ labeled ss-dsDNA constructs using the same temperature-dependent Hamiltonian parameters that we determined from analyses of absorbance and CD data (see Table 2 and Table 3) (5, 7). In general, our simulations of the 2DFS spectra exhibited peaks and cross-peaks with positions and relative intensities that matched closely our experimental results. We thus performed optimization calculations on our 2DFS data to determine the best fit homogeneous and inhomogeneous line width parameters, Γ_*H*_ and *σ*_*I*_, respectively, as a function of temperature. These results are presented in Table 2 and Table 3, and the line width parameters are plotted in Fig. 6. We note that in the case of the (iCy3)_2_ dimer −1 ss-dsDNA ‘fork’ construct, the highest temperature that we investigated is 45°C, which is ∼20 degrees lower than the melting point of the double-stranded region of the ss-dsDNA constructs, *T*_*m*_ ∼65°C. This is due to the relatively low CD signals of this ss-dsDNA construct at elevated temperatures, which prevented us from obtaining accurate values of the structural parameters above 45°C. As we discuss further below, the structural parameters of this construct appear to have converged to plateau values at temperatures below 45°C.

**Figure 6.**
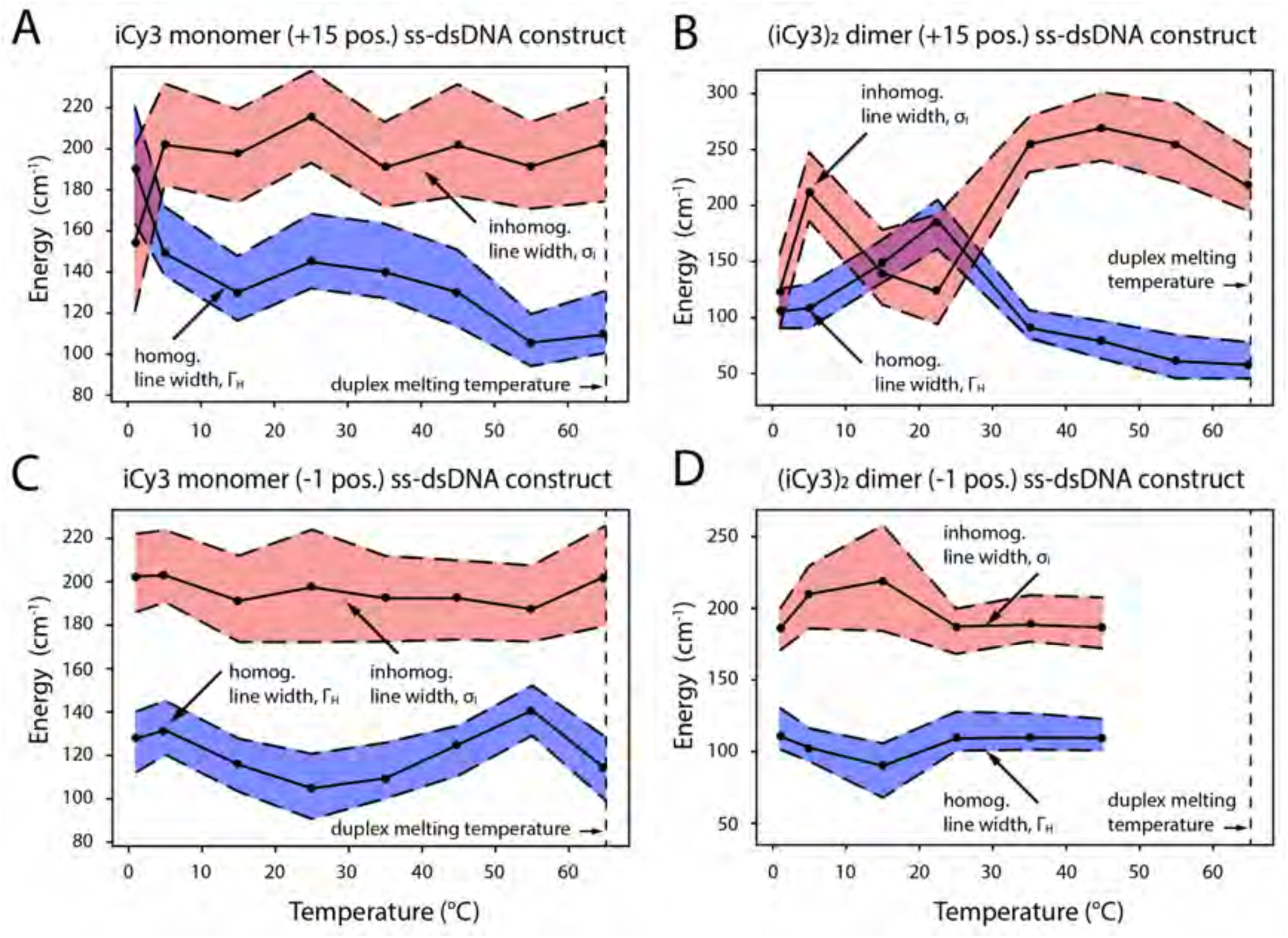
Optimized homogeneous and inhomogeneous line width parameters as a function of temperature obtained from 2DFS lineshape analyses. (***A***) iCy3 monomer +15 ss-dsDNA construct; (***B***) (iCy3)_2_ dimer +15 ss-dsDNA construct; (***C***) iCy3 monomer −1 ss-dsDNA construct; and (***D***) (iCy3)_2_ dimer −1 ss-dsDNA construct. Shaded regions bounded by dashed lines indicate error bars, which were calculated based on a 1% deviation of the 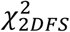 function [Eq. (20)] from its minimum value. The vertical dashed line at 65°C indicates the melting temperature *T*_*m*_ of the duplex regions of the DNA constructs. Direct comparisons between experimental and optimized simulated 2DFS data are presented in Figs. S2 – S17 of the SI.

In our previous studies we estimated the inhomogeneous line widths based solely on analyses of linear spectroscopic data, for which we assumed that the homogeneous line width was constant (Γ_*H*_ = 186 cm^-1^) for all temperatures and probe labeling positions (7). The results of those studies suggested that for both the iCy3 monomer and (iCy3)_2_ dimer labeled +15 (duplex) ss-dsDNA constructs, the inhomogeneous line width parameter increased monotonically with temperature over the range 15 – 65°C. However, for the iCy3 monomer and (iCy3)_2_ dimer labeled −1 (fork) ss-dsDNA constructs, the inhomogeneous line width parameter remained relatively constant over this same temperature range.

In Fig. 6, we present the results of our 2DFS lineshape analysis, which provides a far more detailed picture of the temperature- and position-dependent behavior of the homogeneous and inhomogeneous line width parameters (shown as blue- and teal-shaded regions, respectively). The optimized values of the line width parameters are presented as points and the shaded regions bounded by dashed lines indicate error bars, which are based on a 1% deviation of the 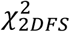 function [see Eq. (20)] from its optimized value. We note that for both the iCy3 monomer and (iCy3)_2_ dimer +15 ‘duplex’ ss-dsDNA constructs (Figs. 6*A* and 6*B*), our results at 1°C appear to behave as ‘outliers’ from the remaining temperature-dependent data shown over the range ∼5 – 65°C, which we discuss below.

We first consider the line width parameters corresponding to the iCy3 monomer labeled +15 ‘duplex’ (Fig. 6*A*) and the −1 ‘fork’ (Fig. 6*C*) ss-dsDNA constructs. Our results indicate that the two iCy3 monomer labeled constructs exhibit qualitatively similar temperature dependencies of the inhomogeneous line widths. At the lowest temperatures (∼5 – 15°C) the inhomogeneous line width (*σ*_*I*_ ∼200 cm^-1^) is significantly larger in magnitude than the homogeneous line width (Γ_*H*_ ∼100 – 140 cm^-1^), which suggests there is significant conformational disorder of the sugar-phosphate backbones labeled by the iCy3 monomer probe in both the duplex region and at the fork −1 position. Furthermore, the inhomogeneous line widths undergo very little variation over the temperature range 5 – 65°C, indicating a relatively constant level of conformational disorder of the sugar-phosphate backbones as the temperature is increased towards the melting point. We note that for the iCy3 monomer +15 ss-dsDNA construct, there is an increase of the inhomogeneous lineshape parameter from *σ*_*I*_ ∼150 – 200 cm^-1^ between 1 – 5°C, suggesting an abrupt increase of local disorder near the monomer probe site within the duplex region at the lowest temperature.

The homogeneous line widths of both iCy3 monomer labeled ss-dsDNA constructs exhibit more complicated temperature-dependent behavior. In the case of the iCy3 monomer labeled +15 ‘duplex’ ss-dsDNA construct, the value of the homogeneous line width decreases rapidly from Γ_*H*_ ∼190 cm^-1^ to ∼130 cm^-1^ over the temperature range 1 – 15°C, followed by a slight increase from Γ_*H*_ ∼130 cm^-1^ to ∼145 cm^-1^ over the range 15 – 25°C, and a gradual decrease from Γ_*H*_ ∼145 cm^-1^ to ∼105 cm^-1^ over the range 25 – 55 °C. The highest temperature of 65 °C corresponds to the melting point of the duplex region, for which 50% of the single strands are expected to be completely separated. At the melting point we observe a significant reduction of the homogeneous line width (Γ_*H*_ ∼114 cm^-1^) in comparison to the value we obtained for the same constructs at room temperature (Γ_*H*_ ∼145 cm^-1^). In contrast, for the iCy3 monomer labeled −1 ‘fork’ ss-dsDNA construct, the homogeneous line width decreases gradually from Γ_*H*_ ∼132 cm^-1^ to ∼105 cm^-1^ over the range 5 – 25 °C, followed by an increase from Γ_*H*_ ∼105 cm^-1^ to ∼140 cm^-1^ over the range 25 – 55°C. It is interesting to note that while the homogeneous line width of the iCy3 monomer +15 ss-dsDNA construct is maximized at room temperature (Γ_*H*_ ∼145 cm^-1^), the value of the homogeneous line width of the −1 ‘fork’ ss-dsDNA construct is minimized at room temperature (Γ_*H*_ ∼105 cm^-1^).

The homogeneous line width is related to the total dephasing time (*T*_2_) according to *T*_2_ = (*πc*Γ_*H*_)^−1^(≈ 100 fs for Γ_*H*_ ≈ 100 cm^-1^). The total dephasing time can be written in terms of the population relaxation time (*T*_1_) and the pure dephasing time 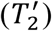, according to 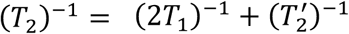 (32). The value of *T*_1_ can be estimated from the room temperature fluorescence lifetime *τ*_*F*_ ∼162 ps (4). Although the fluorescence lifetime of iCy3 labeled DNA constructs can vary with temperature due to, for example, thermally activated photoisomerization (4, 40-42), such processes are many orders of magnitude slower than the tens-of-femtosecond time scales of pure dephasing. The homogeneous line width is thus dominated by pure dephasing, which depends on interactions between the electronic transitions and the phonon bath. This suggests that we may interpret the temperature-dependent variation of the homogeneous line width in terms of changes to the iCy3 probe’s direct interactions with its local environment, which is comprised primarily by the sugar-phosphate backbones.

We next turn to the homogeneous and inhomogeneous line width parameters of the (iCy3)_2_ dimer labeled +15 ‘duplex’ (Fig. 6*B*) and −1 ‘fork’ (Fig. 6*D*) ss-dsDNA constructs. These data show that the 2D spectral lineshapes of the (iCy3)_2_ dimer labeled duplex and fork ss-dsDNA constructs exhibit strikingly different and more complex temperature-dependent behavior than the iCy3 monomer labeled ss-dsDNA constructs discussed above. The results shown in Fig. 6*B* were obtained from our analysis of 2DFS data of the (iCy3)_2_ dimer labeled +15 duplex ss-dsDNA construct shown in Figs. 5*C* and 5*D* and Figs. S2 and S3 of the SI. In this case, both the homogeneous and inhomogeneous lineshape parameters undergo sensitive temperature-dependent variations. At the lowest temperature investigated (1°C), both the inhomogeneous and homogeneous line width parameters have the relatively low values, *σ*_*I*_ ∼122 cm^-1^ and Γ_*H*_ ∼105 cm^-1^, respectively. When the temperature is raised to 5°C, the inhomogeneous line width increases rapidly to *σ*_*I*_ ∼213 cm^-1^, followed by an abrupt decrease over the temperature range 5 – 23°C to the value *σ*_*I*_ ∼137 cm^-1^. The room temperature value of the inhomogeneous line width appears to be a local minimum, since increasing the temperature from 23 – 35°C leads to a rapid increase to the value, *σ*_*I*_ ∼253 cm^-1^, suggesting a dramatic increase of local conformational disorder just above room temperature. This relatively high value of the inhomogeneous line width parameter persists (*σ*_*I*_ >∼250 cm^-1^) over the temperature range 35 – 55°C, and through the melting point of the duplex region. In contrast, the homogeneous line width parameter increases over the range 1 – 23°C to its maximum value Γ_*H*_ ∼180 cm^-1^, followed by an abrupt decrease to Γ_*H*_ ∼92 cm^-1^ over the range 23 – 35°C. This relatively low value of the homogeneous line width persists (Γ_*H*_ < ∼70 cm^-1^) over the temperature range 35 – 55°C, and through the melting point of the duplex region.

We note that the homogenous and inhomogeneous line width parameters appear to depend on temperature in a reciprocal manner, with extremum values attained at room temperature. Evidently at room temperature, the local conformation of the (iCy3)_2_ dimer probes at the +15 position, which is deep within the duplex region of the ss-dsDNA construct, is minimally disordered such that the probes interact uniformly with their local environments to maximize the electronic dephasing rate. The room temperature condition appears to be unique. As the temperature is raised or lowered from 23°C, the local conformational disorder increases abruptly, while the mean coupling strength between the electronic transitions of the probe chromophores and the phonon bath decreases. These results suggest that the W-C local conformation of the (iCy3)_2_ dimer labeled sugar-phosphate backbones at sites deep within the duplex region is the optimally stable structure at 23°C, and that the distribution of conformations broadens substantially at temperatures just above or below room temperature. These findings indicate that the activation barriers for thermally induced breathing at these positions are readily surmounted just above room temperature. At the same time, decreasing the temperature below 23°C destabilizes the W-C local conformation in the duplex region, like ‘cold denaturation’ in proteins (43-45).

We next discuss the temperature-dependent lineshape parameters that we obtained for the (iCy3)_2_ dimer labeled −1 ‘fork’ ss-dsDNA construct, shown in Fig. 6*D*. At the lowest temperature (1°C), the inhomogeneous and homogeneous line widths have values *σ*_*I*_ ∼187 cm^-1^ and Γ_*H*_ ∼118 cm^-1^, respectively. Like the (iCy3)_2_ dimer labeled +15 duplex construct, the homogeneous and inhomogeneous line widths appear to vary with temperature in a reciprocal manner. However, in this case the principal variation occurs over the temperature range 1 – 15°C for which the inhomogeneous line width increases to *σ*_*I*_ ∼218 cm^-1^ and the homogeneous line width decreases to Γ_*H*_ ∼87 cm^-1^. Increasing the temperature to 23°C results in the inhomogeneous and homogeneous line width parameters recovering their low-temperature values: *σ*_*I*_ ∼187 cm^-1^ and Γ_*H*_ ∼114 cm^-1^, respectively. Upon further increasing the temperature, the inhomogeneous and homogeneous line width parameters do not undergo significant additional changes, suggesting that – unlike the duplex labeled ss-dsDNA constructs – the distribution of local conformations of the (iCy3)_2_ labeled fork construct is not broadened by thermally activated processes near room temperature.

### C. Local conformations and spectral line width parameters of (iCy3)_2_ dimer ss-dsDNA constructs labeled at the +2, +1, −1, and −2-positions

We next performed room temperature absorbance, CD and 2DFS experiments on (iCy3)_2_ dimer labeled ss-dsDNA constructs in which the dimer probe position was systematically varied across the ss-dsDNA fork junction. The results of these studies are summarized in Fig. 7, in which columns *A* – *D* correspond to the probe label positions: +2, +1, −1, and −2, respectively. The corresponding optimized values for the mean structural parameters and homogeneous and inhomogeneous line widths are listed in Table 4. Comparisons between the experimental and optimized simulated 2DFS data are presented in Figs. S18 – S21 of the SI, respectively. In the top two rows of Fig. 7, the experimental CD and absorbance spectra (green curves) are shown overlaid with simulated spectra resulting from our H-F model analyses. The color schemes are the same as those used in Fig. 4 above. Experimental and simulated 2DFS data using laser spectrum with FWHM 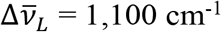 (Δ*λ*_*L*_= ∼33 nm, overlaid with absorbance spectra in gray) are shown in the third and fourth rows, respectively.

**Table 4.** Conformational parameters and 2D spectral line widths determined from H-F model analyses of absorbance, CD and 2DFS of (iCy3)_2_ dimer labeled ss-dsDNA fork constructs.

| Construct | $\bar{J}$ (cm<sup>-1</sup>) | $\bar{\phi}_{AB}$ (°) | $\bar{\theta}_{AB}$ (°) | $\bar{R}_{AB}$ (Å) | $\sigma_I$ (cm<sup>-1</sup>) | $\Gamma_H$ (cm<sup>-1</sup>) |
| --- | --- | --- | --- | --- | --- | --- |
| +2 | 443 | 84.2 ± 0.1 | 6.6 ± 32 | 2.8 ± 0.1 | 167.1 +21.8/-16.6 | 96.2 +14.8/-10 |
| +1 | 426 | 77.0 ± 0.3 | 48.5 ± 3.3 | 5.8 ± 0.1 | 212.6 +28.2/-22.8 | 100.6 +18.2/-12.9 |
| -1 | -367 | 96.0 ± 0.2 | 26.4 ± 2.5 | 4.3 ± 0.1 | 202.5 +20.3/-18.0 | 82.9 +11.2/-12.4 |
| -2 | -420 | 97.3 ± 1.8 | 27.1 ± 54 | 4.4 ± 0.7 | 192.4 +20.0/-16.2 | 87.3 +13.6/-9.6 |

**Figure 7.**
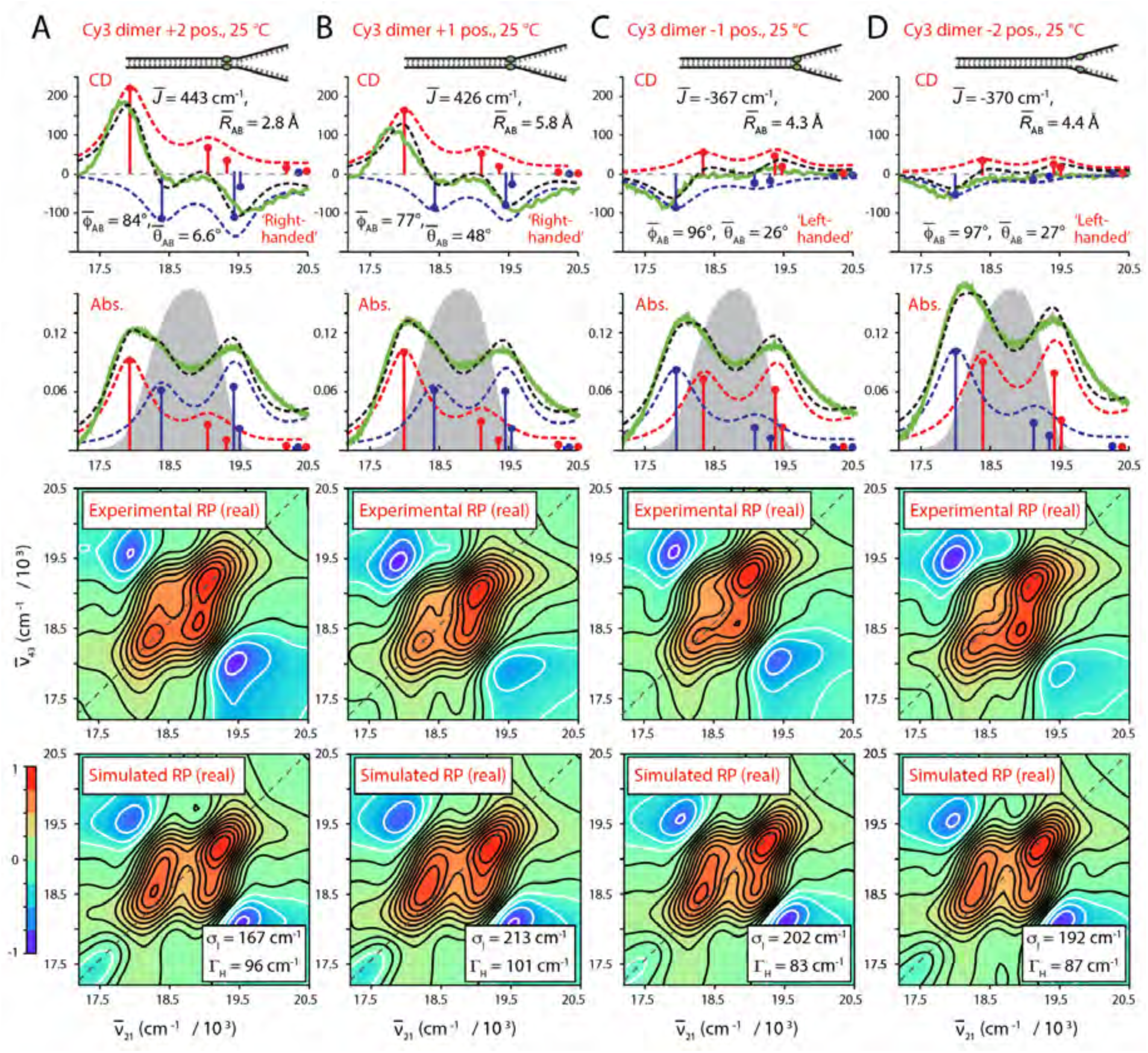
Experimental and simulated spectroscopic measurements performed at room temperature (25°C) for (iCy3)_2_ dimer ss-dsDNA constructs as a function of probe labeling position. (***A***) +2; (***B***) +1; (***C***) −1; and (***D***) −2. The experimental CD (top row) and absorbance spectra (second row) are shown (in green) overlaid with vibronic spectral features (black dashed curves) obtained from optimized fits to the H-F model. The symmetric (+) and anti-symmetric (–) excitons are shown in blue and red, respectively. Values of the optimized parameters are shown in the insets of the corresponding panels. The laser spectrum (in gray) is shown overlaid with the absorbance spectrum and is the same as in Fig. 4. Experimental RP spectra (third row) are compared to the optimized simulated RP spectra (fourth row). Simulated spectra are based on the structural parameters obtained from our optimization analyses of the CD and absorbance spectra, the fluorescence quantum yield parameter Γ_2*D*_= 0.3, and the homogeneous and inhomogeneous line width parameters (Γ_*H*_ and *σ*_*I*_, respectively) listed in Table 4. Comparisons between experimental and optimized simulated 2DFS data are presented in Figs. S18 – S21 of the SI.

From our H-F model analyses of the absorbance and CD data, we see that the mean electrostatic coupling 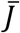 undergoes a sign inversion as the (iCy3)_2_ dimer probe position is changed from the +1 to −1 position across the ss-dsDNA junction. This is accompanied by non-continuous changes of the local conformational coordinates: the mean interchromophore separation 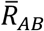 [2.8 Å (+2), 5.8 Å (+1), 4.3 Å (−1) and 4.4 Å (−2)], the mean twist angle 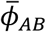 [84º (+2), 77º (+1), 96º (−1) and 97º (−2)], and the mean tilt angle 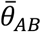 [6.6º (+2), 48º (+1), 26º (−1) and 27 (−2)]. The sign inversion of the electrostatic coupling between the +1 and −1 positions indicates that the conformation of the (iCy3)_2_ dimer probe, and presumably that of the sugar-phosphate backbones labeled at these sites, changes from right-handed to left-handed. This change of handedness is correlated to a change of the mean twist angle 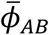 from values that are less than 90º (right-handed) to values that are greater than 90º (left-handed). In addition, the mean tilt angle 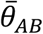 undergoes an abrupt increase from 7º to 48º between the +2 and +1 positions, followed by a decrease to ∼27º at the −1 and −2 positions. This indicates that the local conformation of the sugar-phosphate backbones within the ss-dsDNA fork constructs undergo an abrupt loss of cylindrical symmetry at the interface between the +2 and +1 positions. However, some of the cylindrical symmetry is recovered at the −1 and −2 positions.

Our analyses of the 2DFS spectral lineshapes show how local conformational disorder of the (iCy3)_2_ dimer probes, in addition to interactions between the electronic transitions and the phonon bath, depend on the probe labeling position. In Fig. 8*A*, we plot the position-dependent values of the inhomogeneous and homogeneous line width parameters. For the +2 ss-dsDNA fork construct, the inhomogeneous line width *σ*_*I*_ ∼167 cm^-1^, and the homogeneous line width Γ_*H*_ ∼96 cm^-1^. The value for the inhomogeneous line width is significantly larger than the one we obtained for the +15 ss-dsDNA construct (*σ*_*I*_ ∼137 cm^-1^), while the value for the homogeneous line width is smaller (compare to 23ºC point of Fig. 6*B*), indicating a higher degree of conformational disorder at the +2 position relative to +15. When the probe labeling position is changed to +1, the inhomogeneous line width parameter increases: *σ*_*I*_ ∼213 cm^-1^, while the homogeneous line width undergoes only a slight increase: Γ_*H*_ ∼101 cm^-1^. This finding suggests that although the sugar-phosphate backbones at the +1 position maintain the right-handed local conformations seen at the +2 and +15 positions (characteristic of the B-form double-helix), the distribution of local conformations is further broadened at the +1 position in comparison to the +2 position. When the probe labeling position is changed to −1, we see that both the inhomogeneous and homogeneous line width parameters decrease to the values *σ*_*I*_ ∼202 cm^-1^ and Γ_*H*_ ∼83 cm^-1^. These values do not change significantly when the probe labeling sites are shifted to the −2 position: *σ*_*I*_ ∼192 cm^-1^ and Γ_*H*_ ∼87 cm^-1^. These findings suggest that the distribution of local conformations decrease slightly at the −1 and −2 positions relative to the +1 position. Thus, the abrupt change in average local conformation that we observe across the +2 to +1 ss-dsDNA junction (from right-handed cylindrically symmetric to right-handed cylindrically asymmetric) is accompanied by the appearance of local conformational disorder at the +1 position. An additional change in the average local conformation occurs across the +1 to −1 positions (from right-handed to left-handed) for which the local conformational disorder persists for positions extending into the single-stranded region of the ss-dsDNA constructs.

**Figure 8.**
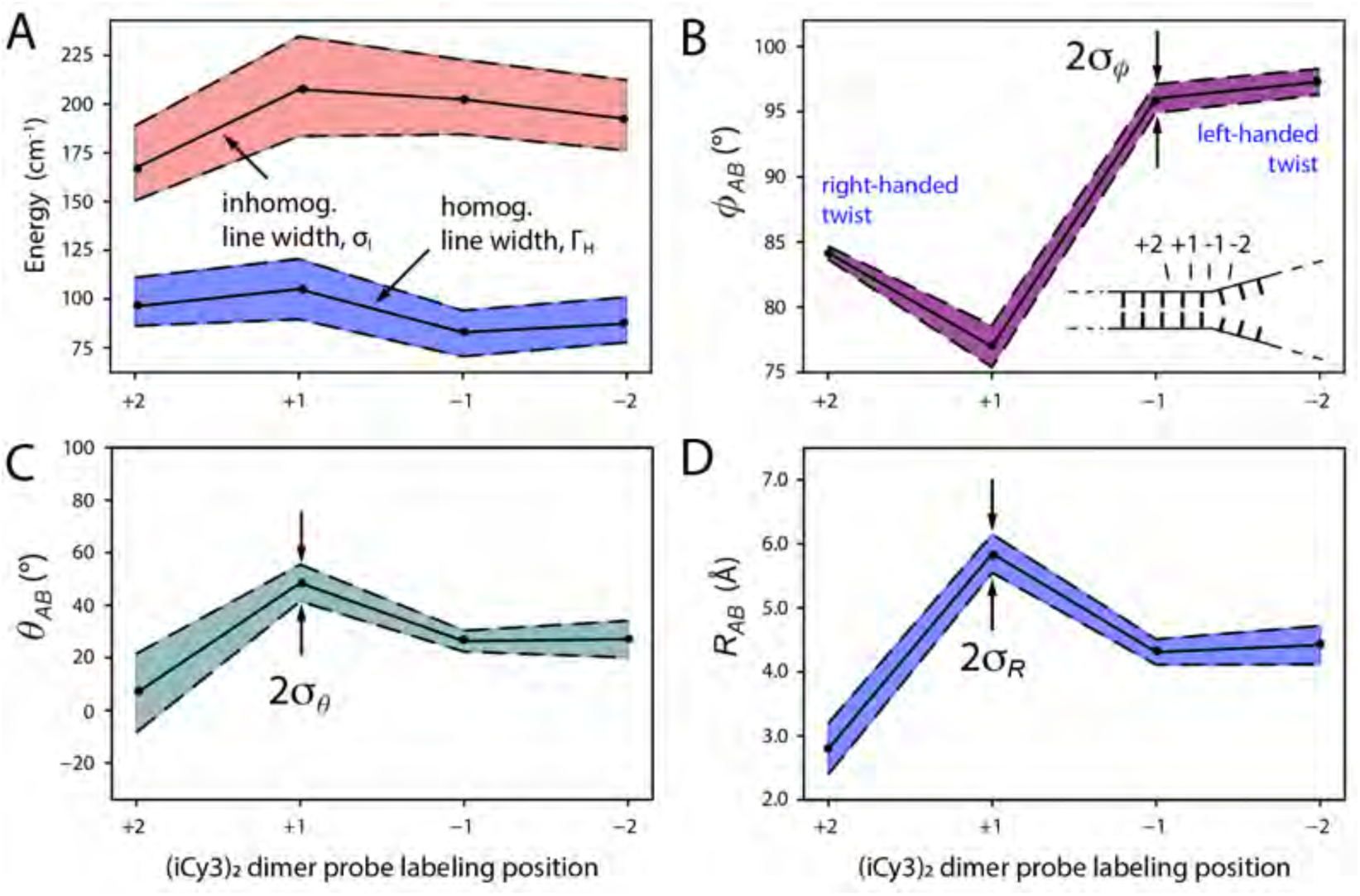
Optimized spectral line width and structural parameters of (iCy3)_2_ dimer labeled ss-dsDNA constructs for varying label position obtained from 2DFS lineshape analyses. (***A***) homogeneous and inhomogeneous line width parameters. (***B***) Mean twist angle 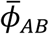; (***C***) mean tilt angle 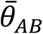; and (***D***) mean inter-chromophore separation 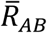. In panel ***A***, the shaded regions bounded by dashed lines indicate error bars, which were calculated based on a 1% deviation of the 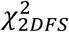 function [Eq. (20)] from its minimized value. In panels (***B*** – ***D***), the shaded regions bounded by dashed lines indicate the standard deviations of the structural parameter distributions obtained from analyses of the 2DFS spectral lineshapes, which are listed in Table 5. These values were obtained by minimizing the least squares error functions shown in Fig. S23 of the SI.

### D. Distribution of local conformational parameters of (iCy3)_2_ dimer ss-dsDNA constructs labeled at the +2, +1, −1, and −2-positions

We used the 2DFS data shown in Fig. 7 and Figs. S18 – S22 of the SI to model the distributions of conformational parameters according to the method outlined in Sect. 3D. The results of these studies are summarized in Table 5 and are shown together with the optimized values for the inhomogeneous and homogeneous line width parameters in Fig. 8. In Figs. 8*B* – 8*D*, the optimized values for the mean conformational parameters, which we determined from our analyses of absorbance and CD spectra, are presented as points. Shaded regions represent the optimized Gaussian widths of the corresponding distributions, which we determined by minimizing the least squares error functions 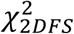 shown in Fig. S23 of the SI. In Fig. 8*A*, the inhomogeneous and homogeneous line width parameters and 1% deviation error bars are presented, which we determined according to the 2D lineshape analysis presented in previous sections.

**Table 5.** Standard deviations of conformational parameters determined from H-F model analyses of absorbance, CD and 2DFS data of (iCy3)_2_ dimer labeled ss-dsDNA fork constructs. The optimized values were obtained by minimizing the least squares error function 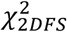 described in Sect. 3D, which are shown in Fig. S23 of the SI.

| Construct | $\sigma_\phi$ (°) | $\sigma_\theta$ (°) | $\sigma_R$ (Å) |
| --- | --- | --- | --- |
| +2 | 0.8 | 17 | 0.4 |
| +1 | 1.8 | 7.0 | 0.28 |
| -1 | 1.1 | 4.0 | 0.22 |
| -2 | 1.0 | 5.5 | 0.32 |

In Fig. 8*B*, we plot the position-dependence of the mean twist angle 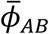 and standard deviation *σ*_*ϕ*_ For the +2 ss-dsDNA construct, the mean twist angle has the value 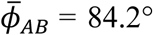, which is very close to that obtained for the +15 position (79.6°). The value we determined for the standard deviation at this position is relatively small, *σ*_*ϕ*_ = ∼0.7°. As the position is changed to +1, the twist angle decreases to 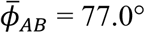 with standard deviation *σ*_*ϕ*_ = ∼1.7°. When the position is changed to −1, the mean twist angle undergoes a significant increase to 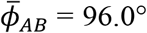 (from right-handed to left-handed) with standard deviation *σ*_*ϕ*_ = ∼1.1°. The values for both the mean twist angle and standard deviation do not change significantly when the probe labeling position is changed from −1 to −2. From these results, we conclude that while the mean twist angle undergoes significant changes as the probe labeling position is varied across the ss-dsDNA fork junction, the distribution of twist angles remains relatively narrow (*σ*_*ϕ*_ < ∼2°) for these positions.

We next consider the position-dependent behavior of the mean tilt-angle 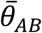 and its standard deviation *σ*_*θ*_, as shown in Fig. 8*C*. For the +2 labeled ss-dsDNA construct, the mean tilt angle is 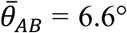 and the standard deviation is *σ*_*θ*_= ∼17°, indicating that there is a significant degree of conformational disorder in the tilt angle parameter at this position, although the sugar-phosphate backbones occupy, on average, a cylindrically symmetric local conformation. When the label position is changed to +1, the mean tilt angle increases dramatically to 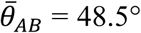 and the standard deviation decreases slightly to *σ*_*θ*_= ∼7°. This indicates that the cylindrical symmetry of the sugar-phosphate backbones, which is normally found in the duplex region, is no longer present at the +1 position and that the conformational disorder of the tilt angle has decreased slightly. When the label position is changed to −1, the mean tilt angle decreases significantly to 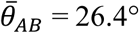 and the standard deviation decreases to *σ*_*θ*_ = ∼3.5°. When the label position is changed from −1 to −2, the mean tilt angle does not change significantly 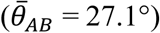, although the standard deviation increases to *σ*_*θ*_= ∼6°.

In Fig. 8*D*, we show the position-dependent behavior of the mean interchromophore separation 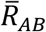. For the +2 labeled construct, the mean separation is 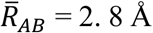 and the standard deviation is *σ*_*R*_ = ∼0.4 Å. We acknowledge that this value for the mean separation is unrealistically small as it lies below the van der Walls contact distance of ∼3.5 Å. This inconsistency is likely due to a breakdown of our H-F model analysis that underestimates the coupling strength 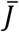 for small interchromophore separations (11). When the probe label position is changed to +1, the mean separation decreases slightly to 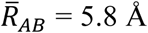 and the standard deviation decreases slightly to *σ*_*R*_ = ∼0.3 Å. However, when the position is changed from +1 to −1, the mean separation decreases significantly to 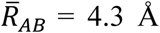 and the standard deviation decreases to *σ*_*R*_ = ∼0.2 Å. When the position is changed further from −1 to −2, the mean separation does not change significantly 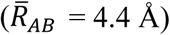 and the standard deviation is relatively unchanged: *σ*_*R*_ = ∼0.2 Å.

It is interesting to compare our results for the distributions of local conformational parameters, as summarized in Figs. 8*B* – 8*D*, to the optimized values we obtained for the homogeneous and inhomogeneous line widths shown in Fig. 8*A*. The position-dependencies of the line width parameters were discussed in the previous section and show that there is relatively little variation in the inhomogeneous line width *σ*_*I*_ as the probe label position is changed from the +2 to −2 positions across the ss-dsDNA junction. Although the mean structural coordinates vary sensitively across the ss-dsDNA junction, the standard deviations of the structural coordinates that we obtained from our 2DFS lineshape analysis indicate that the distributions of conformational coordinates at these positions remain relatively narrow at room temperature.

## 5. Conclusions

The conservation of base sequence integrity within genomic DNA is critical to maintaining well-regulated gene expression and replication. However, the structure of DNA within the cell must be dynamic, allowing for thermally induced fluctuations (i.e., DNA ‘breathing’) to facilitate productive interactions, including binding-site recognition and the assembly of functional protein-DNA complexes. For example, at ss-dsDNA fork junctions, transient local conformation fluctuations of the sugar-phosphate backbones are likely transition states for the formation of a stable helicase-primase (primosome) sub-assembly during DNA replication, with the existence of multiple DNA conformers working to facilitate competition between different protein regulatory factors and replisome proteins.

In this work, we have probed the average local conformations and the degree of conformational disorder at and near model ss-dsDNA replication fork junctions through site-specific internal labeling with two cyanine dyes (iCy3) rigidly inserted within the sugar-phosphate backbones at opposite positions within complementary single strands. We performed linear (absorbance and CD) and nonlinear (2DFS) spectroscopic studies of iCy3 monomer and (iCy3)_2_ dimer labeled ss-dsDNA constructs as a function of temperature and probe label position (see Table 1). Our analyses of the absorbance spectra of the iCy3 monomer labeled ss-dsDNA constructs indicate that the monomer Hamiltonian parameters (i.e., the mean electronic transition energy 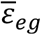, the Huang-Rhys vibronic coupling parameter *λ*^2^, and the vibrational frequency *ω*_0_) are largely insensitive to temperature and probe label position (see Table S1 and Table S2 of the SI). Conversely, the absorbance and CD spectra of (Cy3)_2_ dimer labeled ss-dsDNA constructs respond sensitively to changing temperature and probe label position, from which we observed systematic changes of the optimized dimer Hamiltonian parameters (i.e., the mean resonant coupling 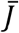, the mean twist angle 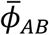, the mean tilt angle 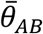, and the mean interchromophore separation 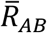, see Tables 2 – 4). Our 2DFS experiments provide information about the local conformational disorder of the Cy3 probes at the probe label positions, from which we found that the Cy3 monomer labeled ss-dsDNA constructs are significantly more disordered than the corresponding (Cy3)_2_ dimer labeled constructs at room temperature (see Fig. 6). The relatively high disorder that we observed in the iCy3 monomer labeled ss-dsDNA constructs is likely due to a mismatch of the W-C base pairing in the vicinity of the probe labeling site. In the iCy3 monomer labeled ss-dsDNA constructs, a single thymine (T) base was positioned directly opposite to the iCy3 probe on the conjugate strand to act as a spacer. Nevertheless, normal W-C pairing is likely perturbed by the incorrect spacing between complementary bases introduced by the presence of the single iCy3 monomer probe. In contrast, we found that the (iCy3)_2_ dimer labeled ss-dsDNA constructs are minimally disordered at room temperature and physiological buffer salt conditions, and their conformation-dependent spectroscopic properties appear to represent the site-specific local conformations of the sugar-phosphate backbones at positions relative to the ss-dsDNA fork junction.

From our temperature-dependent studies of the (iCy3)_2_ dimer labeled ss-dsDNA +15 ‘duplex’ construct, we found that local conformations of the sugar-phosphate backbones deep within the double-strand region occupy a minimally disordered, right-handed B-form conformation at room temperature (23°C). The B-form conformation is destabilized when the temperature is either raised or lowered from room temperature (see Fig. 6*B* and Table 2). These observations are consistent with the notion that the room temperature stability of the B-form conformation results from a nearly equal balance between opposing thermodynamic forces, and that small departures from room temperature (in either the positive or negative direction) alters the free energy landscape to populate non-B-form conformations. Such a picture is sometimes invoked in ‘DNA breathing and trapping models,’ in which non-canonical local conformations of the DNA framework are transiently populated under physiological conditions and function as activated states for protein-DNA complex assembly (2, 35, 46).

Our temperature-dependent studies of the (iCy3)_2_ dimer labeled ss-dsDNA −1 ‘fork’ construct showed that the average local conformation of the sugar-phosphate backbones at the probe position is left-handed and relatively disordered (in comparison to the +15 position) at room temperature. Increasing the temperature above 23°C did not significantly change the average local conformation or conformational disorder at the −1 position, suggesting that the free energy landscape is insensitive to increasing temperature. However, like our observation for the +15 duplex ss-dsDNA construct, decreasing temperature below 23°C did lead to a significant increase of the conformational disorder at the −1 position. These observations suggest, as has been studied in protein systems, that the concept of ‘cold denaturation’ (defined as positions in the phase diagram for the folding-unfolding transition of the protein where changes in temperature in either direction decreases the stability of the folded form) might be productively applied to investigations of the stability of ss-dsDNA transitions as well (44, 45, 47-49). Future studies of nucleic acid stability, using the 2DFS approach described in the current work to investigate conformational disorder at nucleic acid positions of possible physiological interest, may help to shed new light on the underlying molecular mechanisms of some of the central processes of genome expression.

The results of our position-dependent studies of the (iCy3)_2_ dimer labeled ss-dsDNA constructs at room temperature provide detailed information about the local conformations of the sugar-phosphate backbones at positions across the ss-dsDNA fork junction and are summarized in Fig. 9. The mean local conformations of the sugar-phosphate backbones are right-handed (with mean twist angle, 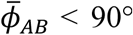) for positive integer positions, and left-handed 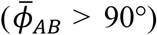 for negative integer positions. Local conformations deep within the duplex region are cylindrically symmetric (with mean tilt angle 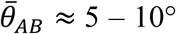) and minimally disordered (with inhomogeneous line width, *σ*_*I*_ = 137 cm^-1^). The disorder increases significantly for positive positions approaching the ss-dsDNA fork junction (*σ*_*I*_ = 167 cm^-1^ at the +2 position). At the +1 position, we observe an abrupt loss of cylindrical symmetry 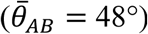, which coincides with an additional gain in conformational disorder (*σ*_*I*_ = 213 cm^-1^). The left-handed conformations at the −1 and −2 positions exhibit somewhat smaller mean tilt angles (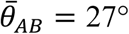 and 26°, respectively), and decreasing conformational disorder (*σ*_*I*_ = 202 cm^-1^ and 192 cm^-1^, respectively), suggesting that the peak perturbation to secondary structure within the ss-dsDNA fork junction occurs at the +1 position.

**Figure 9.**
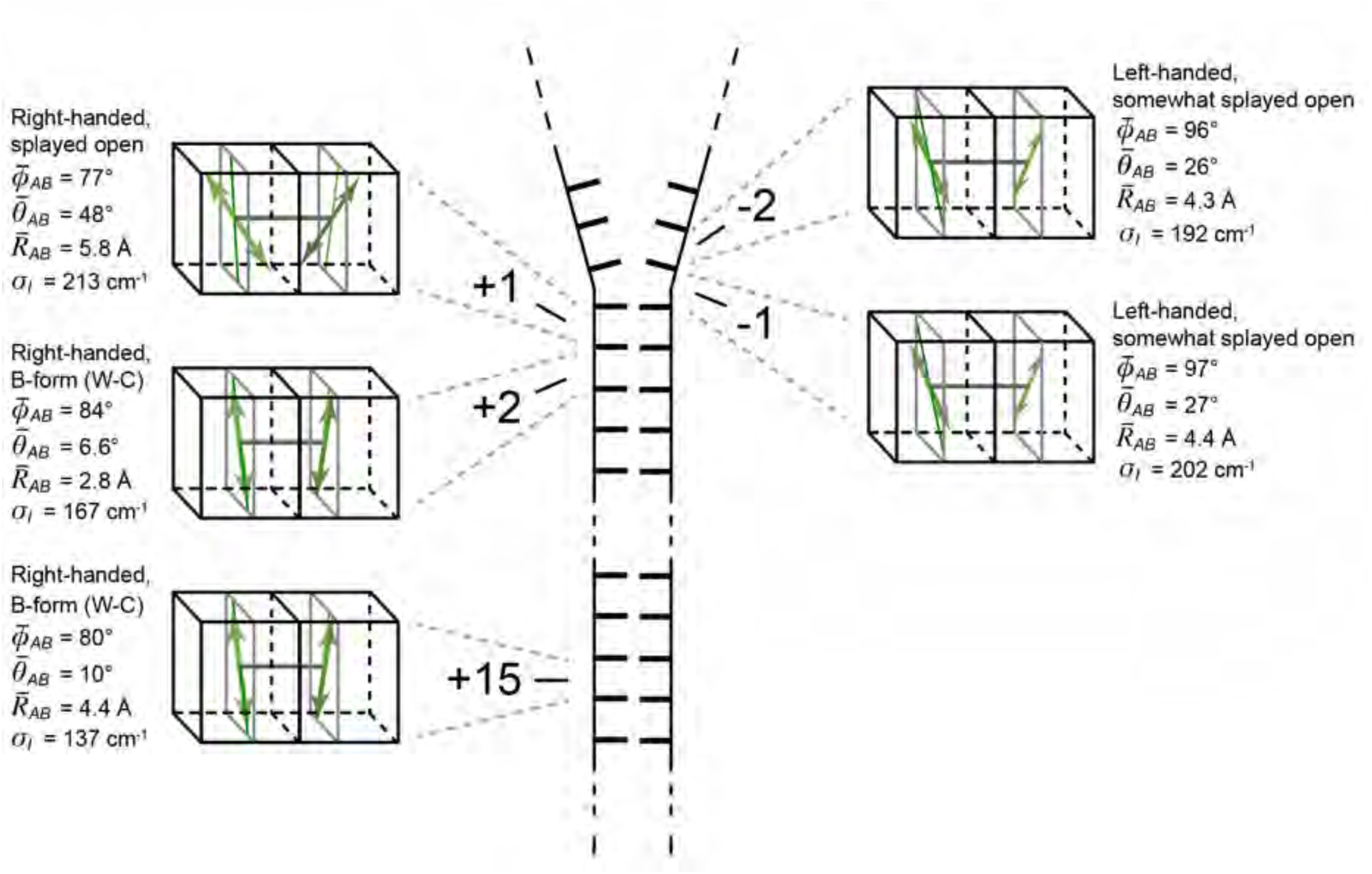
Schematic illustration of the average local conformations of the (iCy3)_2_ dimer labeled ss-dsDNA junction at positions +15, +2, +1, −1 and −2 at room temperature.

Our 2DFS experiments provide additional information about the standard deviations of the distributions of conformational coordinate (see Fig. 8 and Table 5). Perhaps surprisingly, our analyses indicate that the distributions of conformational parameters at positions traversing the ss-dsDNA fork junction are narrow, suggesting that regions of the junction extending into the single strands are relatively well-ordered. We emphasize that while the above conclusions are based on the interpretation of ensemble spectroscopic measurements, future single-molecule experiments performed on (iCy3)_2_ dimer ss-ds DNA constructs can, in principle, probe directly the individual conformational states that underlie the distributions reported here.

Our findings provide a detailed picture of the variation of local conformation and conformational disorder of the sugar-phosphate backbones at and near the ss-dsDNA fork junction. The relatively narrow distributions of local conformations at key positions at and near DNA junctions implies that the number of possible states that mediate protein binding may be rather limited and suggests a possible structural framework for understanding the roles of transient DNA junction conformations in driving the processes of DNA-protein complex assembly and function.

## Acknowledgements

The authors are grateful to our laboratory colleagues in the Marcus and von Hippel groups for many helpful discussions. This work was supported by grants from the National Institutes of Health General Medical Sciences (GM-15792 to A.H.M. and P.H.v.H.) and the National Science Foundation Chemistry of Life Processes Program (CHE-1608915 to A.H.M.). P.H.v.H. is an American Cancer Society Research Professor of Chemistry.

## Supporting Information

**Figure S1.**
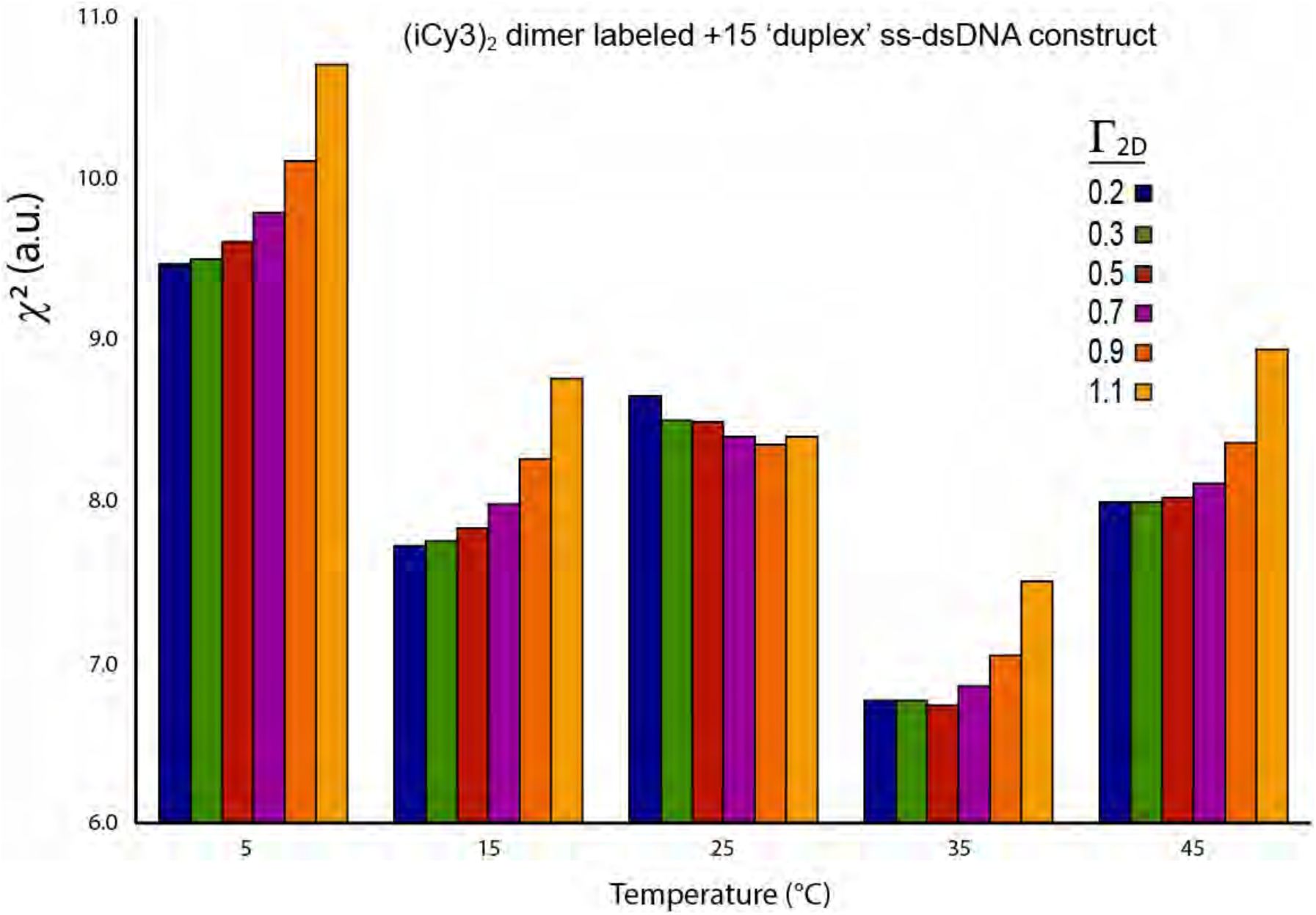
Temperature-dependent *χ*^2^ parameter resulting from optimizations of the dimer Hamiltonian given by Eq. (4) of the main text to experimental 2DFS data of the (iCy3)_2_ +15 ‘duplex’ labeled ss-dsDNA construct.

**Table S1.** Temperature-dependent Hamiltonian parameters and 2DFS line widths determined for the iCy3 monomer labeled +15 ‘duplex’ ss-dsDNA construct. The parameters determined from absorbance and CD are the electronic transition energy, *ε*_*eg*_, the vibrational frequency, *ω*_0_, and the Huang-Rhys electronic-vibrational coupling parameter, *λ*^2^. The parameters determined from 2DFS data are the inhomogeneous and homogeneous line widths, *σ*_*I*_ and Γ_*H*_. Error bars were calculated based on a 1% deviation of the *χ*^2^function from its minimum (optimized) value.

| Optimized Parameters for iCy3 Monomer +15 ‘Duplex’ ss-dsDNA Constructs |  |  |  |  |  |
| --- | --- | --- | --- | --- | --- |
| From Absorbance and CD Optimization |  |  |  | From 2DFS Optimization |  |
| $T$ (°C) | $\varepsilon_{eg}$ (cm<sup>-1</sup>) | $\omega_0$ (cm<sup>-1</sup>) | $\lambda^2$ | $\sigma_I$ (cm<sup>-1</sup>) | $\Gamma_H$ (cm<sup>-1</sup>) |
| 1 | 18,285 +40/-39 | 1,116 +99/-103 | 0.54 +0.07/-0.06 | 157.0 +40.2/-37.94 | 189.2 +29.3/-27.3 |
| 5 | 18,285 +40/-39 | 1,116 +99/-103 | 0.54 +0.07/-0.06 | 202.5 +26.3/-21.31 | 149.4 +17.2/-12.7 |
| 15 | 18,285 +40/-39 | 1,116 +99/-103 | 0.54 +0.07/-0.06 | 197.5 +20.8/-22.3 | 131.6 +12.7/-13.8 |
| 25 | 18,277 +38/-37 | 1,109 +88/-90 | 0.56 +0.06/-0.06 | 217.7 +24.6/-29.1 | 144.9 +20.4/-16.2 |
| 35 | 18,266 +36/-35 | 1,119 +82/-84 | 0.56 +0.06/-0.06 | 192.4 +24.2/-17.3 | 140.5 +14.6/-12 |
| 45 | 18,262 +36/-35 | 1,113 +82/-82 | 0.56 +0.06/-0.05 | 202.5 +29.8/-25.4 | 131.6 +18.3/-15 |
| 55 | 18,280 +39/-38 | 1,124 +93/-96 | 0.55 +0.06/-0.06 | 192.4 +22.3/-19.9 | 105.1 +15/-10.4 |
| 65 | 18,301 +45/-45 | 1,103 +103/-107 | 0.54 +0.07/-0.07 | 202.5 +24.8/-25.1 | 113.9 +15.1/-13.5 |

**Table S2.** Temperature-dependent Hamiltonian parameters and 2DFS line widths determined for the iCy3 monomer labeled −1 ‘fork’ ss-dsDNA construct. The parameters listed are the same as those described in Table S1. Error bars were calculated based on a 1% deviation of the *χ*^2^ function from its minimum (optimized) value.

| Optimized Parameters for iCy3 Monomer -1 ‘Fork’ ss-dsDNA Constructs |  |  |  |  |  |
| --- | --- | --- | --- | --- | --- |
| From Absorbance and CD Optimization |  |  |  | From 2DFS Optimization |  |
| $T$ (°C) | $\epsilon_{eg}$ (cm<sup>-1</sup>) | $\omega_0$ (cm<sup>-1</sup>) | $\lambda^2$ | $\sigma_I$ (cm<sup>-1</sup>) | $\Gamma_H$ (cm<sup>-1</sup>) |
| 1 | 18,223 +37/-36 | 1,112 +83/-85 | 0.58 +0.06/-0.06 | 202.5 +22/-16.5 | 127.2 +11.8/-10.9 |
| 5 | 18,223 +37/-36 | 1,112 +83/-85 | 0.58 +0.06/-0.06 | 202.5 +22.5/-12.3 | 131.6 +12/-12.2 |
| 15 | 18,223 +37/-36 | 1,112 +83/-85 | 0.58 +0.06/-0.06 | 192.4 +20.8/-19.2 | 118.4 +11.4/-12.7 |
| 25 | 18,240 +37/-35 | 1,116 +81/-82 | 0.59 +0.06/-0.06 | 197.5 +26.5/-23.2 | 105.1 +15.4/-12.8 |
| 35 | 18,252 +36/-35 | 1,105 +80/-80 | 0.58 +0.06/-0.06 | 192.4 +21/-17 | 109.5 +13.7/-9.3 |
| 45 | 18,249 +37/-35 | 1,088 +79/-79 | 0.59 +0.06/-0.06 | 192.4 +18.3/-18.3 | 122.8 +10.6/-12 |
| 55 | 18,257 +39/-38 | 1,086 +83/-83 | 0.58 +0.06/-0.06 | 187.3 +17.9/-17.9 | 140.5 +12.1/-11.4 |
| 65 | 18,263 +38/-37 | 1,074 +83/-83 | 0.58 +0.06/-0.05 | 202.5 +26.1/-24.1 | 113.9 +14.2/-14.6 |

**Figure S2.**
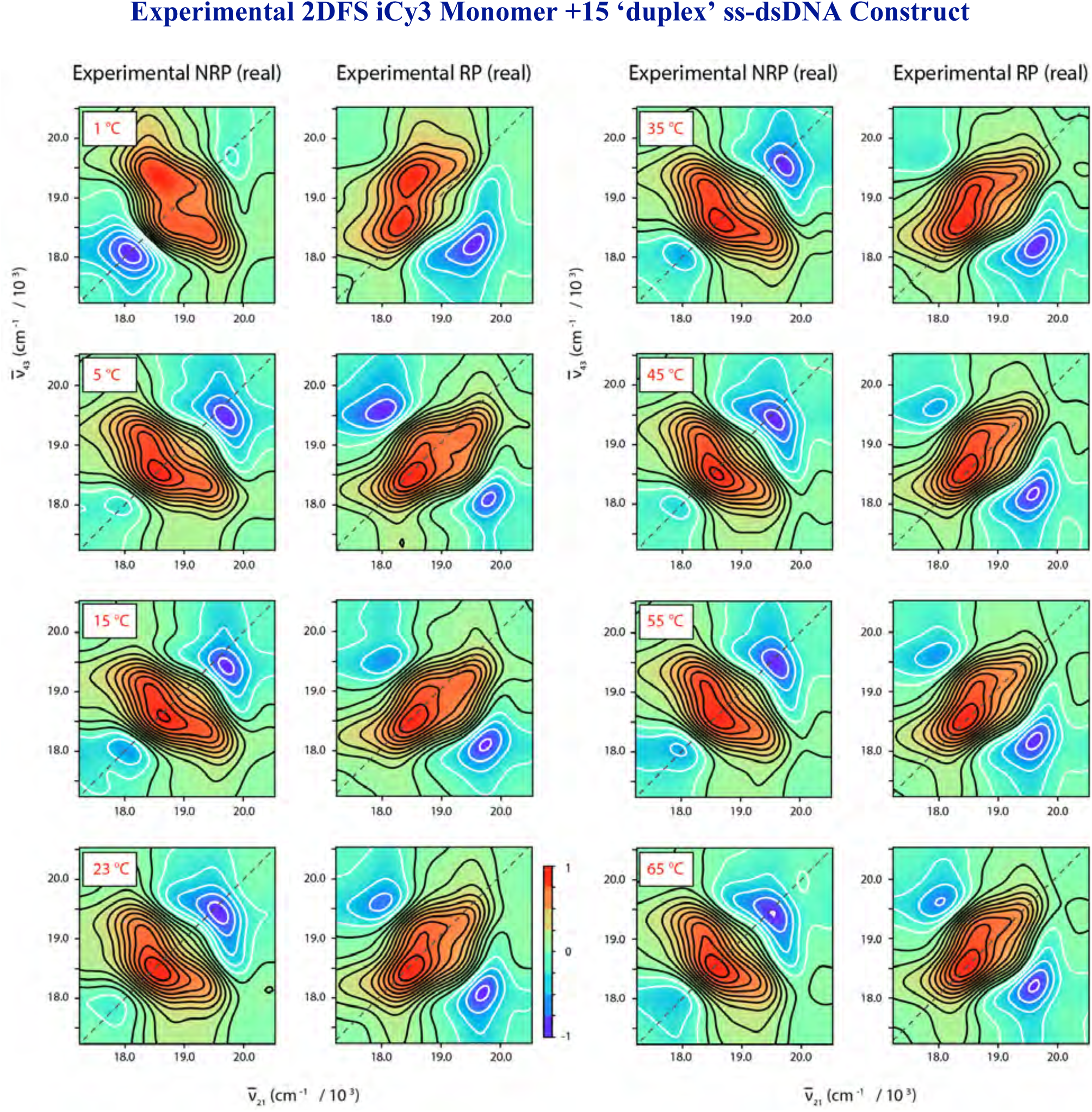
Temperature-dependent experimental NRP and RP 2DFS data (real part) for the iCy3 monomer labeled +15 ‘duplex’ ss-dsDNA construct. The laser spectrum used for these measurements is shown in Fig. 4 of the main text. The melting temperature of the duplex region of the construct is ∼65 °C.

**Figure S3.**
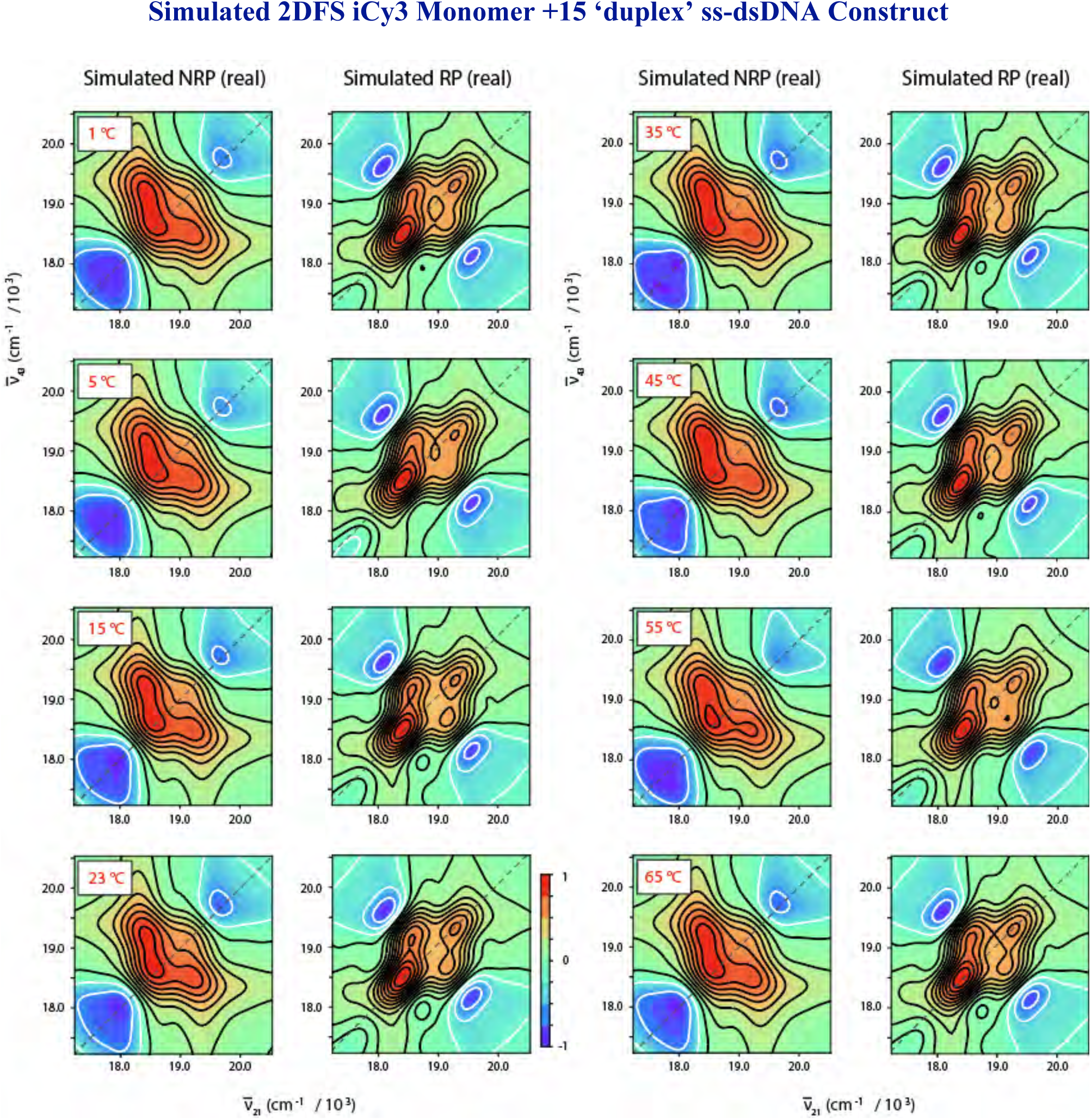
Temperature-dependent simulated NRP and RP 2DFS data (real part) for the iCy3 monomer labeled +15 ‘duplex’ ss-dsDNA construct. Simulations were performed using the monomer Hamiltonian given by Eq. (1) of the main text. The monomer Hamiltonian parameters used as input, in addition to the optimized homogeneous and inhomogeneous spectral line widths determined from our analyses, are listed in Table S1.

**Figure S4.**
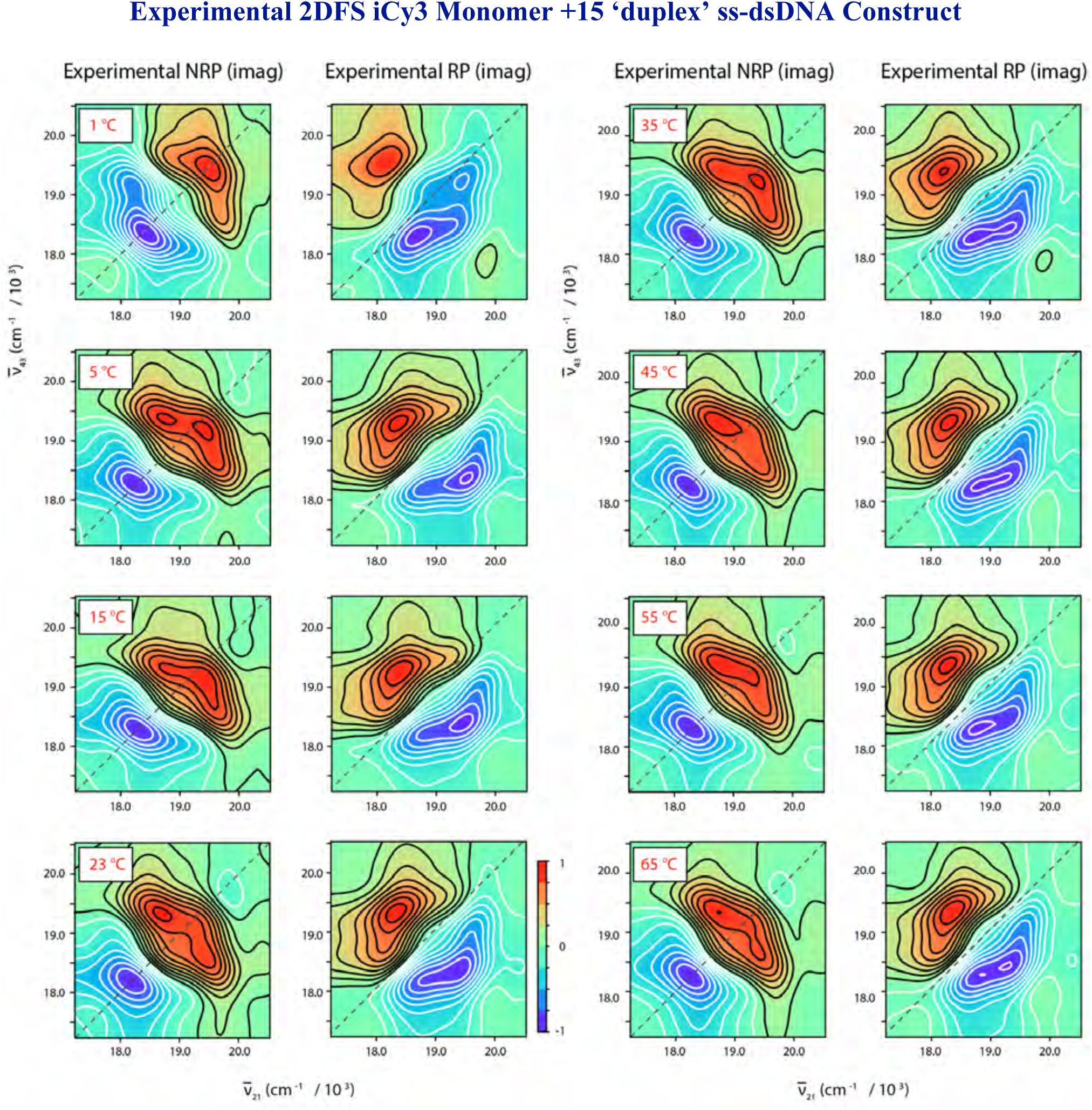
Temperature-dependent experimental NRP and RP 2DFS data (imaginary part) for the iCy3 monomer labeled +15 ‘duplex’ ss-dsDNA construct. The laser spectrum used for these measurements is shown in Fig. 4 of the main text.

**Figure S5.**
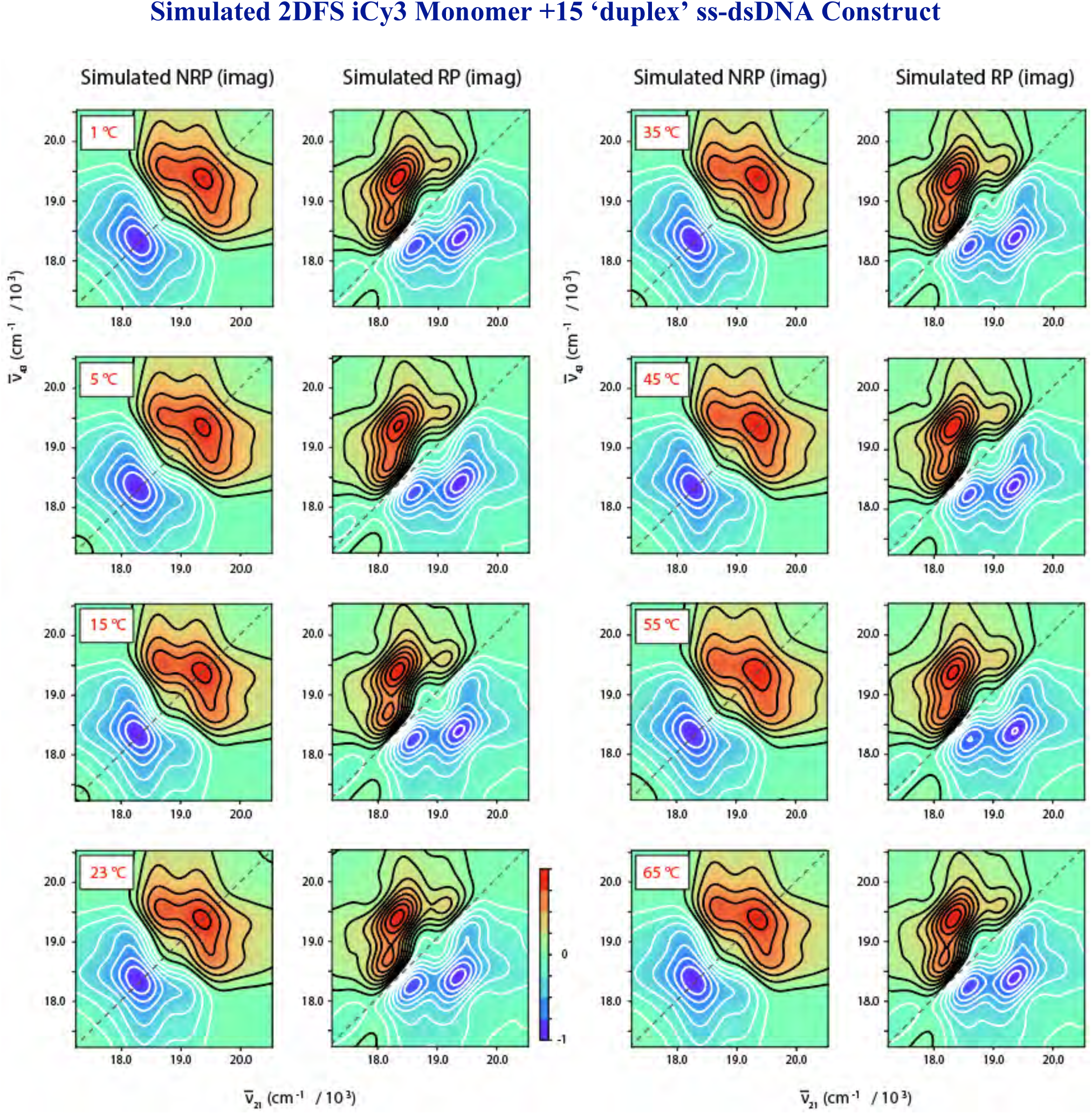
Temperature-dependent simulated NRP and RP 2DFS data (imaginary part) for the iCy3 monomer labeled +15 ‘duplex’ ss-dsDNA construct. Simulations were performed using the monomer Hamiltonian given by Eq. (1) of the main text. The monomer Hamiltonian parameters used as input, in addition to the optimized homogeneous and inhomogeneous spectral line widths determined from our analyses, are listed in Table S1.

**Figure S6.**
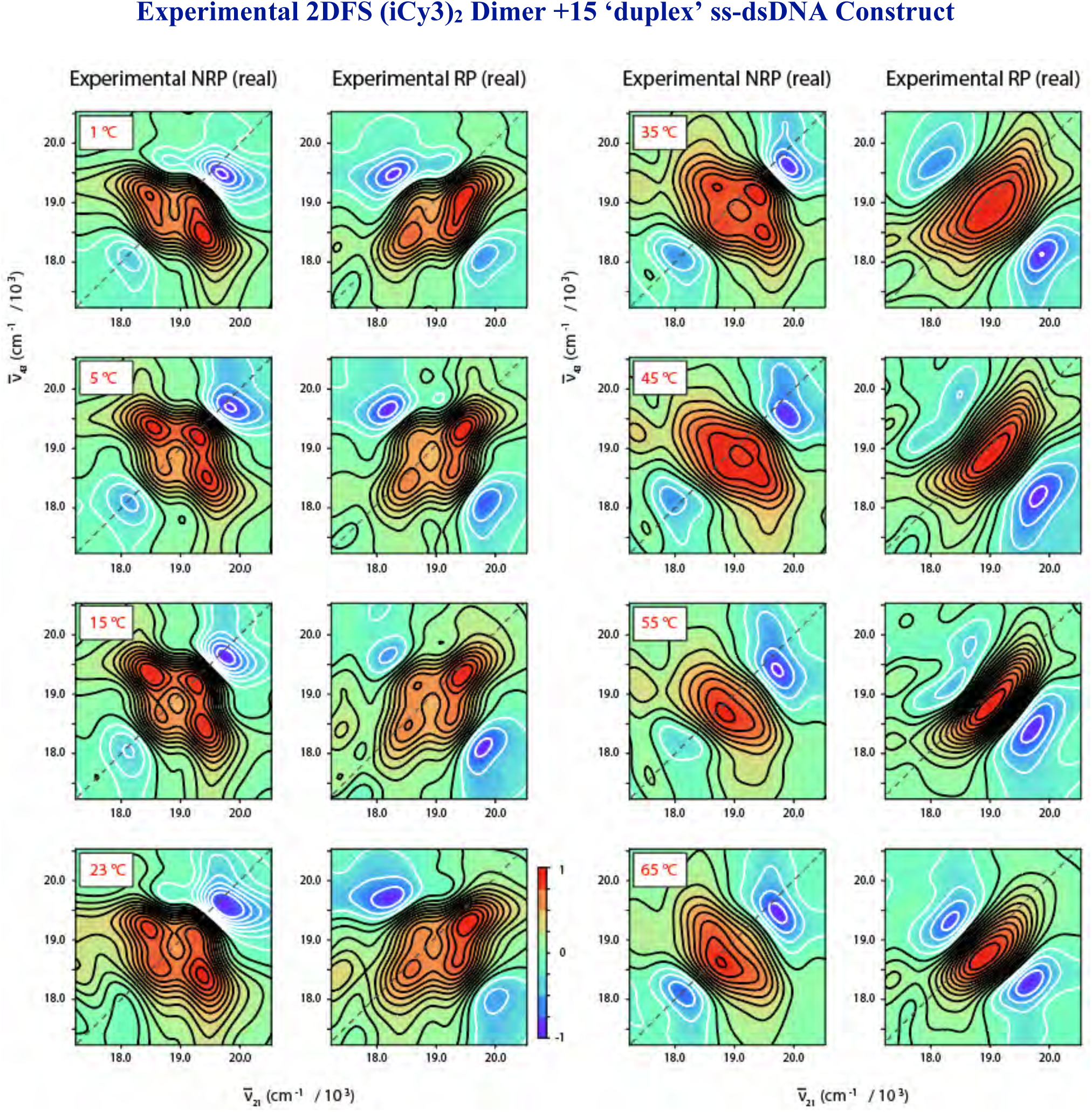
Temperature-dependent experimental NRP and RP 2DFS data (real part) for the (iCy3)_2_ dimer labeled +15 ‘duplex’ ss-dsDNA construct. The laser spectrum used for these measurements is shown in Fig. 4 of the main text. The melting temperature of the duplex region of the construct is ∼65 °C.

**Figure S7.**
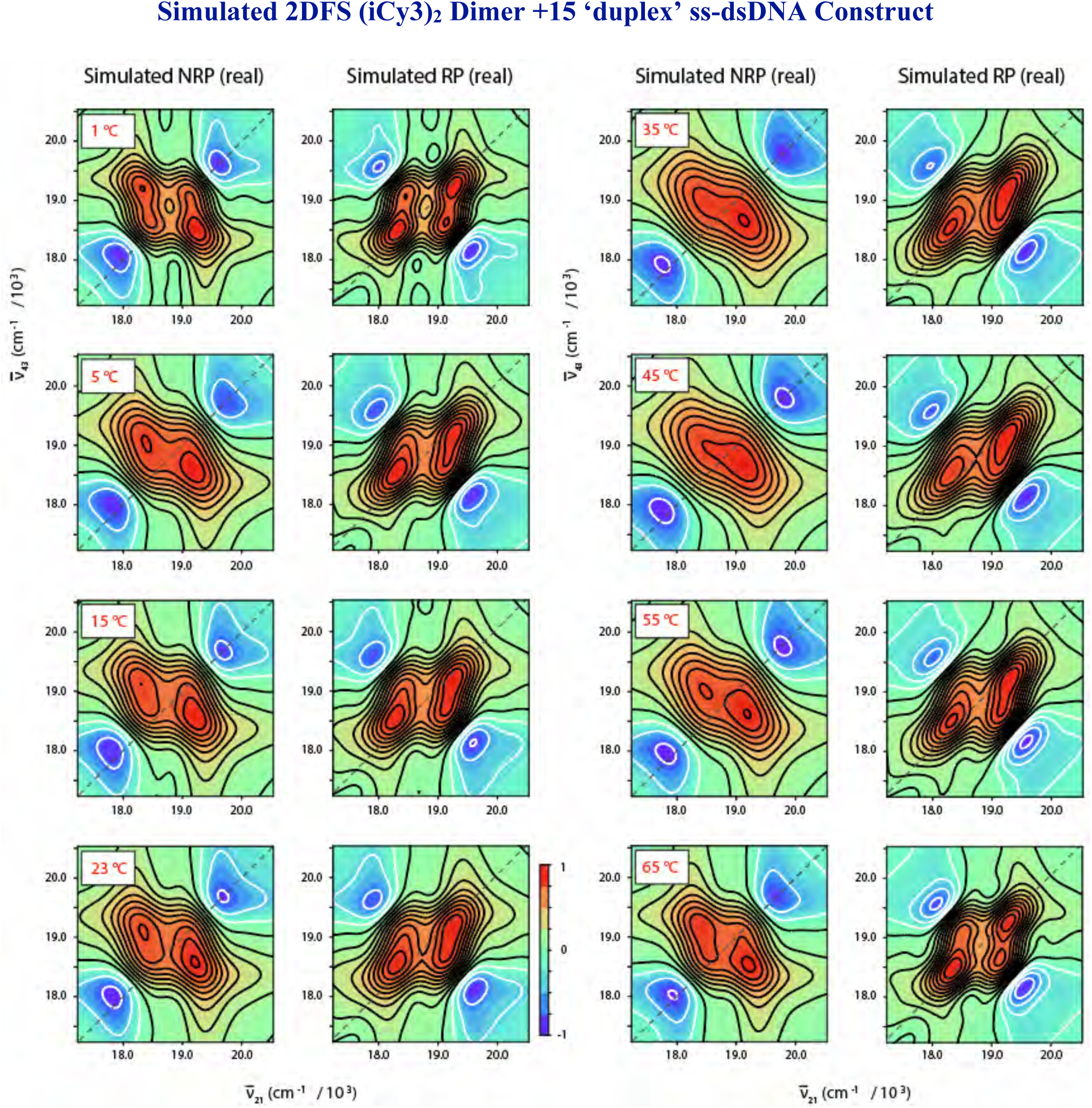
Temperature-dependent simulated NRP and RP 2DFS data (real part) for the (iCy3)_2_ dimer labeled +15 ‘duplex’ ss-dsDNA construct. Simulations were performed using the dimer Hamiltonian given by Eq. (4) of the main text. The dimer Hamiltonian parameters used as input, in addition to the optimized homogeneous and inhomogeneous spectral line widths determined from our analyses, are listed in Table 2 of the main text.

**Figure S8.**
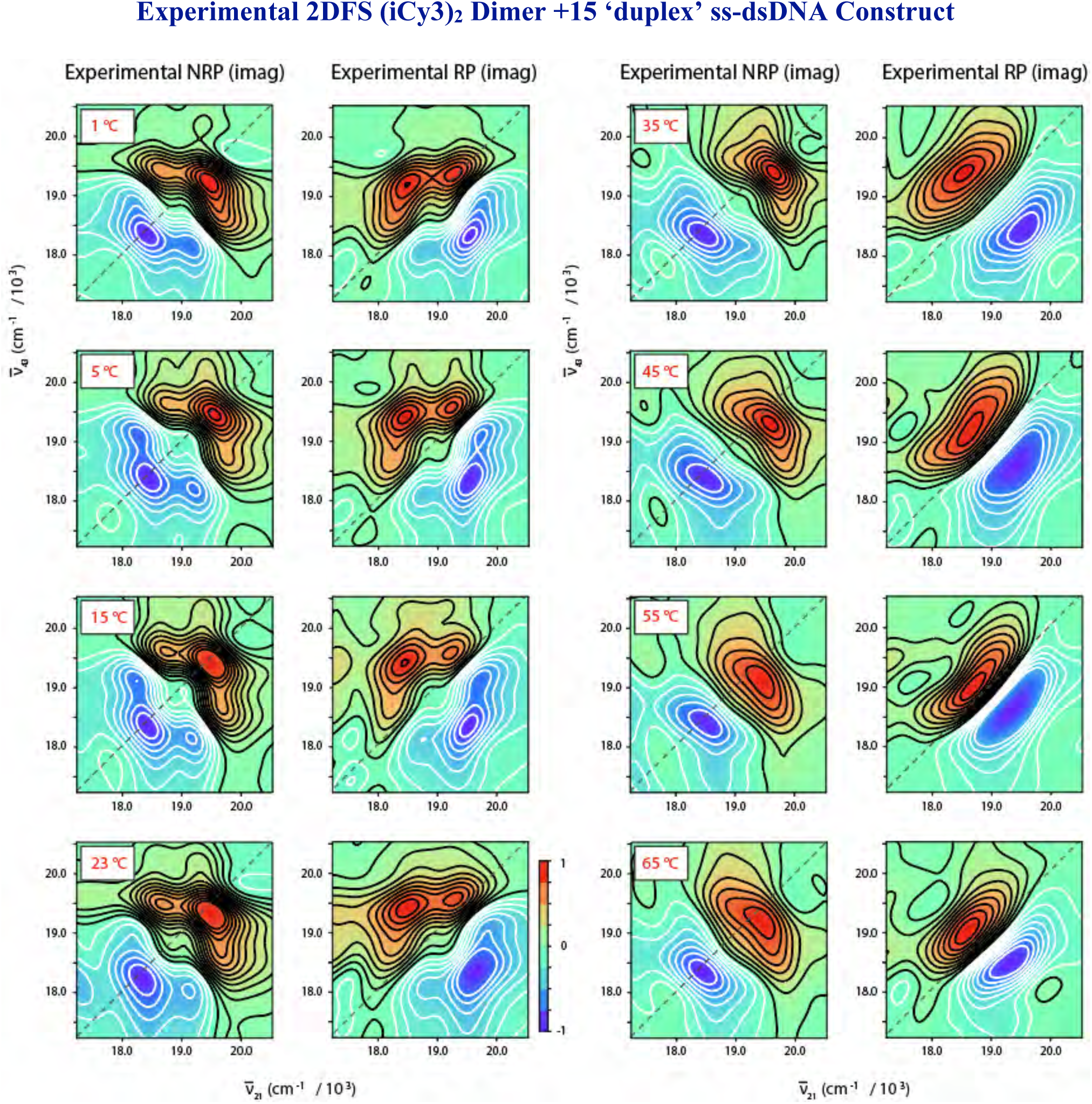
Temperature-dependent experimental NRP and RP 2DFS data (imaginary part) for the (iCy3)_2_ dimer labeled +15 ‘duplex’ ss-dsDNA construct. The laser spectrum used for these measurements is shown in Fig. 4 of the main text.

**Figure S9.**
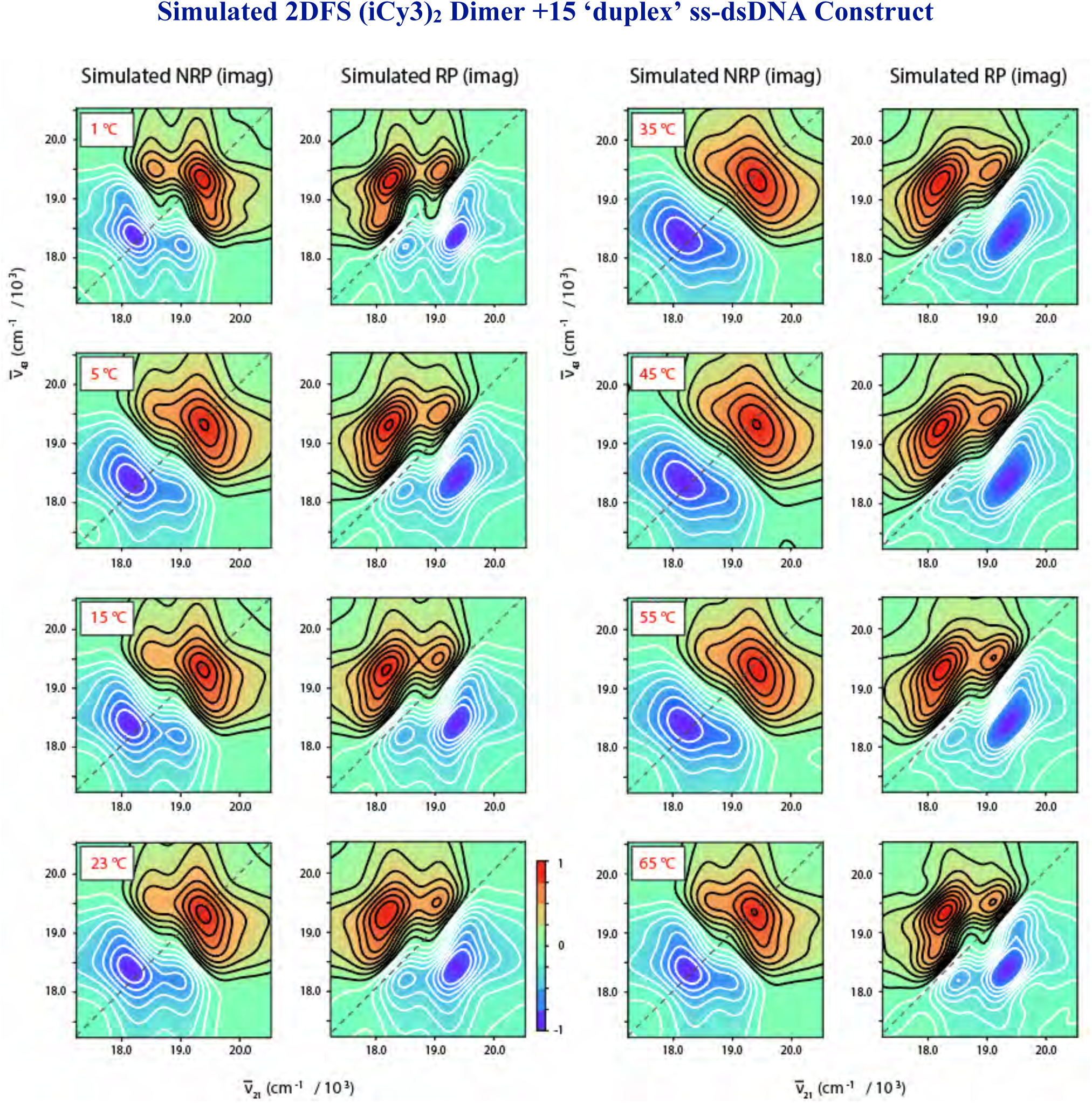
Temperature-dependent simulated NRP and RP 2DFS data (imaginary part) for the (iCy3)_2_ dimer labeled +15 ‘duplex’ ss-dsDNA construct. Simulations were performed using the dimer Hamiltonian given by Eq. (4) of the main text. The dimer Hamiltonian parameters used as input, in addition to the optimized homogeneous and inhomogeneous spectral line widths determined from our analyses, are listed in Table 2 of the main text.

**Figure S10.**
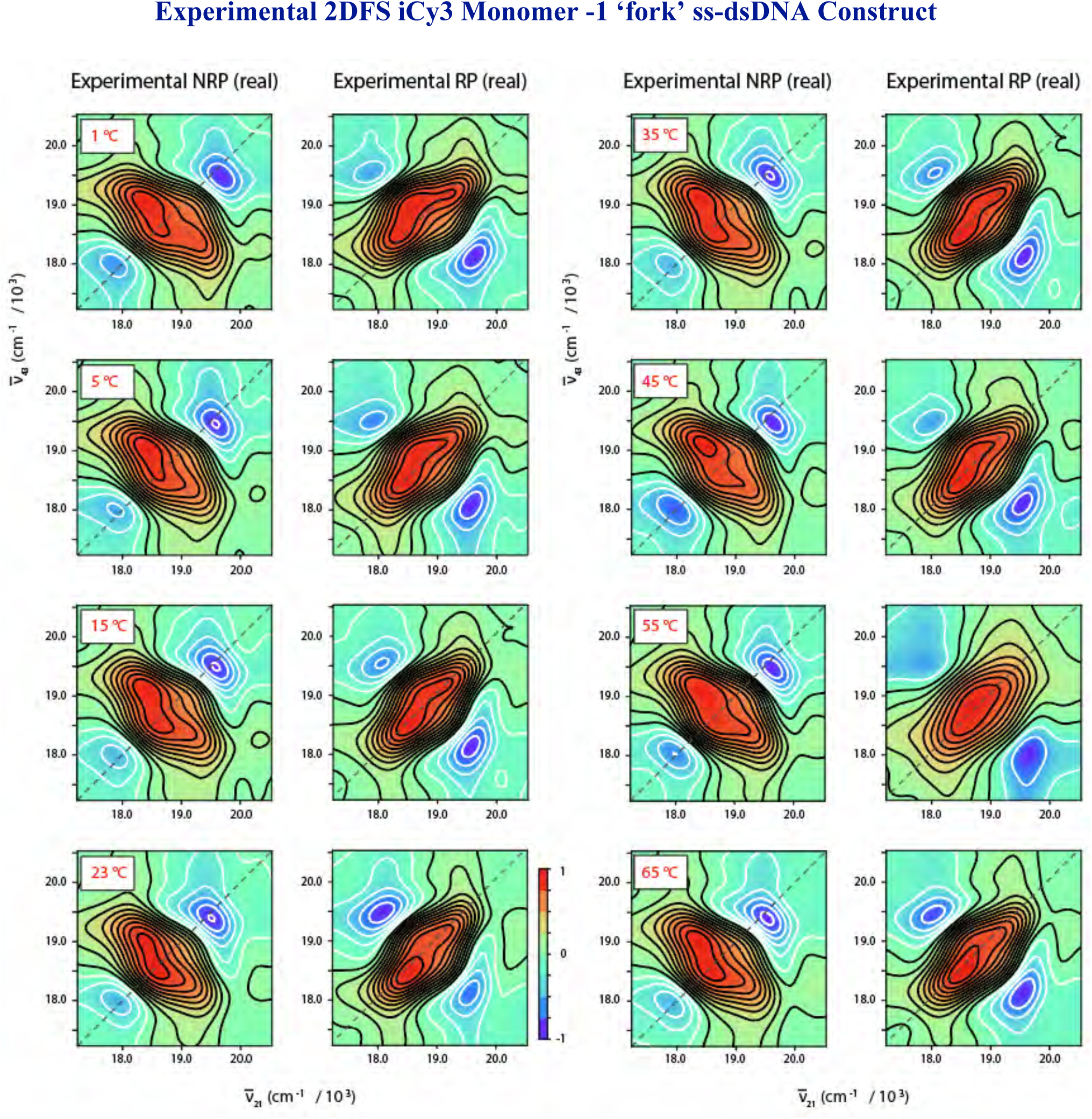
Temperature-dependent experimental NRP and RP 2DFS data (real part) for the iCy3 monomer labeled −1 ‘fork’ ss-dsDNA construct. The laser spectrum used for these measurements is shown in Fig. 4 of the main text. The melting temperature of the duplex region of the construct is ∼65 °C.

**Figure S11.**
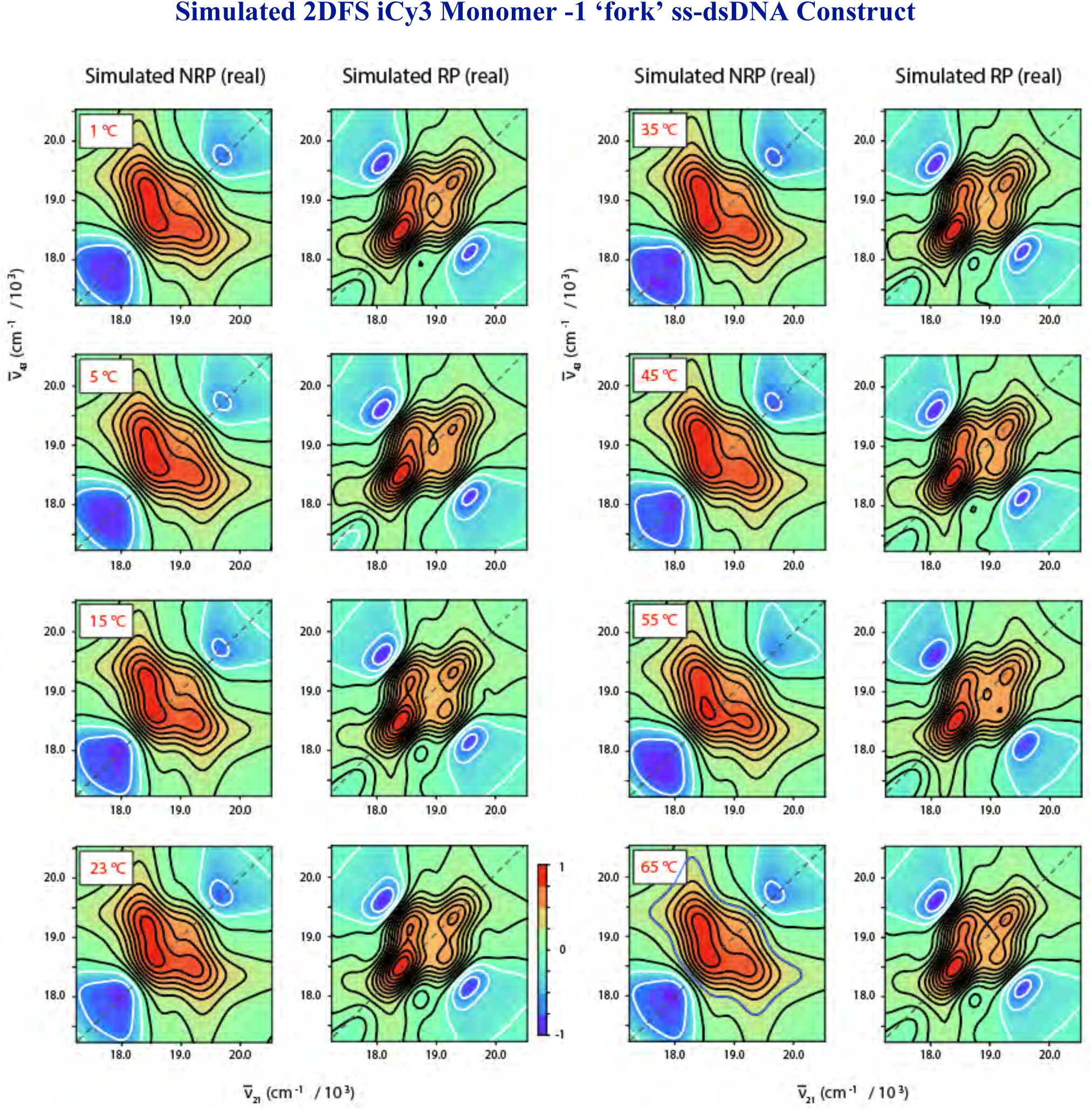
Temperature-dependent simulated NRP and RP 2DFS data (real part) for the iCy3 monomer labeled −1 ‘fork’ ss-dsDNA construct. Simulations were performed using the monomer Hamiltonian given by Eq. (1) of the main text. The monomer Hamiltonian parameters used as input, in addition to the optimized homogeneous and inhomogeneous spectral line widths determined from our analyses, are listed in Table S2.

**Figure S12.**
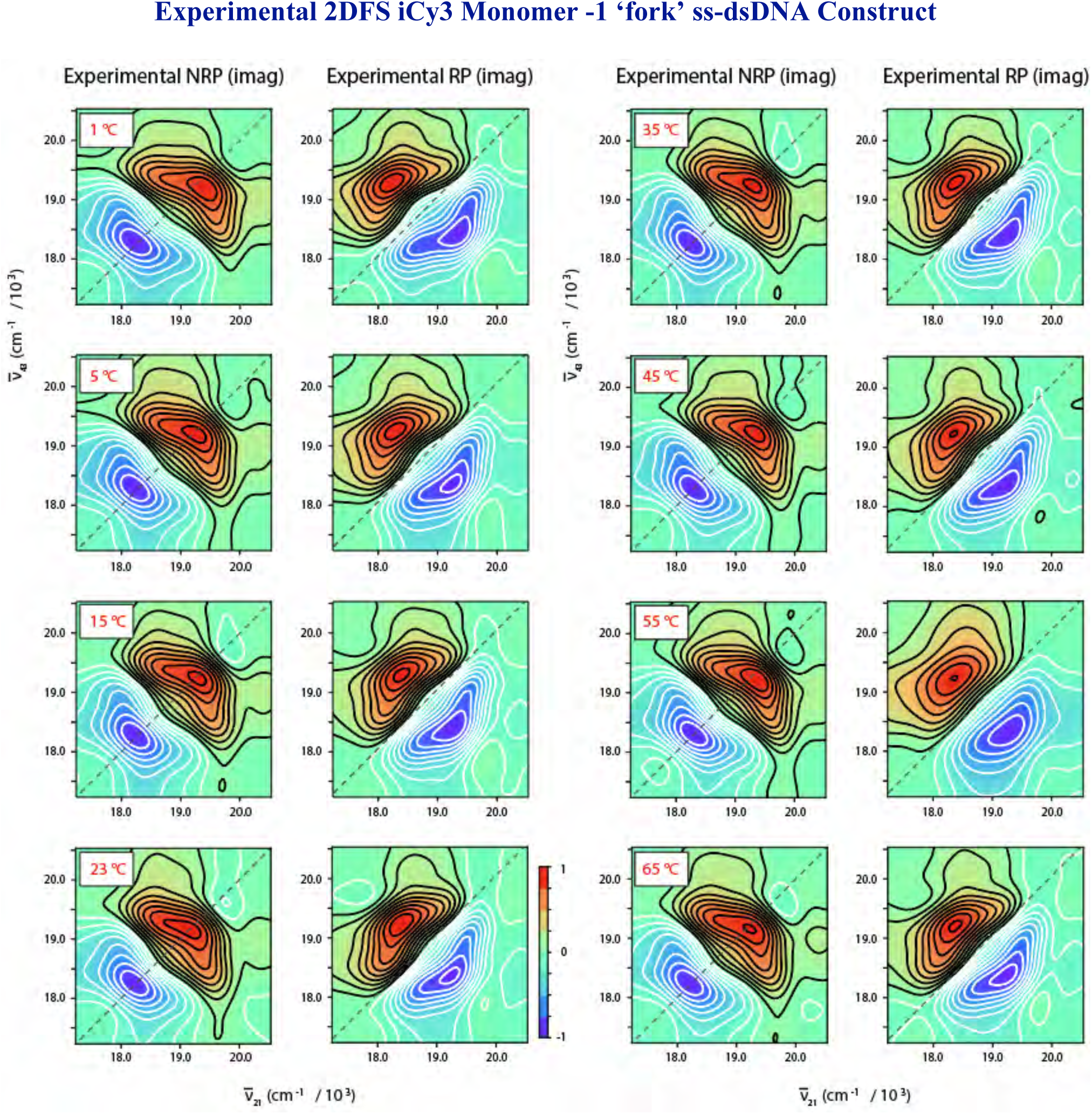
Temperature-dependent experimental NRP and RP 2DFS data (imaginary part) for the iCy3 monomer labeled −1 ‘fork’ ss-dsDNA construct. The laser spectrum used for these measurements is shown in Fig. 4 of the main text.

**Figure S13.**
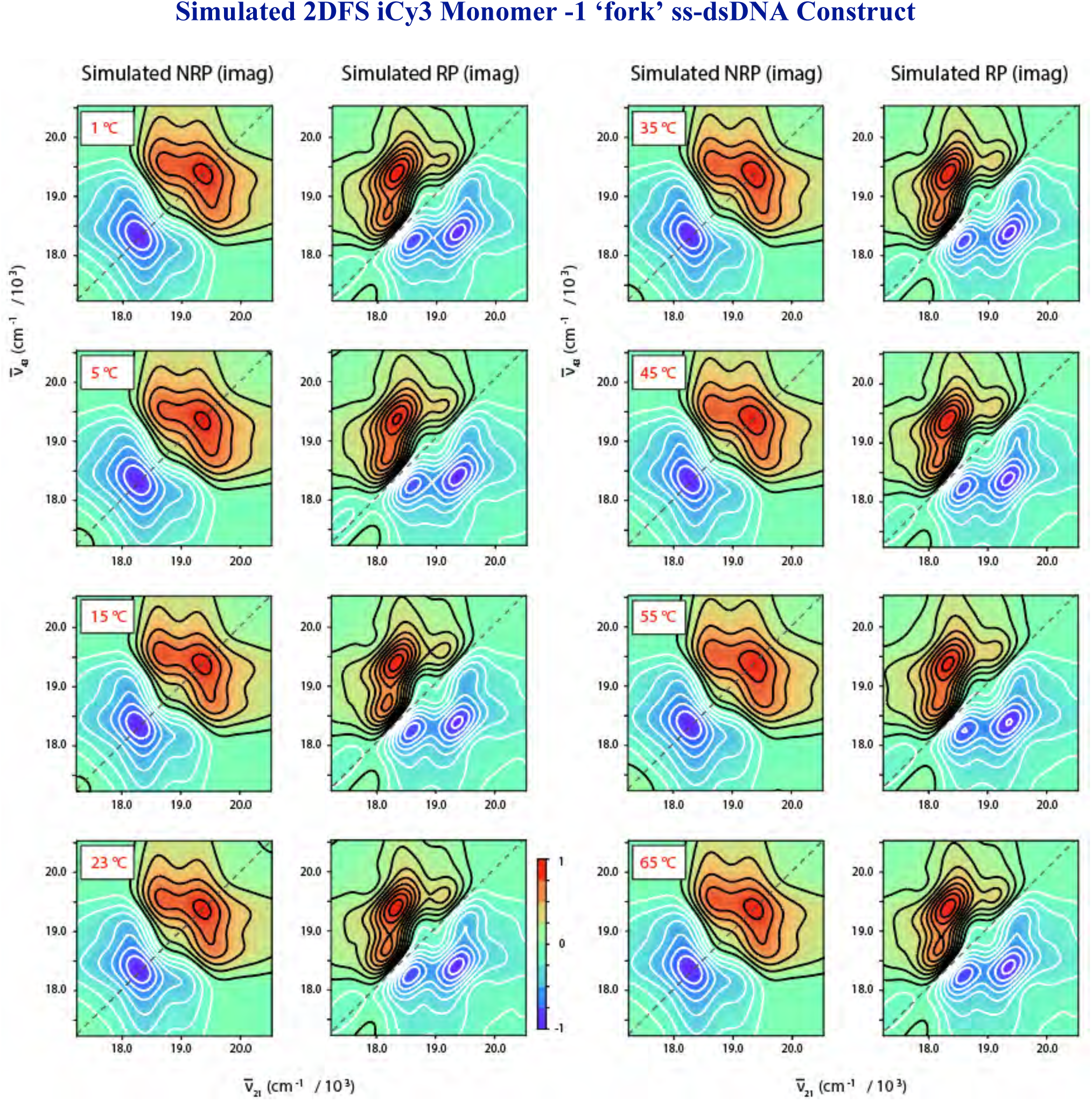
Temperature-dependent simulated NRP and RP 2DFS data (imaginary part) for the iCy3 monomer labeled −1 ‘fork’ ss-dsDNA construct. Simulations were performed using the monomer Hamiltonian given by Eq. (1) of the main text. The monomer Hamiltonian parameters used as input, in addition to the optimized homogeneous and inhomogeneous spectral line widths determined from our analyses, are listed in Table S2.

**Figure S14.**
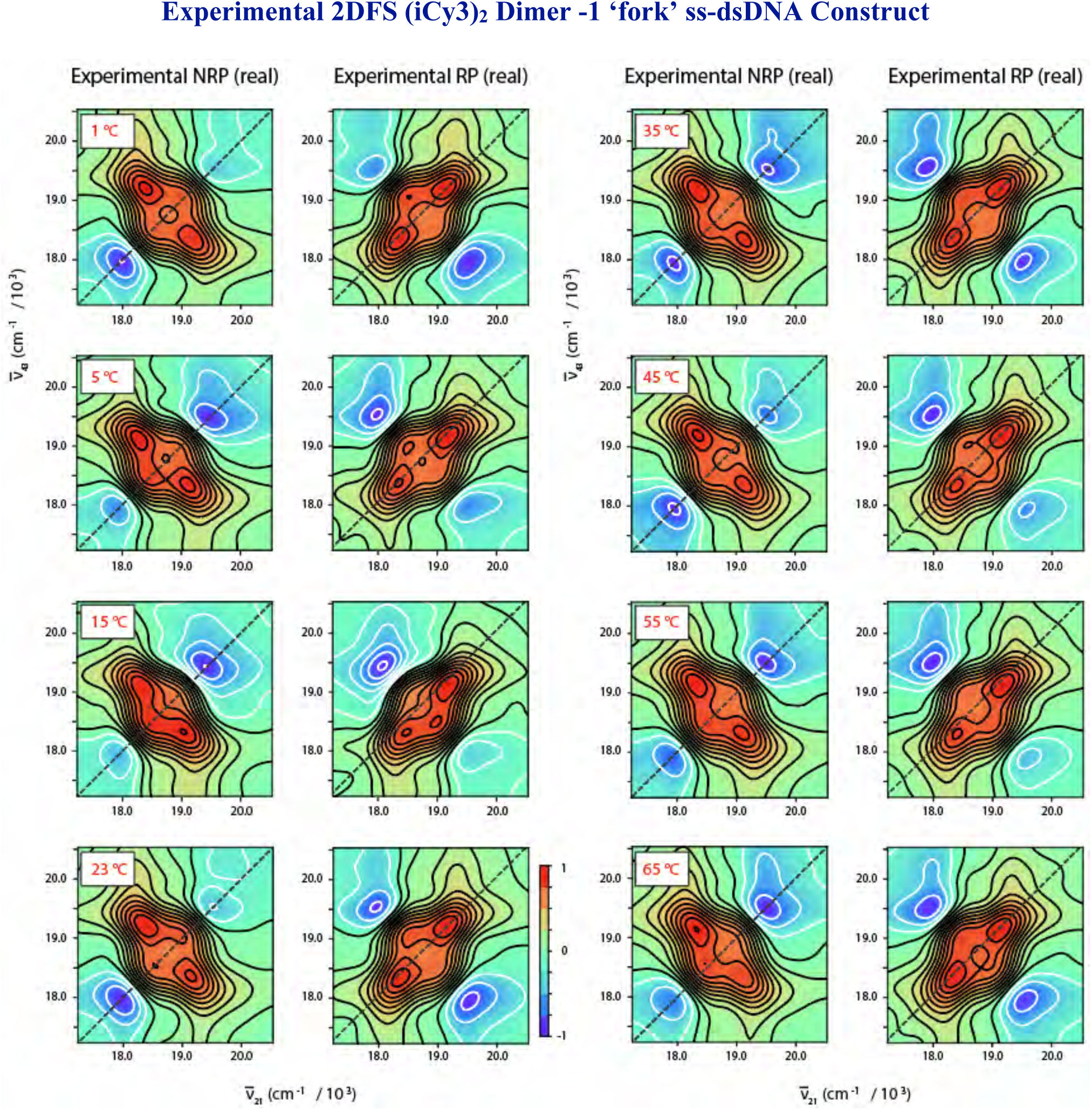
Temperature-dependent experimental NRP and RP 2DFS data (real part) for the (iCy3)_2_ dimer labeled −1 ‘fork’ ss-dsDNA construct. The laser spectrum used for these measurements is shown in Fig. 4 of the main text. The melting temperature of the duplex region of the construct is ∼65 °C.

**Figure S15.**
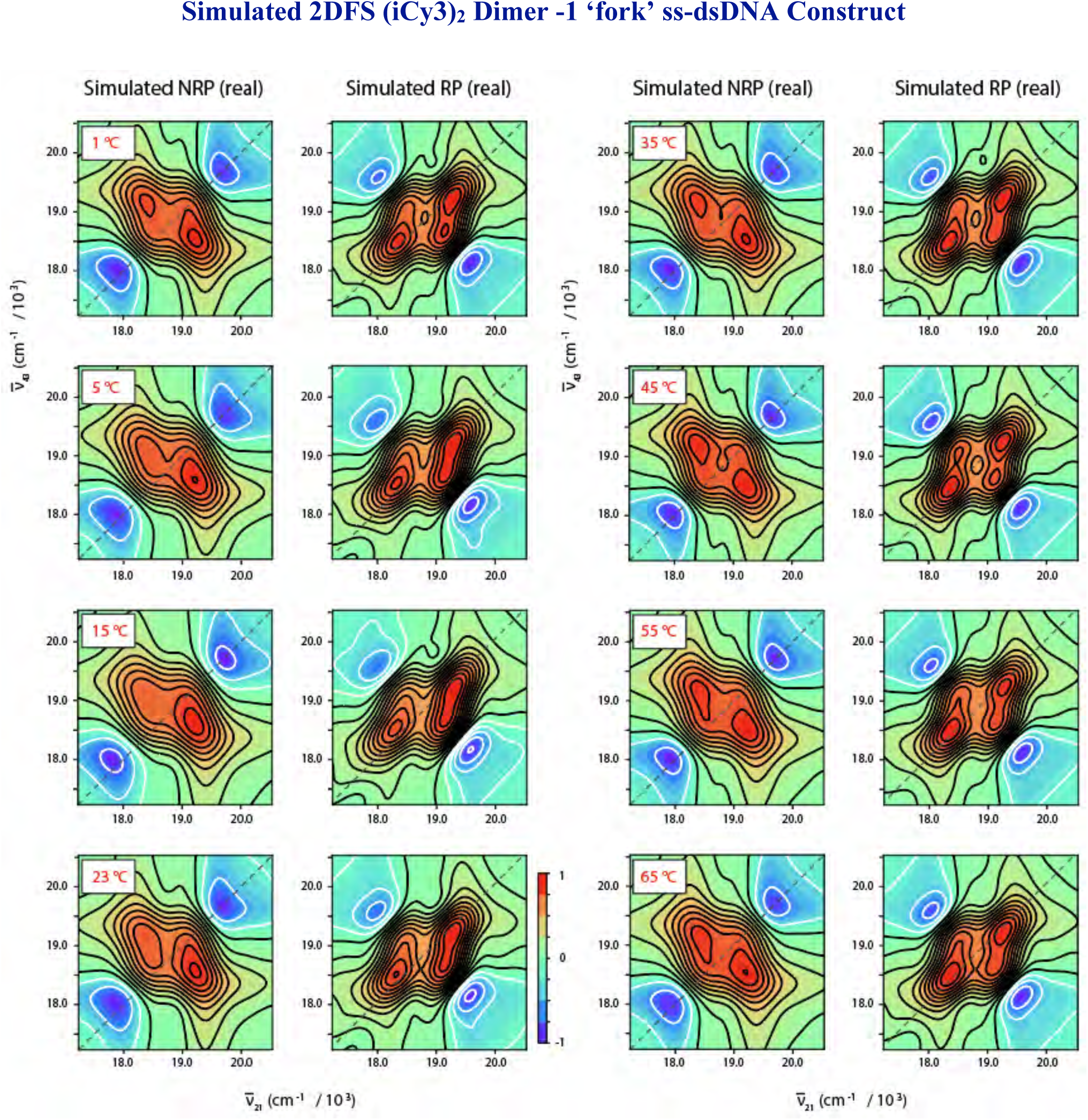
Temperature-dependent simulated NRP and RP 2DFS data (real part) for the (iCy3)_2_ dimer labeled −1 ‘fork’ ss-dsDNA construct. Simulations were performed using the dimer Hamiltonian given by Eq. (4) of the main text. The dimer Hamiltonian parameters used as input, in addition to the optimized homogeneous and inhomogeneous spectral line widths determined from our analyses, are listed in Table 3 of the main text.

**Figure S16.**
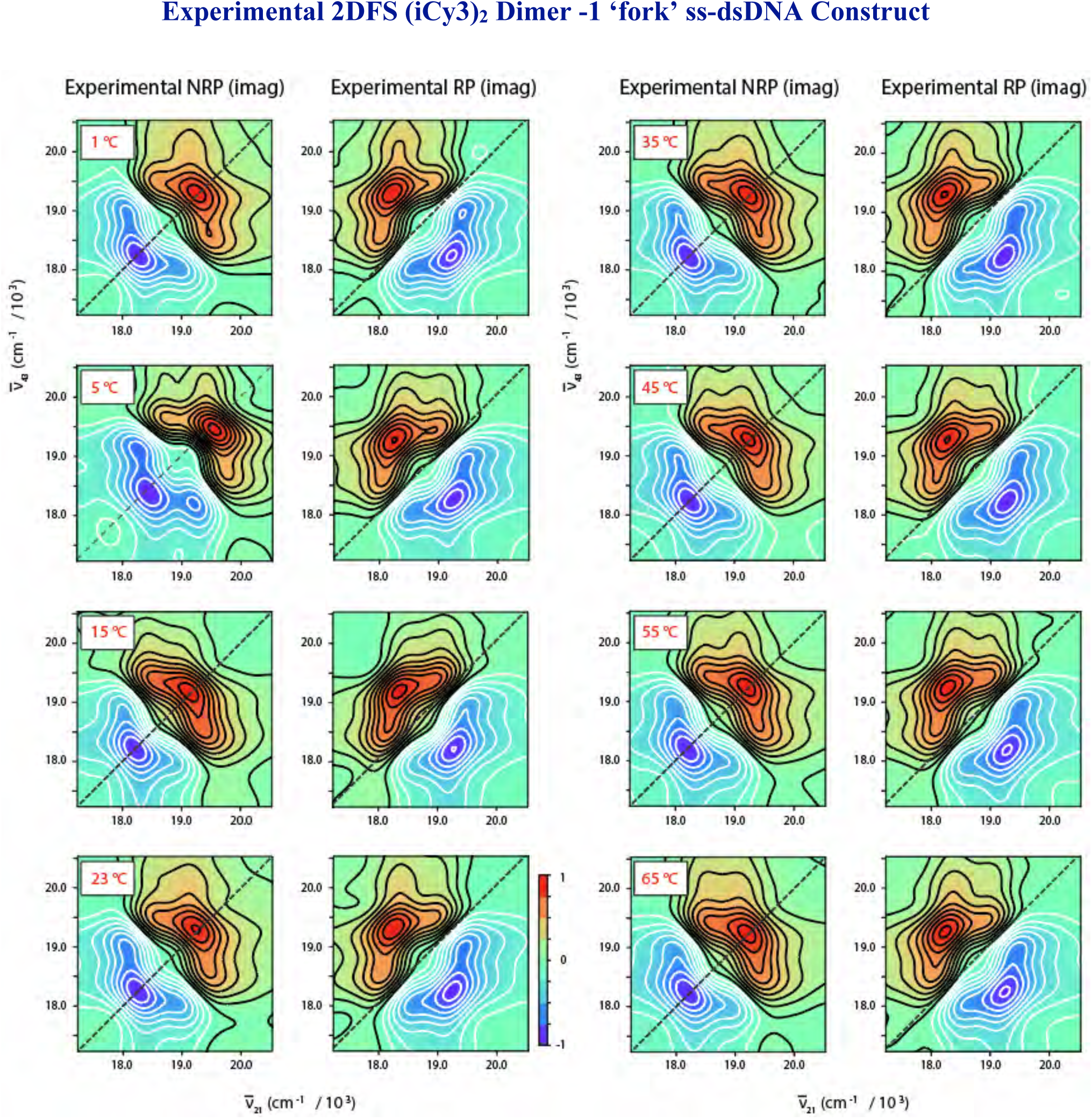
Temperature-dependent experimental NRP and RP 2DFS data (imaginary part) for the (iCy3)_2_ dimer labeled −1 ‘fork’ ss-dsDNA construct. The laser spectrum used for these measurements is shown in Fig. 4 of the main text.

**Figure S17.**
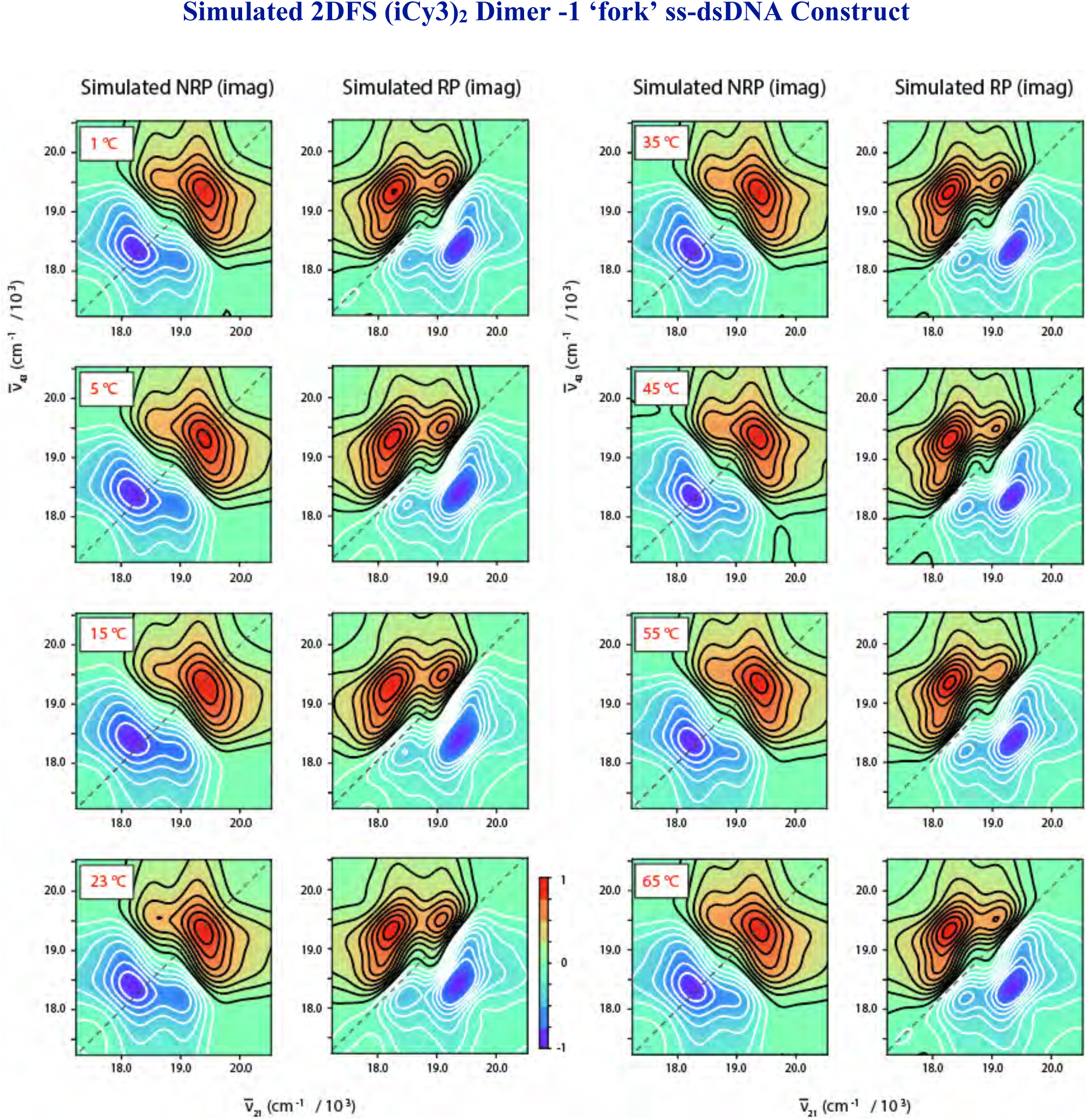
Temperature-dependent simulated NRP and RP 2DFS data (imaginary part) for the (iCy3)_2_ dimer labeled −1 ‘fork’ ss-dsDNA construct. Simulations were performed using the dimer Hamiltonian given by Eq. (4) of the main text. The dimer Hamiltonian parameters used as input, in addition to the optimized homogeneous and inhomogeneous spectral line widths determined from our analyses, are listed in Table 3 of the main text.

**Figure S18.**
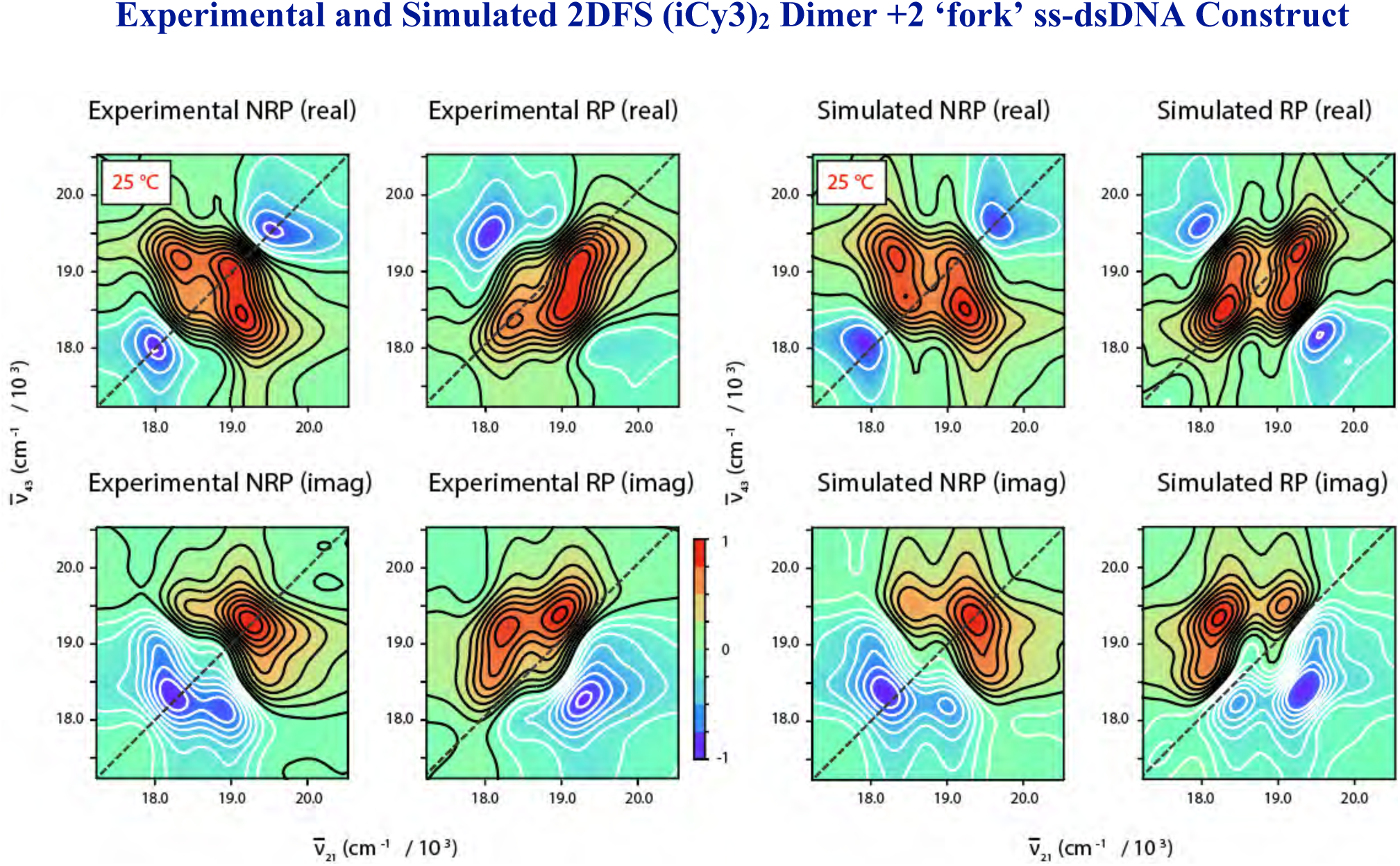
Room temperature (25 °C) experimental and simulated 2DFS data (RP, NRP, real, imaginary) for the (iCy3)_2_ dimer labeled +2 ‘fork’ ss-dsDNA construct. The laser spectrum used for the measurements is shown in Fig. 4 of the main text. Simulations were performed using the dimer Hamiltonian given by Eq. (4) of the main text. The dimer Hamiltonian parameters used as input, in addition to the optimized homogeneous and inhomogeneous spectral line widths determined from our analyses, are listed in Table 4 of the main text.

**Figure S19.**
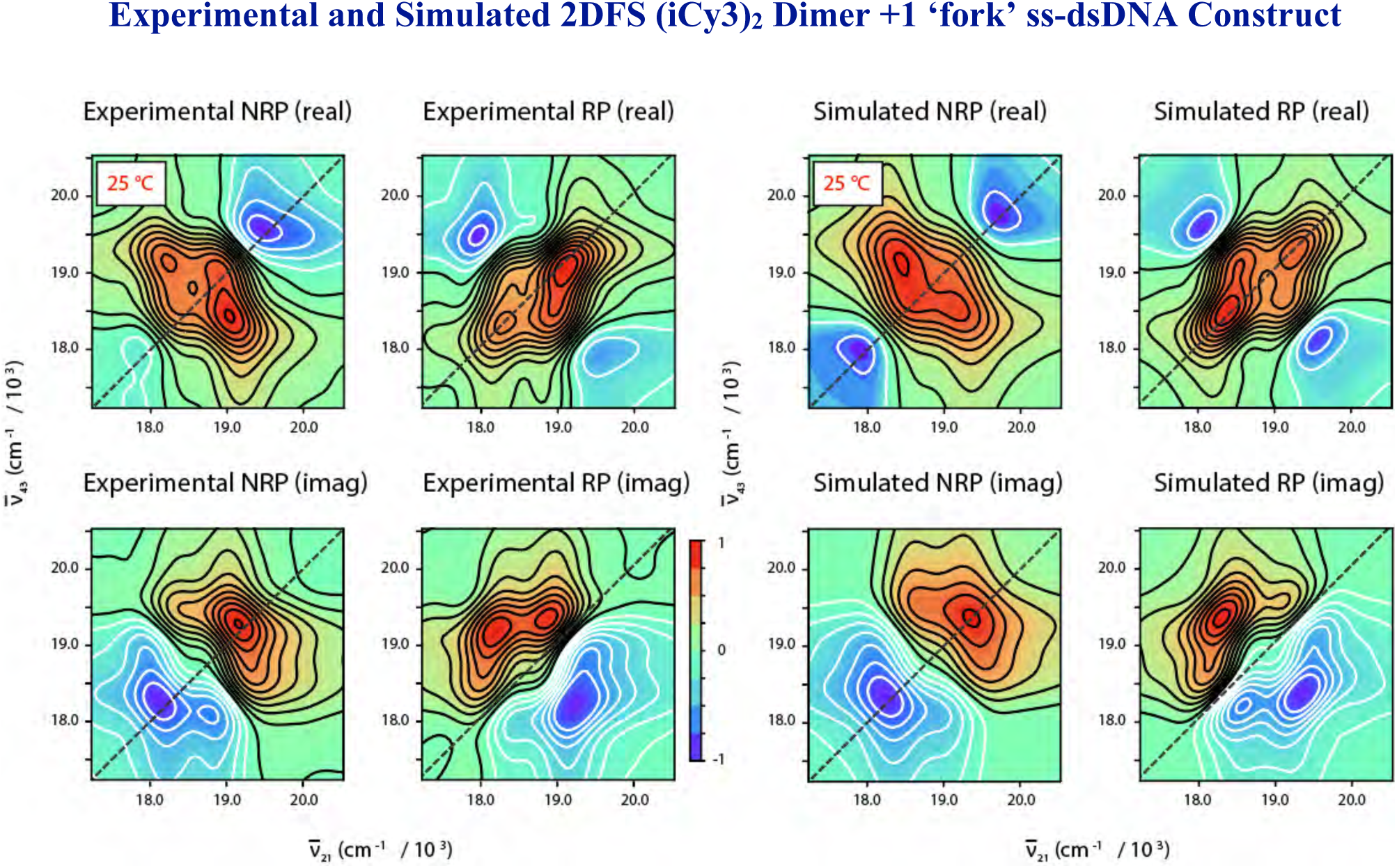
Room temperature (25 °C) experimental and simulated 2DFS data (RP, NRP, real, imaginary) for the (iCy3)_2_ dimer labeled +1 ‘fork’ ss-dsDNA construct. The laser spectrum used for the measurements is shown in Fig. 4 of the main text. Simulations were performed using the dimer Hamiltonian given by Eq. (4) of the main text. The dimer Hamiltonian parameters used as input, in addition to the optimized homogeneous and inhomogeneous spectral line widths determined from our analyses, are listed in Table 4 of the main text.

**Figure S20.**
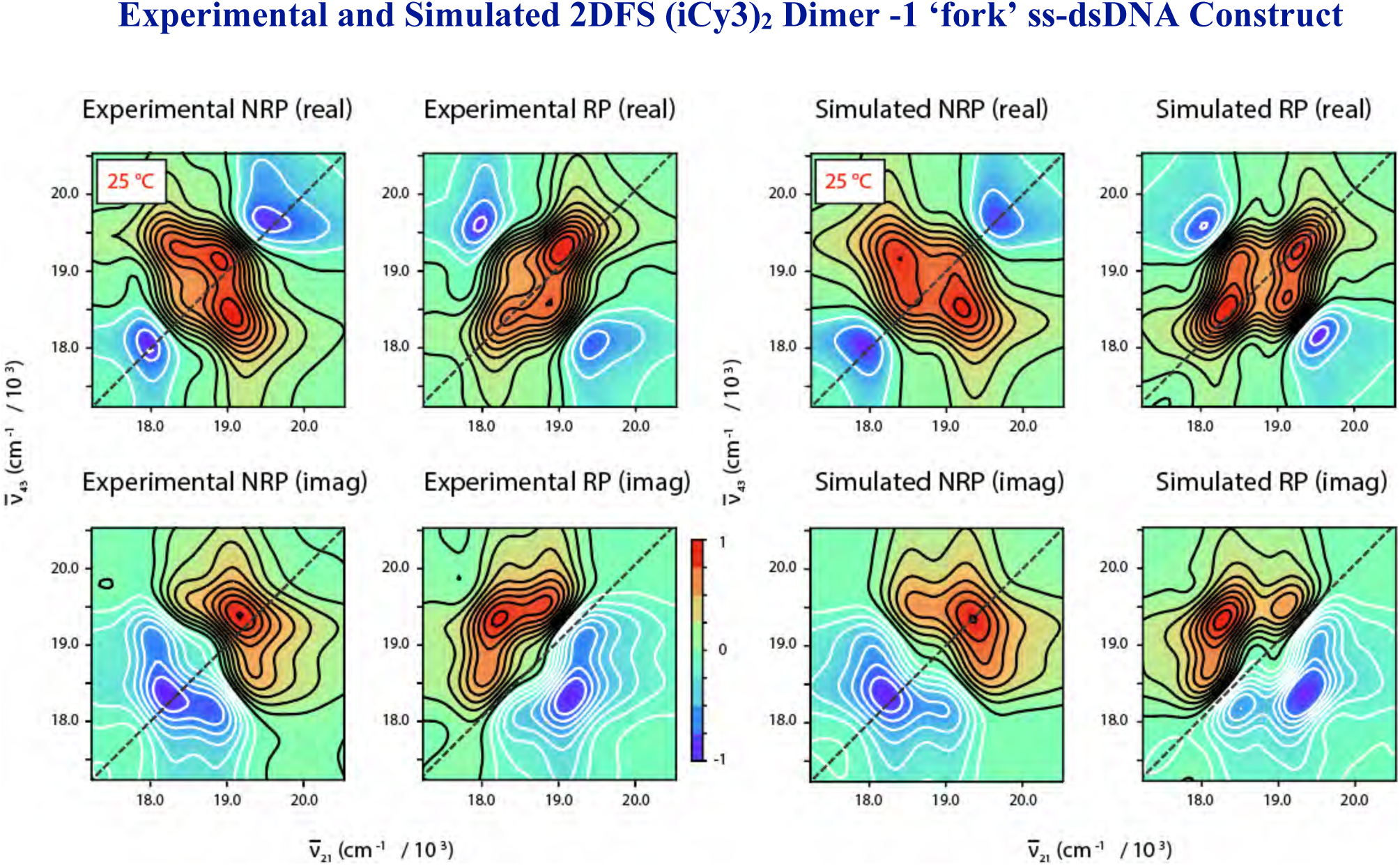
Room temperature (25 °C) experimental and simulated 2DFS data (RP, NRP, real, imaginary) for the (iCy3)_2_ dimer labeled −1 ‘fork’ ss-dsDNA construct. The laser spectrum used for the measurements is shown in Fig. 4 of the main text. Simulations were performed using the dimer Hamiltonian given by Eq. (4) of the main text. The dimer Hamiltonian parameters used as input, in addition to the optimized homogeneous and inhomogeneous spectral line widths determined from our analyses, are listed in Table 4 of the main text.

**Figure S21.**
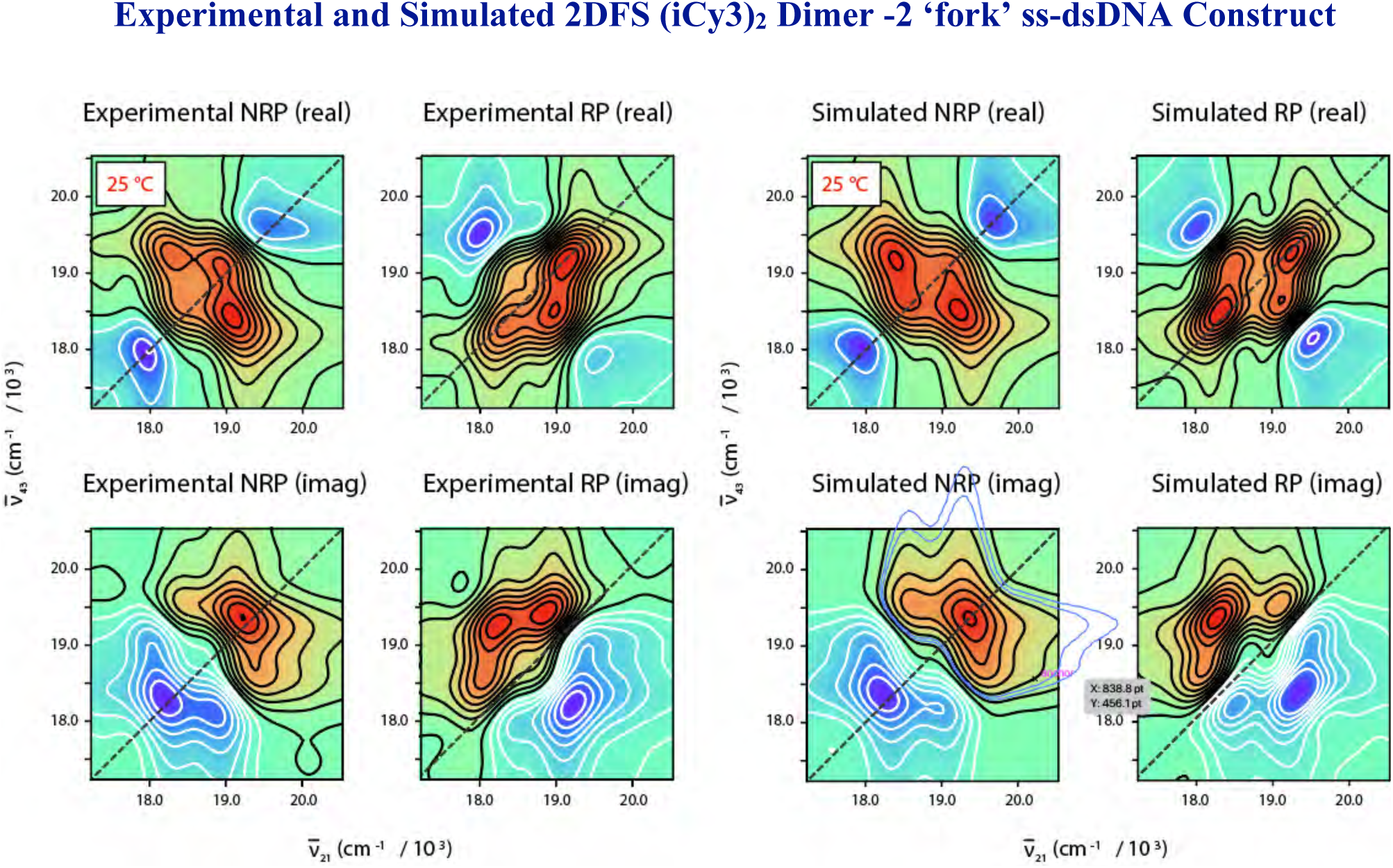
Room temperature (25 °C) experimental and simulated 2DFS data (RP, NRP, real, imaginary) for the (iCy3)_2_ dimer labeled −2 ‘fork’ ss-dsDNA construct. The laser spectrum used for the measurements is shown in Fig. 4 of the main text. Simulations were performed using the dimer Hamiltonian given by Eq. (4) of the main text. The dimer Hamiltonian parameters used as input, in addition to the optimized homogeneous and inhomogeneous spectral line widths determined from our analyses, are listed in Table 4 of the main text.

**Figure S22.**
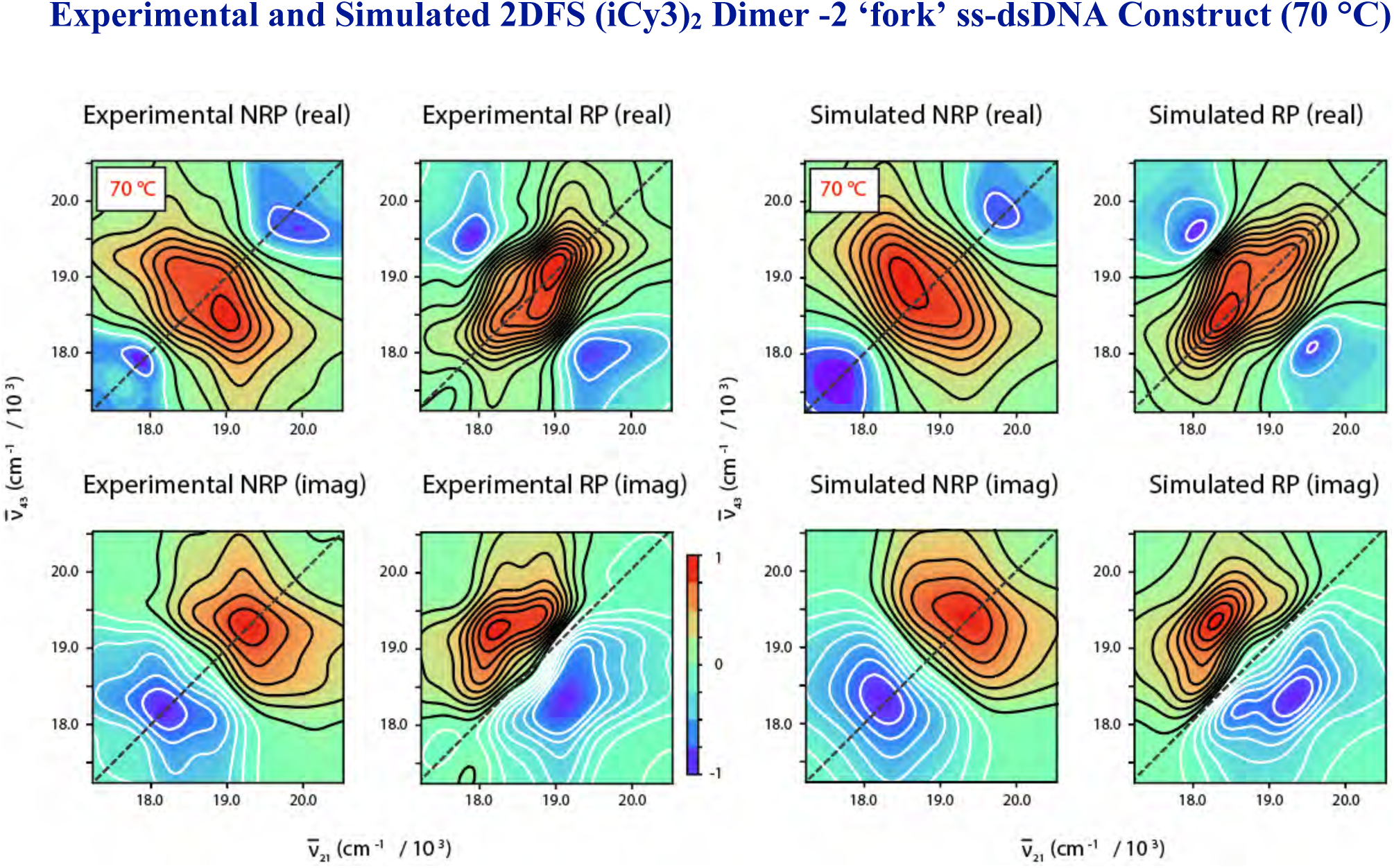
High temperature (70 °C) experimental and simulated 2DFS data (RP, NRP, real, imaginary) for the (iCy3)_2_ dimer labeled −2 ‘fork’ ss-dsDNA construct. The laser spectrum used for the measurements is shown in Fig. 4 of the main text. Simulations were performed using the monomer Hamiltonian given by Eq. (1) of the main text. The monomer Hamiltonian parameters used as input, in addition to the optimized homogeneous and inhomogeneous spectral line widths determined from our analyses, are listed in Table S1.

**Figure S23.**
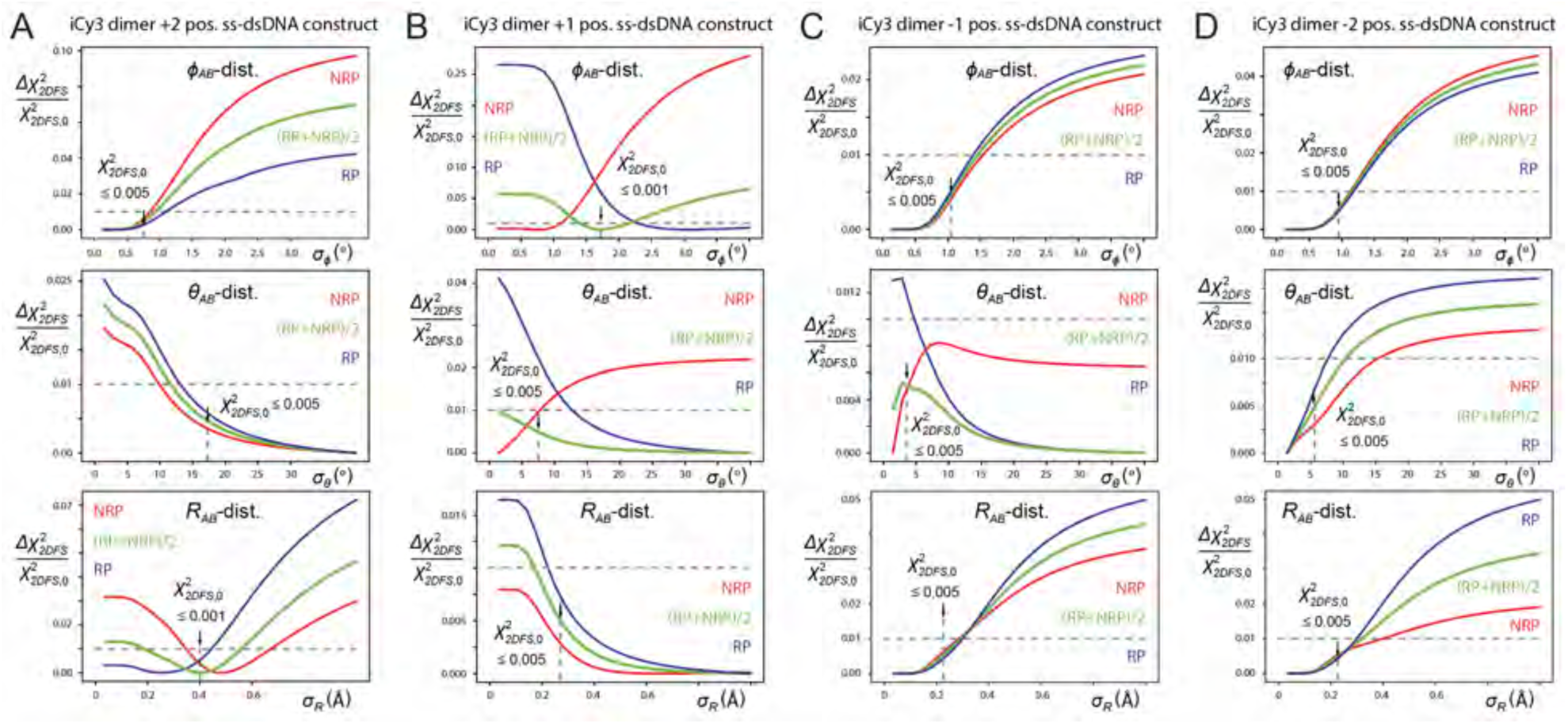
Cross-sections of the relative deviation of the linear least squares error functions, 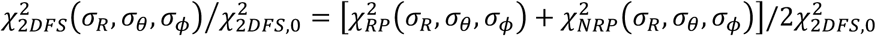, versus the standard deviations of the conformational coordinates *σ*_*ϕ*_ (top), *σ*_*θ*_(middle) and *σ*_*R*_(bottom) for the (iCy3)_2_ dimer ss-dsDNA constructs at probe labeling sites (***A***) +2, (***B***) +1, (***C***) −1 and (***D***) −2. The error functions cross-sections are plotted relative to their ‘optimized’ values, 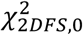 (indicated by vertical arrows). Optimized values of the standard deviation are defined as the 0.5% threshold for cases in which the error function approached its minimum asymptotically (as do the *σ*_*ϕ*_ and *σ*_*θ*_ cross-sections), and the 0.1% threshold for cases in which the error function exhibited a distinct minimum (as shown for the *σ*_*R*_ cross-section). ‘Confidence intervals’ are defined as a 1% deviation of the error function from its optimized value (indicated by horizontal dashed lines).

